# Novel VHH targeting a unique TfR1 epitope for efficient cross-species delivery of drugs in the CNS

**DOI:** 10.1101/2025.05.23.649291

**Authors:** Marion David, Romy Cohen, Diane Beuzelin, Géraldine Ferracci, Marina Cresci, Aude Faucon, Karima Abouzid, Gersande Godefroy, Maxime Masse, Yasmine Mechioukhi, Stéphane D. Girard, Ingrid Julien-Gau, Cléa Boursery, Christophe Fraisier, Karine Varini, Morgane Thomas, Guillaume Jacquot, Pascaline Lecorche, Bastien Serrano, Andréa Romette, Gauthier Dangla Pélissier, Magali Godard, Vincent Saillant, Tiphanie Pruvost, Raphael Sierocki, Patrick Chames, Brigitte Kerfelec, Jamal Temsamani, Michel Khrestchastisky

**Author notes:** **Corresponding authors:** Dr. Michel Khrestchatisky, Institut de Neurophysiopathologie (INP), UMR 7051, Faculté de Médecine, 27 Bd Jean Moulin, 13005 Marseille, France., Marion David, Vect-Horus, Faculté de Médecine, 27 Bd Jean Moulin, 13005 Marseille, France. Authors contributed equally to the manuscript (Marion David, Romy Cohen).

## Abstract

The blood-brain barrier (BBB) is a major obstacle for delivering therapeutic agents to the central nervous system (CNS), posing significant challenges for treating neurological disorders. Among current strategies to improve brain drug exposure, hijacking physiological pathways involved in receptor-mediated transcytosis (RMT) has emerged as a promising strategy. While targeting transferrin receptor 1 (TfR1) is widely explored, many TfR1-antibodies lack cross-species reactivity, limiting translational development. In the present study, we identified and characterized camelid-derived single-domain antibodies (VHHs) with robust cross-reactivity to rodent, rhesus monkey, and human TfR1. Epitope mapping of the VHH revealed a novel binding site at the interface of the TfR1 dimer. When fused to a human IgG1 Fc domain, these VHHs, as monomers or homodimers, were efficiently internalized by engineered CHO cells and brain endothelial cells expressing TfR1 from different species. Systemic administration of VHH-Fc constructs in mice demonstrated significantly improved brain uptake compared to irrelevant controls. Functional delivery was confirmed using neurotensin (NT)-induced hypothermia, and we established correlations between *in vivo* effects and TfR1 VHH binding properties determined by surface plasmon resonance. Notably, efficient BBB transcytosis was associated with intermediate affinity and rapid dissociation rates. Engineered variants maintained favorable cross-species binding, including similar affinities to human and non-human primate TfR1, facilitating translational studies. The cross-reactive anti-TfR1 VHHs we developed offer a versatile and modular platform for CNS drug delivery and hold promise as molecular shuttles for transporting therapeutic agents across the BBB. Our work establishes a robust foundation for developing next-generation brain-targeted biotherapeutics, including peptides, enzyme replacement therapies, antibody-based treatments for neurodegenerative diseases, and oligonucleotide delivery for CNS disorders, enabling seamless translation from preclinical to clinical applications.

## Introduction

The blood-brain barrier (BBB) controls and restricts the passage of molecules from blood to the nervous tissue parenchyma. This barrier protects very efficiently the central nervous system (CNS) from endogenous (plasma proteins, immune cells…) or exogenous (viruses, toxins, xenobiotics…) potentially neurotoxic agents. This interface also prevents entry in the brain of the vast majority of therapeutic agents. To cross the BBB in pharmacologically significant amounts, molecules should be less than 500 Da in molecular weight, highly lipophilic (Pardridge 2005), and should not be substrate of BBB efflux pumps (Begley 2004). For these reasons, most CNS therapeutics currently on the market are based on small organic molecules that treat mainly affective disorders, chronic pain, epilepsy and insomnia (Ajay, Bemis et al. 1999, Ghose, Viswanadhan et al. 1999). However, new generations of drug candidates, with enhanced efficacy and specificity, such as peptides, oligonucleotides, growth factors, cytokines, enzymes or therapeutic antibodies, are under development for the treatment of other severe CNS diseases, including neurodegenerative and lysosomal storage diseases.

Considering that the number of individuals with neurodegenerative diseases increases with aging of the population, and that the number of individuals over 60 should increase by 50 % by 2050 (WHO 2022), the pharmaceutical industry is facing a real challenge in developing strategies to efficiently transport therapeutic agents across the BBB (reviewed in (Pardridge 2003, Vlieghe and Khrestchatisky 2010, Vlieghe and Khrestchatisky 2013, Pardridge 2020)). Among the strategies evaluated to deliver protein therapeutics into the brain, the “Trojan horse” approach that hijacks the cellular machinery present at the BBB appears as the safest and most effective (Pardridge, Boado et al. 1992, Jones and Shusta 2007, Fang, Zou et al. 2017). The transport of molecules across the BBB can be facilitated by receptor-mediated transcytosis (RMT). This physiological process involves binding of a ligand to its receptor expressed by brain microvessel endothelial cells (BMEC), internalization by endocytosis, receptor release and transcytosis to the abluminal side of the BBB (Tuma and Hubbard 2003, Villasenor, Schilling et al. 2017, Ayloo and Gu 2019, Haqqani, Bélanger et al. 2024). One of the most studied RMT receptors is transferrin receptor 1 (TfR1). Structurally, this receptor consists of a homodimer of 170 kDa, where each monomer of the ectodomain bears three distinct domains: i) a protease-like domain, ii) a helical domain involved in dimer contacts and in the binding of holo transferrin (holo-Tf), the iron transporter (Cheng, Zak et al. 2004), and iii) an apical domain involved in the binding of another ligand, ferritin heavy chain (H-Ft), an iron storage and transporter molecule (Li, Fang et al. 2010, Montemiglio, Testi et al. 2019). TfR1 is expressed in brain endothelium (Jefferies, Brandon et al. 1984, Pardridge, Eisenberg et al. 1987), and is involved in iron delivery into the brain (Fishman, Rubin et al. 1987). TfR1 is also abundant in blood and lung cells (Chan and Gerhardt 1992). Tf and H-Ft have been studied as brain shuttles (Chang, Jallouli et al. 2009, Wiley, Webster et al. 2013, Thomsen, Johnsen et al. 2022, Sevieri, Mazzucchelli et al. 2023). However, they compete with endogenous ligands and especially with holo-Tf, whose blood concentration of 0.75 g/L is estimated to occupy 99.99 % of TfR1 at the BBB (Pardridge and Chou 2021). TfR1 ligands that do not compete with Tf, including antibodies, antibody fragments and single domain antibodies from camelids (Variable domain of Heavy chain only antibodies from camelids, VHH) or cartilaginous fish (VNARs), have been studied for brain delivery (Yu, Atwal et al. 2014, Karaoglu Hanzatian, Schwartz et al. 2018, Stocki, Szary et al. 2020, Wouters, Jaspers et al. 2020, Stocki, Szary et al. 2023). These ligands with specificity for rodent TfR1 have the capacity to deliver different types of cargos (peptides, enzymes, antibodies, growth factors, oligonucleotides) to the brain (Pardridge, Buciak et al. 1991, Moos and Morgan 2001, Pardridge 2005, Boado, Zhang et al. 2009, Manich, Cabezón et al. 2013, Pardridge 2015, Hultqvist, Syvanen et al. 2017, Sehlin, Fang et al. 2017, Haqqani, Thom et al. 2018, Wouters, Jaspers et al. 2020). Non-human primate (NHP) and human cross-reactive TfR1 targeting molecules have also been described but developing rodent-primate cross-species reactive ligands remains challenging (Kariolis, Wells et al. 2020, Edavettal, Cejudo-Martin et al. 2022, Rué, Jaspers et al. 2023). The potential of TfR1 in brain delivery is underscored by the clinical development of several TfR1-targeting conjugates, including the approved antibody-iduronate-2-sulfatase fusion for the treatment of neuronopathic mucopolysaccharidosis II (Sonoda, Morimoto et al. 2018). Considering the promising potential of TfR1 for BBB transcytosis and for delivering different therapeutic cargos into the brain, we set up strategies to identify cross-species ligands that bind to TfR1. These ligands are based on VHHs that present the advantage of high specific binding to their epitopes with a molecular weight (MW) approximately 10 times lower (15 kDa) than conventional antibodies (Chames and Baty 2009, Vincke and Muyldermans 2012).

In the present study, we identified novel VHHs that bind mouse, rat, rhesus monkey and human TfR1 with different binding profiles according to species, and that do not compete with the Tf ligand. These VHHs target an original TfR1 epitope localized at the dimer interface of the homodimeric TfR1 that we characterized. We showed that the VHHs fused to a human Immunoglobulin 1 (IgG1) Fc region (hFc) can target *in vitro* engineered cells expressing the TfR1 of different species, as well as mouse, rat and human BMECs. *In vivo*, we showed that these VHHs efficiently transported the hFc into the CNS. Neurotensin (NT) is a 13 amino-acid neuropeptide that induces hypothermia (HT) when administered intracerebrally. However, these effects were not observed if NT is administered peripherally, due to its rapid degradation and poor passage of the BBB (Banks, Wustrow et al. 1995). When fused to NT, the parental VHH elicited significant HT following systemic administration to WT mice. Optimized VHH-NT affinity variants also induced HT when administered to B-hTfR transgenic mice that express the human TfR1 ectodomain. Among these affinity variants, some presented a remarkable human/rhesus monkey cross-reactivity together with TfR1 binding properties compatible with efficient BBB crossing capacity. Our VHHs are expected to enhance the delivery of various therapeutic agents across the BBB into the brain parenchyma of multiple species thereby facilitating the progression from preclinical research to clinical applications.

## Results

### Establishment of stable cell lines expressing mouse, human or rhesus monkey TfR1

The full-length coding regions of mouse, human and rhesus monkey TfR1 were cloned in frame with EGFP in a pEGFP-C1 vector (Supplemental Figure S1A) and plasmids were transfected in CHO cells. We next isolated the CHO cell lines stably expressing the TfR1 constructs from the different species. Immunocytochemistry confirmed co-localization of TfR1-EGFP with fluorescently labeled transferrin (Tf-Alexa647), indicating proper receptor expression at the cell surface (Supplemental Figure S1B). Western blot analysis using anti-TfR1 and anti-GFP antibodies revealed consistent profiles across species, with a major 130 kDa band and a minor higher molecular weight band likely corresponding to non-reduced receptor dimers (Supplemental Figure S1C).

### Selection of VHHs that bind mouse and human TfR1

A young adult male llama was immunized with a subcutaneous injection of CHO-hTfR1-EGFP membrane preparation, followed by a first booster injection with the same membrane preparation, and two additional booster injections with membrane preparations from both CHO-mTfR1-EGFP and CHO-hTfR1-EGFP cell lines. After total RNA isolation from llama peripheral mononuclear cells, RT-PCR amplification and cloning of VHH cDNA, a library of ∼10^8^ clones was obtained. The VHH repertoire was then displayed on the M13 phage pIII protein following infection with a helper phage in *E. coli*. The library was used to perform four cross-selection strategies on CHO cells expressing either mouse or human TfR1, with a depletion step on CHO-TRVb cells and masking of irrelevant epitopes to enrich for cross-reactive binders (Even-Desrumeaux, Nevoltris et al. 2014). To favor the isolation of VHHs that bind both mouse and human TfR1, a total of 4 cross-selection strategies were performed with the library: the same cell line (CHO-mTfR1-EGFP or CHO-hTfR1-EGFP) for the 2 rounds or alternating the cell lines (1^st^ round with CHO-mTfR1-EGFP and 2^nd^ round with CHO-hTfR1-EGFP, and conversely). Following each selection strategy, 186 clones were randomly screened for their ability to bind TfR1 by flow cytometry experiments on CHO-mTfR1-EGFP or CHO-hTfR1-EGFP. Among a total of 744 clones randomly screened, 105 clones bound mouse TfR1 and 188 clones bound human TfR1, and more importantly, 284 clones bound the receptors of both species (Table 1). The 174 clones with the highest signal in flow cytometry (list of antibodies in Supplemental Table S1) on both receptors were sequenced and further grouped as 73 different sequences. Surprisingly, only 2 distinct VHH families were isolated, defined by the same (or very similar) complementary determining region 3 (CDR3) sequence. Four VHHs were retained for further characterization, production in *E. coli* and purification. Based on CDR3 sequence homology (see selected VHH sequences in Supplemental Table S2), we finally selected 3 VHHs namely C5, B8 and H3, that belonged to the same family and that targeted both mouse and human TfR1. In addition, one VHH, B6, belonged to a different family and targeted human TfR1 only.

**Table 1.**
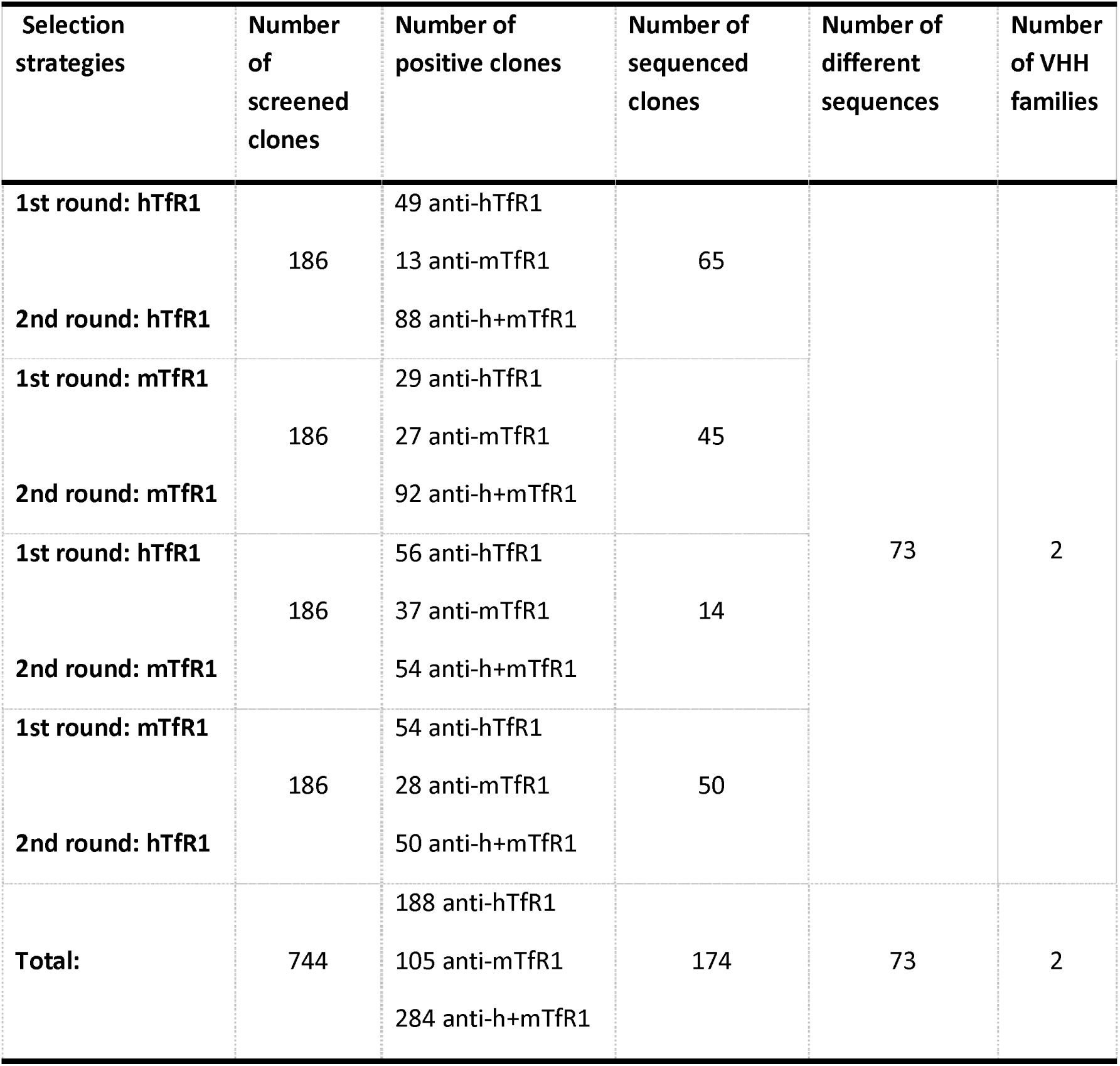
VHH selection and screening results. All selection strategies were performed using CHO cell lines expressing the human and mouse TfR1 and using depletion and mask steps. Clones were screened using flow cytometry and the candidates with the highest signals were sequenced. A VHH family contains VHHs with the same (or very similar) CDR3 sequence.

| Selection strategies | Number of screened clones | Number of positive clones | Number of sequenced clones | Number of different sequences | Number of VHH families |
| --- | --- | --- | --- | --- | --- |
| 1st round: hTfR1 | 186 | 49 anti-hTfR1 | 65 | 73 | 2 |
|  |  | 13 anti-mTfR1 |  |  |  |
| 2nd round: hTfR1 |  | 88 anti-h+mTfR1 |  |  |  |
| 1st round: mTfR1 | 186 | 29 anti-hTfR1 | 45 |  |  |
|  |  | 27 anti-mTfR1 |  |  |  |
| 2nd round: mTfR1 |  | 92 anti-h+mTfR1 |  |  |  |
| 1st round: hTfR1 | 186 | 56 anti-hTfR1 | 14 |  |  |
|  |  | 37 anti-mTfR1 |  |  |  |
| 2nd round: mTfR1 |  | 54 anti-h+mTfR1 |  |  |  |
| 1st round: mTfR1 | 186 | 54 anti-hTfR1 | 50 |  |  |
|  |  | 28 anti-mTfR1 |  |  |  |
| 2nd round: hTfR1 |  | 50 anti-h+mTfR1 |  |  |  |
| Total: | 744 | 188 anti-hTfR1<br>105 anti-mTfR1<br>284 anti-h+mTfR1 | 174 | 73 | 2 |

### Characterization of VHH properties and cross-species reactivity for TfR1

The VHH molecular weights and their theoretical isoelectric points (pI) were calculated (Table 2). As expected, all VHHs had molecular weights of about 15 kDa (with cMyc and 6-His tags), characteristic of VHH domains. The four TfR1-binding VHHs and the irrelevant D12 VHH used as negative control had a slightly acidic pI of approximately 6 with cMyc and 6-His tags.

**Table 2.** Characteristics of selected VHH. CHO-mTfR1-EGFP, CHO-hTfR1-EGFP, and CHO-rhTfR1-EGFP cells were incubated 1 h at 4 °C with various concentrations of VHH, to determine their apparent K_d_. The VHHs were detected with a mouse anti-6His (1/1000) and an Alexa647-conjugated anti-mouse secondary antibody (1/400). Measurements were performed using flow cytometry. Data are presented as mean ± SD of at least 3 independent experiments. NB: no binding.

| VHH name | Target | Molecular Weight (Da) | Theoretical pI with tags | Theoretical pI without tags | K<sub>d/app</sub> on mTfR1 (nM) | K<sub>d/app</sub> on hTfR1 (nM) | K<sub>d/app</sub> on rhTfR1 (nM) |
| --- | --- | --- | --- | --- | --- | --- | --- |
| C5 | TfR1 | 14854 | 6.31 | 8.03 | 60 (± 21) | 3.1 (± 1.5) | 4.2 (± 1.1) |
| B8 | TfR1 | 14948 | 6.05 | 6.49 | 7.1 (± 4.2) | 2.4 (± 1.4) | 2.1 (± 0.3) |
| B6 | TfR1 | 15159 | 6.32 | 8.06 | NB | 2.7 (± 1.4) | NB |
| H3 | TfR1 | 15009 | 6.05 | 6.49 | 56 (± 27) | 2.4 (± 0.9) | 1.5 (± 0.3) |
| D12 | unknown | 15073 | 6.15 | 6.85 | NB | NB | NB |

To evaluate VHH cross-species reactivity, we performed immunocytochemistry experiments illustrated with VHH C5 and B8 in Figure 1A and 1B respectively. Using the stable CHO cell lines expressing the mouse, human or rhesus monkey TfR1, we demonstrated the co-localization between the TfR1-EGFP from the different species and VHH immunostaining. Qualitatively, C5 labelling was less intense on mouse TfR1 compared to human and rhesus monkey TfR1, suggesting a higher affinity of C5 for the human and rhesus monkey compared to the mouse receptor, while B8 immunostaining was more homogeneous. We obtained similar results as those with C5 when evaluating the H3 VHH on the TfR1 variants of the 3 species. There was no binding of B6 VHH on mouse and rhesus monkey TfR1 expressing cells (not shown). There was no binding of the C5 and B8 VHHs on the parental WT CHO cell line (Figure 1) nor with D12 VHH negative control on TfR1 expressing cells (not shown).

**Figure 1.**
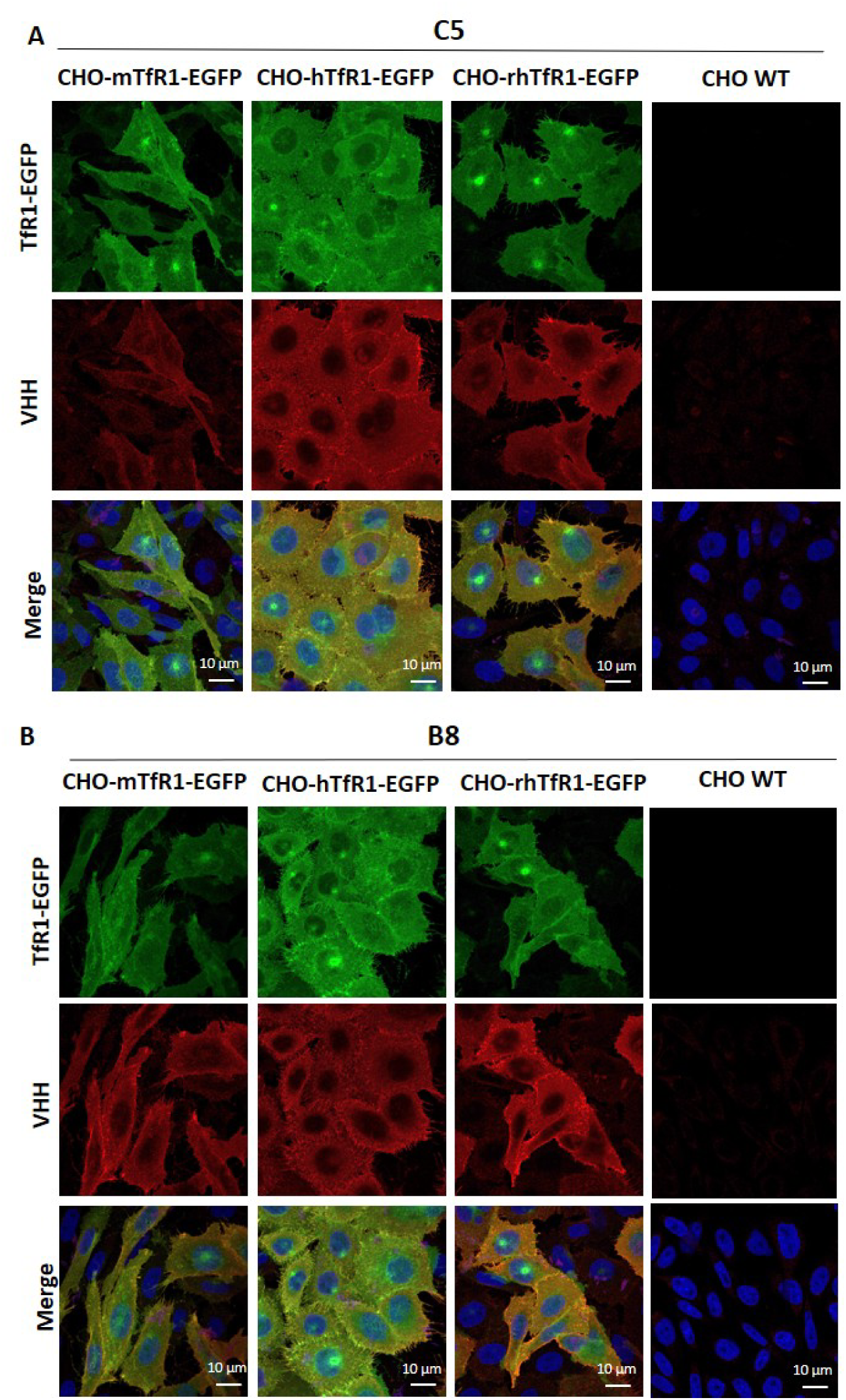
Cell surface binding of C5 and B8 VHHs on CHO cells stably expressing mTfR1, hTfR1, and rhTfR1. Representative confocal photomicrographs of C5 and B8 VHHs binding to CHO cells stably expressing mTfR1-EGFP, hTfR1-EGFP, rhTfR1-EGFP compared to CHO WT. (A, B) CHO-hTfR1-EGFP, CHO-mTfR1-EGFP, CHO-rhTfR1-EGFP (green) and CHO WT cells were incubated 1 h at 37 °C with the C5 (A) and B8 (B) VHHs at 50 µg/ml. VHHs were detected following PFA fixation and Triton X-100 permeabilization of cell membranes, with a mouse anti-cMyc (1/1000) and an Alexa594-conjugated anti-mouse secondary antibody (1/800, red). Cell nuclei were labeled with Hoechst#33342 at 0.5 µg/ml (blue). Co-labeling appears in yellow/orange in the merged pictures.

The apparent affinities (K_d/app_) of the VHHs for mouse, human and rhesus monkey TfR1 were assessed by flow cytometry on TfR1-EGFP cells (Table 2). The C5, B8 and H3 VHHs bound the mouse TfR1 with a K_d/app_ ranging between 7.1 nM and 60 nM. These VHHs bound the human and rhesus monkey TfR1 cells with a similar high affinity within a nanomolar range (K_d/app_ between 1.5 and 4.2 nM). The B6 VHH was human-specific and bound the human TfR1 with a 2.7 nM apparent affinity. Interestingly, the C5 and H3 VHHs showed the same affinities on the TfR1 of the 3 species despite their slightly differing sequences. For all VHHs, apparent affinities were 3- to 20-times higher for human TfR1 than for mouse TfR1. No binding was observed with the irrelevant VHH D12 even at high concentrations. Using SPR, we further determined the affinity (K_D_) and binding kinetic parameters (association rate (k_on_) and dissociation rate (k_off_)) of C5 and B8 VHHs for recombinant mouse, human and rhesus monkey TfR1 ectodomains fused to a mouse IgG2a Fc (mFc), (Figure 2 and Table 3). The K_D_ and K_d/app_ on mouse TfR1 and rhesus monkey TfR1 were in the same range but affinities were more than 1-log lower than those measured using a cell-based assay on human TfR1, suggesting a bias between the two approaches. Based on the SPR data, the k_off_ was found to be the main parameter influencing K_D_ differences between species since it varied 2 to 3 orders of magnitude for C5 and B8 respectively, while the k_on_ remained in the same range. Globally the affinities of C5 and B8 for TfR1 can be ranked as intermediate on mouse TfR1, strong on rhesus monkey TfR1 and very strong on human TfR1, with B8 presenting a 3- to 6-fold stronger binding than C5.

**Figure 2.**
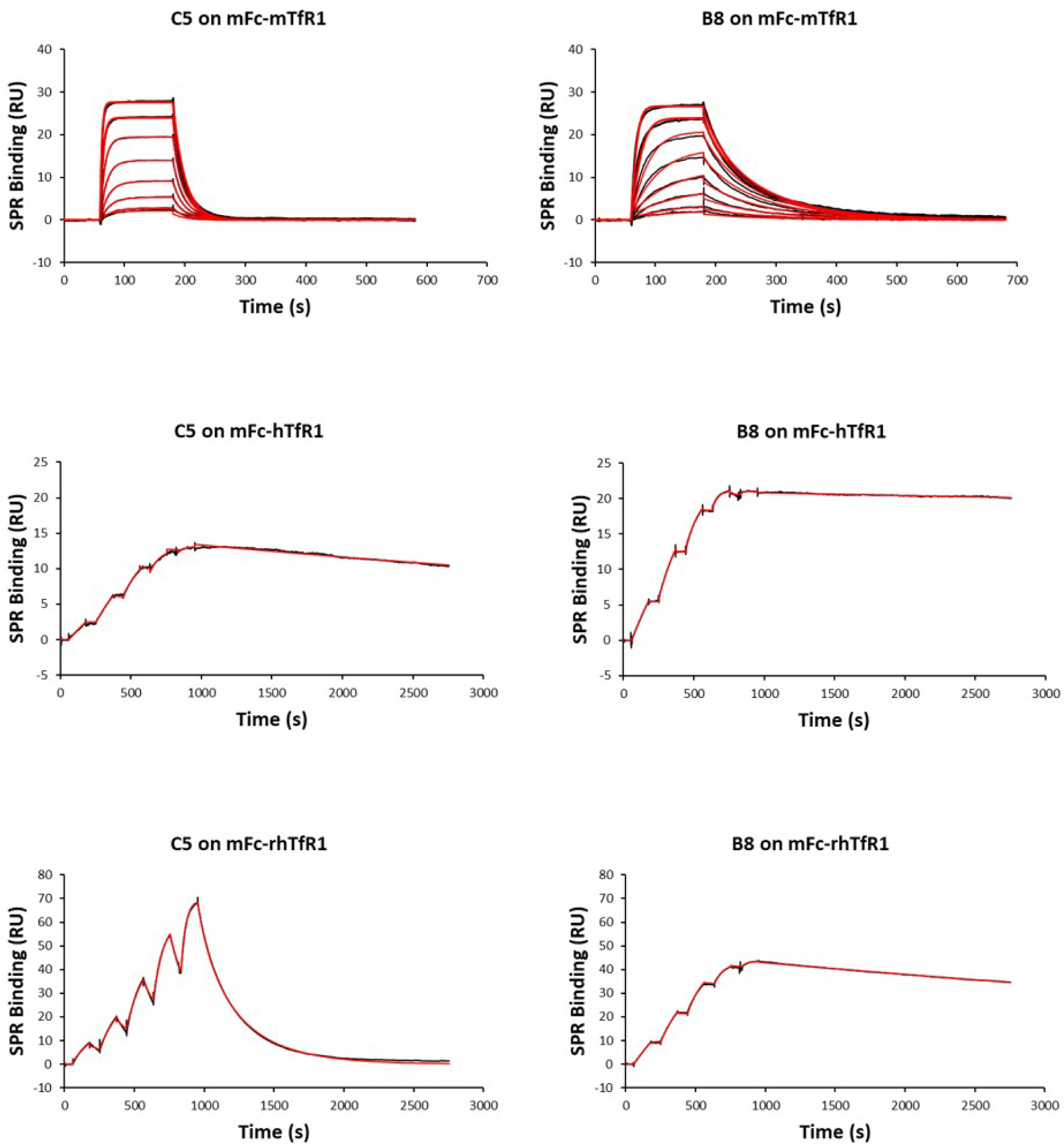
Characterization of C5 and B8 binding to mFc-mTfR1, mFc-hTfR1 and mFc-rhTfR1 by SPR. Increasing concentrations of C5 or B8 were injected over immobilized mFc-mTfR1, mFc-hTfR1 or mFc-rhTfR1 using SCK protocol (2.5 - 40 nM) except on mFc-mTfR1 where MCK protocol (5 - 640 nM) was performed. The red traces correspond to a global fit of the experimental data (black traces) with a 1:1 interaction model. Results are representative of at least 3 independent experiments.

**Table 3.**
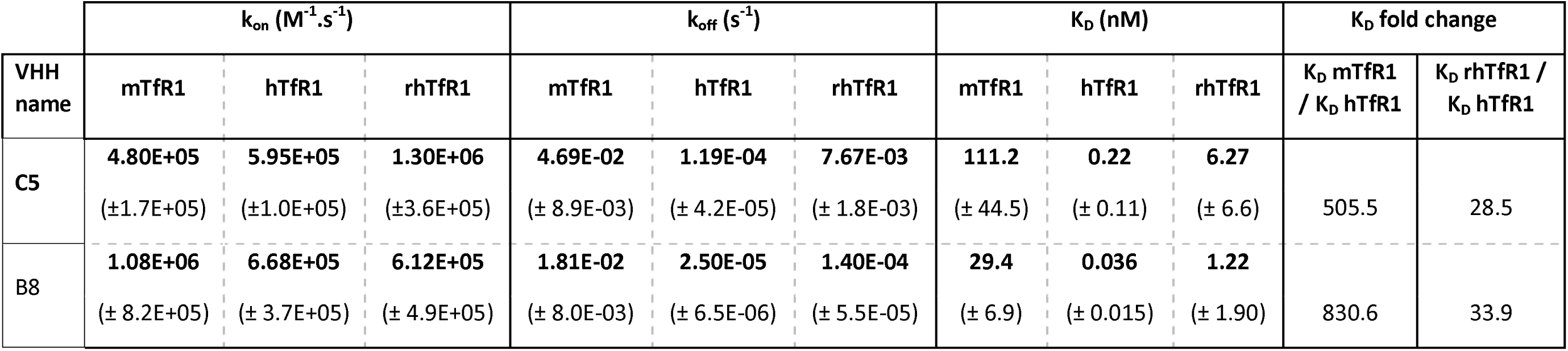
C5 and B8 Affinity (K_D_) and binding kinetic parameters (k_on_, k_off_) determined by SPR. The affinity (K_D_) and binding kinetic parameters (association rate (k_on_) and dissociation rate (k_off_)) of C5 and B8 VHH for recombinant mouse, human and rhesus monkey TfR1 ectodomains fused to a mouse IgG2a Fc (mFc) were determined using SPR and SCK (hTfR1 and rhTfR1) or MCK (mTfR1) protocols. Data are presented as mean ± SD of 3 independent experiments.

We next assessed whether the VHHs interfered with transferrin (Tf) binding to TfR1 as described in the scheme of Figure 3A. Competition assays showed that all VHHs retained their binding even in the presence of excess Tf, and conversely, VHHs did not block Tf binding, indicating they targeted a distinct epitope (Figures 3B–C). Furthermore, C5, B8, and H3 competed with each other for TfR1 binding, suggesting a shared epitope, while B6 did not, confirming it binds a different region (data not shown).

**Figure 3.**
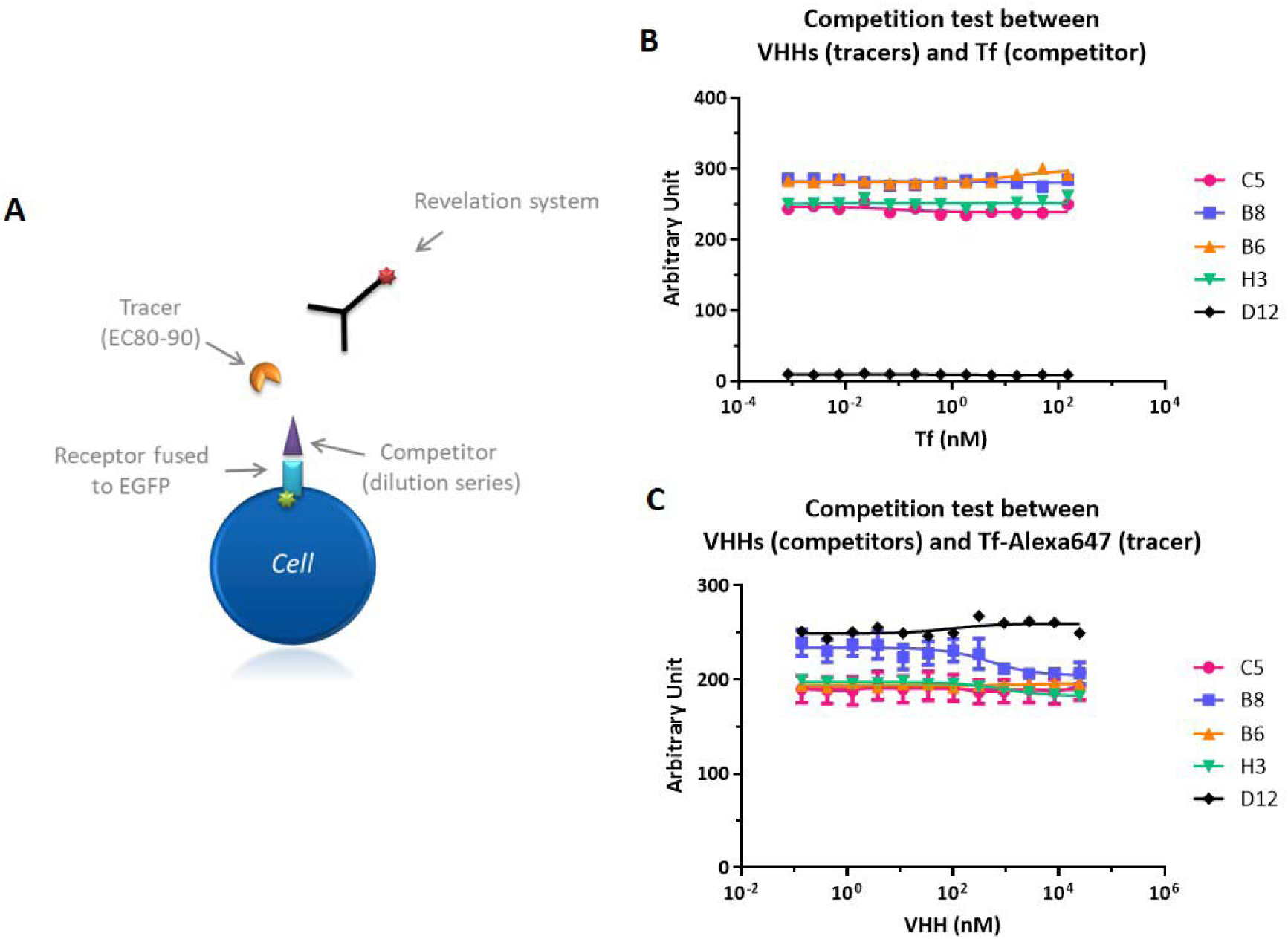
Competition assays between VHHs and Tf. (A) Principle of competition test. In a first step, CHO-hTfR1-EGFP cells were incubated 1 h at 4 °C with the competitor in dilution series. Second, the tracer at EC90 was added and incubated for 1 h at 4°C. Tracer was then revealed with the appropriate revelation system. Measurements were performed using flow cytometry. The ratio of fluorescence intensity for each point was normalized with the corresponding EGFP signal (receptor expression) and gave rise to the arbitrary unit. (B) CHO-hTfR1-EGFP cells were incubated with the competitor (Tf). Tracers (VHHs) at EC90 were then added and detected with a mouse anti-cMyc antibody (1/50) and an Alexa647-conjugated anti-mouse secondary antibody (1/200). (C) CHO-hTfR1-EGFP cells were incubated with competitors (VHHs). Tracer (Tf-Alexa647) at EC90 was then added and detected directly. Data are presented as mean ± SEM of 3 independent experiments.

### Characterization of the C5 and B8 VHH functional epitopes on human TfR1

Next, a two-step approach was used to characterize the functional epitopes of the C5 and B8 parental VHHs considering that they belong to the same family (similar CDR3) while presenting distinct TfR1 binding properties across species. First, a set of human TfR1 variants with single mutations distributed throughout the human TfR1 sequence were expressed in FreeStyle HEK293 cells. Binding assays based on flow cytometry led us to investigate in more detail the area around positions H318, K633 and R719, suggesting an epitope at the dimer interface. Again, no competition was found between the C5 or B8 VHHs and Tf suggesting that the C5 and B8 epitopes do not overlap with the Tf binding site. Second, a series of 12 human TfR1 mono- or bi-mutants (listed in Figure 4A) containing the hypothetic epitope zone was generated to confirm and refine the localization of the C5 and B8 VHH epitopes. Following FreeStyle HEK293 cells transfection with the 12 human TfR1 mutants, a qualitative cytometric evaluation of their loss of binding was compared to the binding of C5 and B8 VHHs to wild-type human TfR1 (Figure 4B). Tf was also used in this experiment to evaluate the level of expression of properly folded mutated TfR1. The mutated positions that have no effect, a moderate deleterious effect, and a strong or total deleterious effect on C5 and B8 VHH binding were identified and illustrated with a color code (Figure 4C and 4D) using the 1SUV structure (Cheng, Zak et al. 2004) of the human TfR1-Tf complex and the Pymol Software (Yuan, Chan et al. 2017). C5 and B8 VHHs engage an epitope close to, but not overlapping, the holo-Tf binding site. It spans both monomers of the human TfR1 receptor homodimer at the apical-helical domain interface. The key residues of the human TfR1 hinge region functionally and strongly involved in the interaction of the C5 and B8 VHHs with the human TfR1 are E634, G636, E728, R732 and E634, G636, E728 respectively. These residues belong to 3 structural regions on human, mouse and rhesus monkey TfR1 and their sequences are listed in Figure 4E. Interestingly, the key residue E634 in the human TfR1 corresponds to D634 in the mouse TfR1. Although E and D residues are both negatively charged, the length of their lateral chains differ. This may contribute to the differences in VHH affinity observed between the TfR1 of both species.

**Figure 4.**
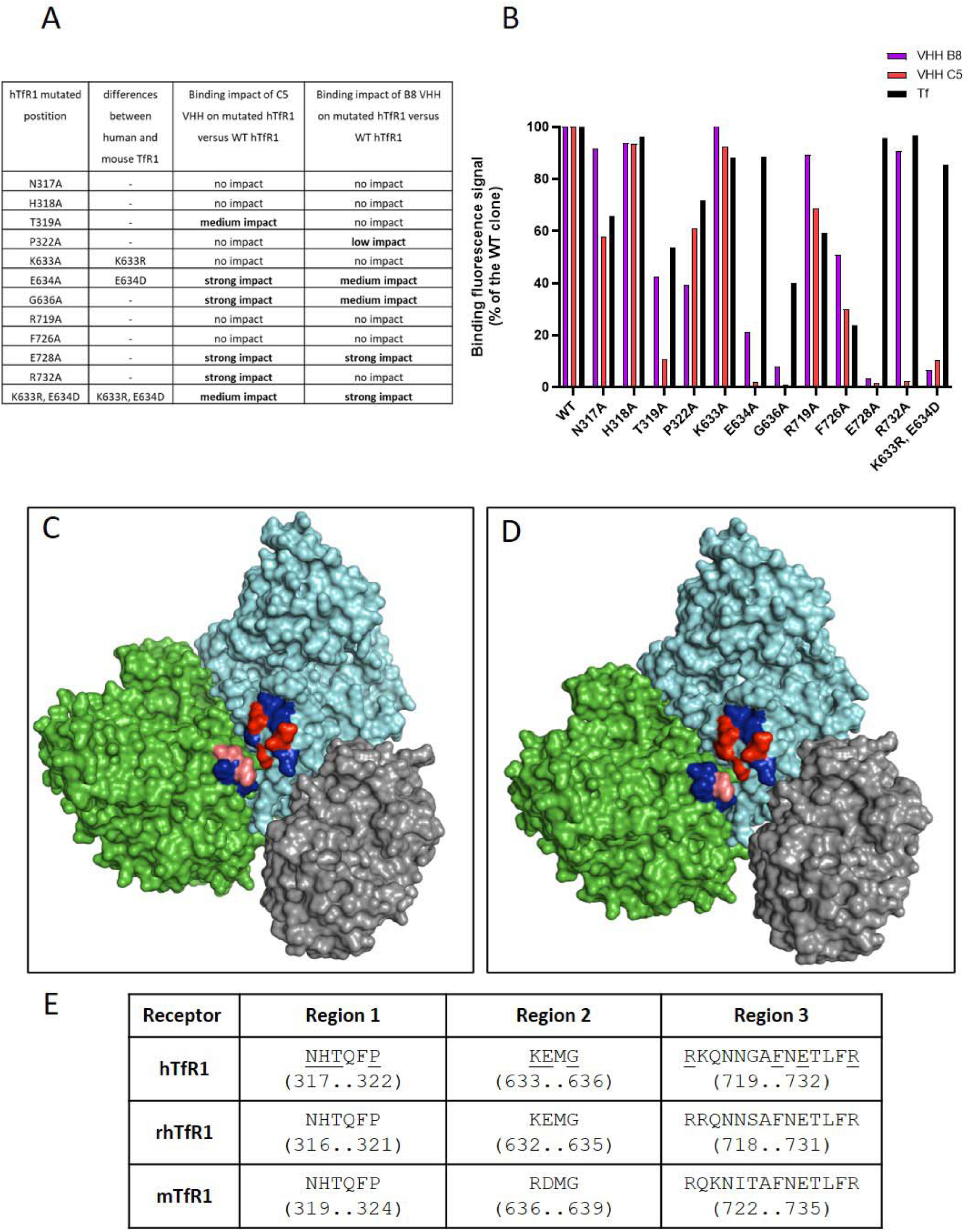
Binding evaluation of VHHs B8 and C5 and Tf on hTfR1 mutants and 3D structure representation of C5 and B8 functional epitopes. (A) Table summarizing the evaluated mutations and their impact on VHHs C5 and B8 binding. (B) Binding fluorescence intensity of VHHs B8 and C5 and Tf on the 12 hTfR1 mutants expressed in HEK293 cells. Fluorescence signals of mutants were normalized with the WT fluorescence. Tf was used to control the level of expression of properly folded hTfR1 mutants. A lower binding signal of the VHHs compared to Tf binding indicated an impact of the mutation on VHH binding. The structure of the hTfR1-Tf complex (1SUV) (Cheng, Zak et al. 2004) was used to illustrate the putative localization of C5 (C) and B8 (D) functional epitopes by using Pymol Software version 3.1.3 (Schrödinger, LLC)(Yuan, Chan et al. 2017). Mutated positions that have no effect on VHH binding were coloured in dark blue, the mutated positions with a moderate deleterious effect on VHH binding were coloured in pink (see arrow in D), and those presenting a strong/total deleterious effect on VHH binding were coloured in red (arrow and arrowhead in D points the epitope positions that do not impact equally the C5 and B8 VHH binding). Each hTfR monomer of the hTfR homodimer was coloured in green and cyan while Tf appears in silver grey. (E) Table summarizing the key residues (underlined) of the human TfR1 hinge region involved in the interaction between the C5 or B8 VHH and the human TfR1. These residues belong to 3 structural regions on human, mouse and rhesus monkey TfR1.

Of note, the C5 or B8 VHH binding region on human TfR1 is 75 % and 91.7 % identical to that of mouse TfR1 and rhesus monkey TfR1, respectively, but 83.3 % and 95.8 % similar, respectively. The root-mean-square deviation (RMSD) between the targeted region on human TfR1 and the predicted targeted region on rhesus monkey or mouse TfR1 receptors were determined. Considering their proximity (RMSD < 0.5 Å), the C5 or B8 binding regions are predicted to be structurally very similar for both mouse and rhesus monkey TfR1 (Supplemental Figure S2). The C5 and B8 VHH binding regions on human TfR1 were distinct from both Tf and H-Ft binding sites that are located in a region of helical and protease-like domains (Eckenroth, Steere et al. 2011) and in the TfR1 receptor apical domain (Montemiglio, Testi et al. 2019), respectively. These findings confirm that C5 and B8 bind a previously uncharacterized epitope at the TfR1 dimer interface, distinct from known ligand binding sites.

### VHH fused to a human IgG1 Fc promotes TfR1-dependent cellular uptake

To establish whether the selected VHHs could promote TfR1-dependent uptake of a protein cargo into cells, we recombinantly fused C5, B8 and D12 (as negative control) to the N-terminus of a human IgG1 Fc region and named the constructs (C5)_2_-hFc*, (B8)_2_-hFc* and (D12)_2_-hFc* (Figure 5A, (*) referring to the presence of effector functions). Due to the fusion to the hFc domain, the VHH, which is normally around 15 kDa, forms a homodimer of approximately 80 kDa. Immunocytochemistry and confocal microscopy were used to validate the binding of the VHH-hFc* fusions to the TfR1 expressed in CHO-mTfR1-EGFP and CHO-hTfR1-EGFP cell lines (Figure 5B). We observed that (C5)_2_-hFc* and (B8)_2_-hFc* bound efficiently to both cell lines, while no binding was detected with the control (D12)_2_-hFc*. Moreover, co-localization of mouse or human TfR1-EGFP with (C5)_2_-hFc* and (B8)_2_-hFc* immunostaining was observed, confirming a TfR1-dependent binding of the Fc fusions (Figure 5B). The apparent affinity of the Fc fusions was determined by flow cytometry-based assay (Table 4). Affinities of the bivalent (C5)_2_-hFc* and (B8)_2_-hFc* fusions were 14- and 2-times higher than monovalent VHHs on CHO-mTfR1-EGFP, with K_d/app_ of 4.3 nM and 3.5 nM, respectively. No binding was observed with the negative control (D12)2-hFc* on both cell lines. Surprisingly, despite the bivalent display of VHHs by the Fc region, (C5)_2_-hFc* and (B8)_2_-hFc* showed affinities 1.7- and 1.9-times lower than monovalent VHHs on CHO-hTfR1-EGFP, with K_d/app_ of 5.3 nM and 4.5 nM (3.1 nM and 2.4 nM for VHHs alone), respectively. We also fused the VHHs to the C-terminus of the hFc and confirmed binding to both human and mouse TfR1 with apparent affinities within the nanomolar range (data not shown). Binding and uptake of the (VHH)_2_-hFc* fusions by mouse, rat and human primary brain endothelial cells (mBMEC, rBMEC and hBMEC respectively) was assessed. Immunocytochemistry showed clear punctate distribution of (C5)_2_-hFc* on mBMEC, rBMEC and hBMEC while (D12)_2_-hFc* showed no labelling (Supplemental Figures S3A, S3B). Quantification of the binding and uptake by flow cytometry on mBMEC and rBMEC showed a 4-to-10-fold increase of (C5)_2_-hFc* labelling compared to (D12)_2_-hFc* (Supplemental Figure S3C).

**Figure 5.**
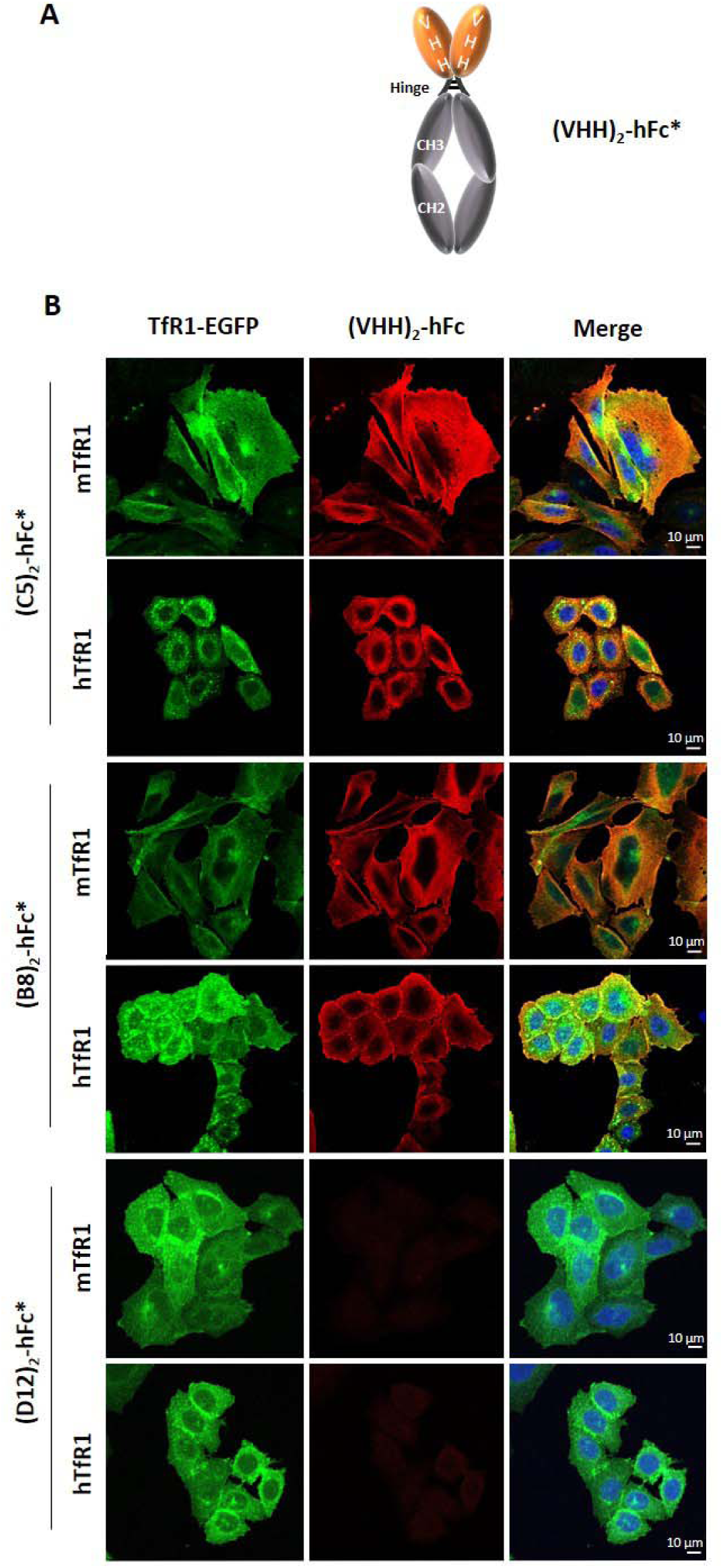
Cell surface binding of (C5)_2_-hFc*, (B8)_2_-hFc* and (D12)_2_-hFc* fusions on hTfR1- and mTfR1-expressing CHO cells. (A) Schematic representation of a (VHH)_2_-hFc*. The VHH is fused to the hinge region, at the N-terminus of the CH2 and CH3 domains of the crystallizable fragment (Fc) of a human IgG1 (hFc). (hFc*) Refers to hFc encompassing effector functions. (B) Representative confocal photomicrographs of CHO-mTfR1-EGFP and CHO-hTfR1-EGFP cells (green) incubated 1 h at 37 °C with 50 nM of (C5)_2_-hFc*, (B8)_2_-hFc* and (D12)_2_-hFc*, detected post-PFA fixation with an Alexa594-conjugated anti-hFc antibody (1/1000, red). Cell nuclei were labeled with Hoechst#33342 at 0.5 µg / ml (blue). Co-labeling appears in yellow/orange in the merged pictures.

**Table 4.**
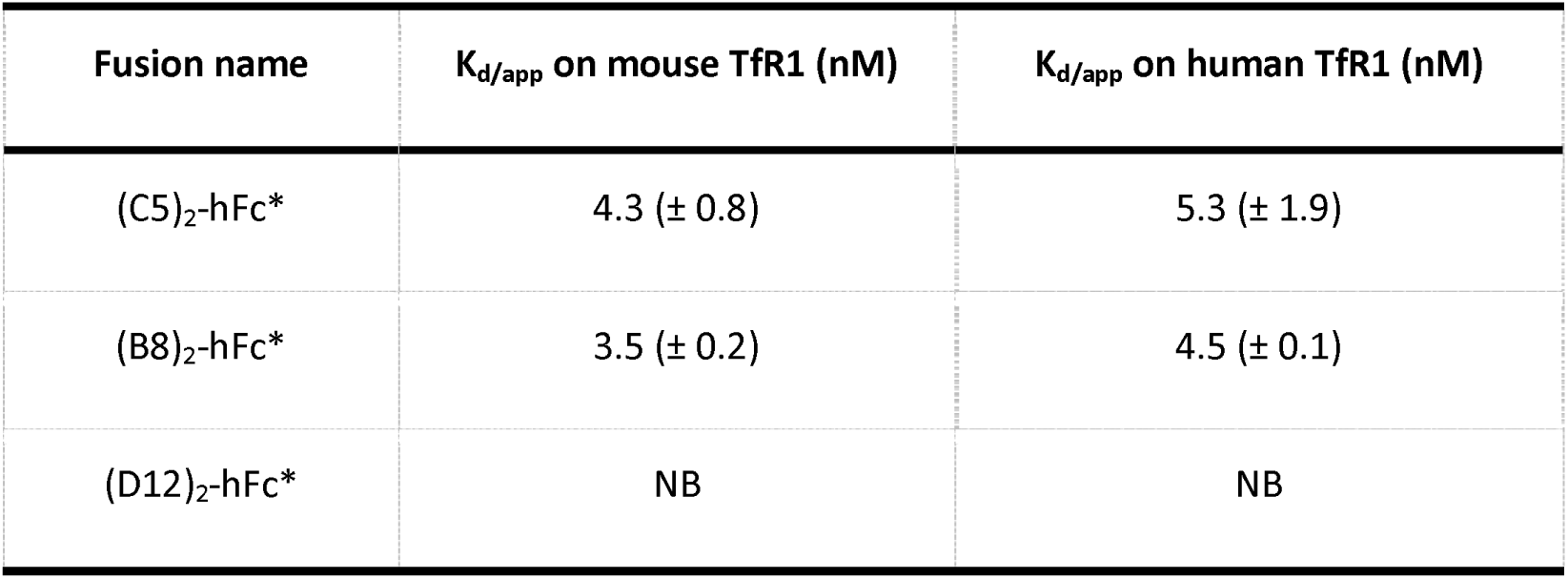
Apparent affinities for mouse and human TfR1 of (VHH)_2_-hFc* fusions. (*) refers to the presence of effector functions. Summary of the apparent affinities (K_d/app_) of (VHH)_2_-hFc toward mTfR1 and hTfR1 determined on CHO-mTfR1-EGFP and CHO-hTfR1-EGFP cells. Each value shows the mean of at least n = 3 independent experiments. NB no binding.

| Fusion name | K<sub>d/app</sub> on mouse TfR1 (nM) | K<sub>d/app</sub> on human TfR1 (nM) |
| --- | --- | --- |
| (C5)<sub>2</sub>-hFc* | 4.3 (± 0.8) | 5.3 (± 1.9) |
| (B8)<sub>2</sub>-hFc* | 3.5 (± 0.2) | 4.5 (± 0.1) |
| (D12)<sub>2</sub>-hFc* | NB | NB |

### Pharmacokinetics of VHH-hFc and brain delivery in C57BL/6 mice after i.v. administration

Having shown that the C5- and B8-hFc fusions were promising molecular vector candidates *in vitro,* we performed *in vivo* experiments in C57BL/6 mice to evaluate their potential to shuttle a prototypic protein cargo across the BBB, into the CNS. We used the above-mentioned hFc for its intrinsic properties such as increased plasma half-life, antibody like features and potential to be engineered to display VHH in dimeric and monomeric format. In order to avoid *in vivo* toxicity related to hFc effector functions, as previously described (Couch et al., 2013; Weber et al., 2018), we introduced the L234A and L235A LALA mutations (amino acid numbering of a full-length antibody) in the hFc fragment. For the following *in vivo* experiments, we also generated a C5 negative control (C5neg) by mutating 2 amino acid residues in the CDR3 of the C5 VHH (Y105A, L107A IMGT numbering), leading to near-total abrogation of binding to TfR1. We used N-terminal genetic fusions, and when appropriate, knob-into-hole (KIH) mutations, to generate VHH-hFc molecules with two or one VHH per molecule allowing theoretically bivalent or monovalent binding to mouse TfR1, respectively (Figure 6A). Indeed, binding valency of TfR1 ligands has been described as a critical parameter for TfR1-mediated transport across the BBB (Niewoehner, Bohrmann et al. 2014).

**Figure 6.**
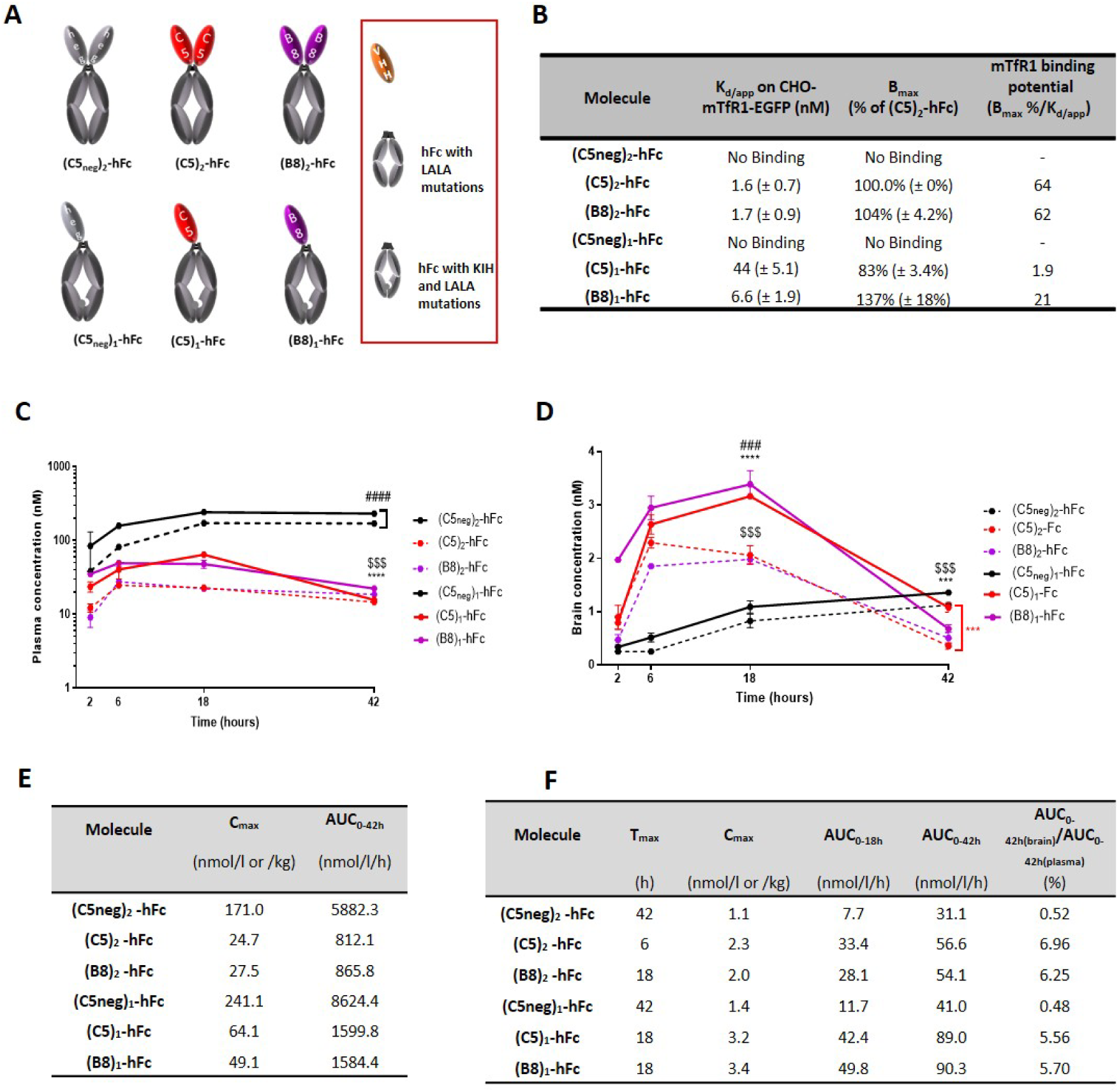
Plasma PK and brain uptake in C57BL/6mice of VHH_TfR_-hFc with a monovalent or bivalent binding mode to mTfR1. (A) Scheme of the VHH-hFc molecules (encompassing LALA mutations in the hFc to reduce effector functions) with two ((C5neg)_2_-hFc; (C5)_2_-hFc; (B8)_2_-hFc)) or one ((C5neg)_1_-hFc; (C5)_1_-hFc; (B8)_1_-hFc)) VHH per molecule allowing bivalent or monovalent binding, respectively. (B) Table summarizing the binding properties of the VHH_TfR_-hFc constructs: VHH-hFc K_d/app_, B_max_ percentage relative to (C5)_2_-hFc and binding potential toward mTfR1 determined on CHO-mTfR1-EGFP cells. Each value shows the mean of at least n = 3 independent experiments. (C, D) VHH-hFc concentration in plasma and total brain respectively at 2 h, 6 h, 18 h and 42 h post-injection. C57BL/6 mice were injected s.c. (35 nmol / kg) with (C5neg)_2_-hFc (black dots and dashed line), (C5neg)_1_-hFc (black dots and solid line), (C5)_2_-hFc (red dots and dashed line), (C5)_1_-hFc (red dots and solid line), (B8)_2_-hFc (purple dots and dashed line), (B8)_1_-hFc (purple dots and solid line). Each symbol represents the mean (±SEM) value of n=4 mice. Only significant differences are shown up to 42 h post-injection for plasma and up to 18 h and 42 h for brain. Statistical differences were calculated with one-way ANOVA with Tukey’s post-hoc test. ***p < 0.001 and ****p < 0.0001 when comparing (VHH)_1-_hFc to (C5neg)_1_-hFc. $$$ p < 0.001 and $$$$ p < 0.0001 when comparing (VHH)_2-_hFc to (C5neg)_2_-hFc. ### p < 0.001 and #### p < 0.0001 when comparing (VHH)_1-_hFc to (VHH)_2_-hFc. (E) Table summarizing the plasma C_max_ and AUC_0-42h_ of each construct calculated with Kinetika software V5.1. (F) Table summarizing the brain T_max_, C_max_, AUC_0-18h_ and AUC_0-42h_ of each construct calculated with Kinetika software V5.1. Brain- to-plasma ratio percentage was calculated by comparing the AUC_0-42h_ (brain) / AUC_0-42h_ (plasma) X 100.

A CHO-mTfR1-EGFP cell-based assay was used to determine the apparent affinity of VHH-hFc, the B_max_ percentage relative to (C5)_2_-hFc, which corresponds to the maximum receptor binding capacity, and the binding potential (ratio of B_max_ percentage over K_d/app_), which characterizes the binding properties of a given VHH-hFc to mouse TfR1 (Figure 6B). We observed a higher apparent affinity for the bivalent (VHH)_2_-hFc compared to the monovalent (VHH)_1_-hFc, consistent with an avidity effect. The mouse TfR1 binding potentials were 34-fold and 3-fold higher for the C5-hFc and B8-hFc, respectively, when comparing their bivalent and monovalent versions.

To evaluate the plasma pharmacokinetics (PK) and brain accumulation potential of VHH-hFc constructs with either monovalent or bivalent TfR1 binding, C57BL/6 mice were injected subcutaneously with 35 nmol/kg of (VHH)₂-hFc and (VHH)₁-hFc constructs. Plasma (Figure 6C) and total brain (Figure 6D) concentrations were measured using an anti-hFc ELISA assay at 2, 6, 18, and 42 hours post-injection. Up to 42 hours, plasma exposure (AUC_0-42h_) for TfR1-targeting constructs with two or one C5 or B8 VHH was 7- to 5.4-fold lower than that of the negative control constructs (C5neg)₂-hFc and (C5neg)₁-hFc, respectively, (Figure 6E). This is consistent with a target-mediated drug disposition (TMDD) effect due to TfR1 binding across tissues. The AUC_0-42h_ of all bivalent (VHH)₂-hFc constructs was 1.8-to 2-fold lower than that of the monovalent (VHH)₁-hFc constructs (Figure 6E). No significant differences in maximum concentration (C_max_) or AUC_0-42h_ were observed when comparing the C5 and B8 VHHs in either bivalent or monovalent formats (Figure 6E). In brain tissue, the PK profiles of TfR1-targeting VHH-hFc constructs displayed a rapid bell-shaped curve, with C_max_ between 6- and 18-hours post-injection, whereas the profiles of negative controls showed a gradual increase over time, reaching T_max_ at 42 hours (Figure 6F). The AUC_0-18h_ for TfR1-targeting versions was 4-fold higher than that of negative controls (C5neg)₂-hFc and (C5neg)₁-hFc and remained higher through 42 hours (Figure 6F), strongly suggesting TfR1-mediated uptake. This TfR1-dependent brain uptake is further supported by the observation that the brain-to-plasma ratio percentages for the bivalent and monovalent C5- and B8-hFc constructs were approximately 13- and 11-fold higher, respectively, than those of the non-targeting C5neg-hFc controls. Overall, the C5- and B8-hFc constructs in their monovalent format demonstrated significantly higher brain exposure than both the negative controls and the bivalent VHH-hFc constructs.

### Induction of hypothermia by VHH-NT(8-13) fusions in C57BL/6 mice after i.v. administration

Having shown that the C5- and B8-hFc fusions were able to reach greater brain concentrations than control molecules, especially with a monovalent binding mode to TfR1, we focused our efforts on evaluating their ability to cross the BBB and elicit efficacy in brain parenchyma. To this aim, the short active version of neurotensin (NT(8-13)), a neuropeptide that can induce a variety of central effects, including HT, but that does not cross the BBB efficiently (Banks, Wustrow et al. 1995, Demeule, Beaudet et al. 2014) was fused to C5neg, C5 and B8 VHHs. Two VHH-NT(8-13) designs enabling monovalent TfR1 binding mode were evaluated, that differ in their molecular weights (Figure 7A). The low MW construct (15 kDa) encompasses the NT(8-13) peptide fused to the C-terminal end of the VHH, a construct hypothesized to have a short *in vivo* plasma half-life. The high molecular weight construct (65 kDa) was generated using the same Fc protraction strategy as previously described, with the VHH fused to the N-terminus of the Fc and the NT(8-13) peptide to its C-terminus (Figure 7A). Moreover, a control (C5)_1_-hFc fusion that binds TfR1 but not the NT receptor (NTSR1) was included to check that HT was not due to TfR1-mediated toxicity related to residual Fc effector functions. The different fusions were purified using an affinity chromatography step followed by a gel filtration step reaching over 95 % purity. As shown in Figure 7B, all the NT(8-13) peptide fusions, including the C5neg-NT(8-13) had a basic pI around 9 for the low MW constructs and around 8 for the high MW constructs. *In vitro*, all the VHH-NT(8-13) fusions were shown to bind to the mouse NTSR1 transiently expressed by Expi293 cells, with a K_i/app_ between 3 and 8.1 nM, similar to that of the NT(8-13) peptide alone (K_i/app_ = 2.9 nM) (Figure 7B). As expected, the control (C5)_1_-hFc did not displace the binding of the reference fluorescent NT(8-13) used in the K_i/app_ cellular assay. Next, we assessed *in vitro* the K_d/app_ of the conjugates for mouse TfR1, the relative B_max_ %, and the mouse TfR1 binding potential of all constructs (Figure 7B). Overall, the mouse TfR1 binding potentials were similar between the low and high MW constructs for a given VHH, with the B8 constructs showing a binding potential at least 20-fold higher than that of the C5 constructs.

**Figure 7.**
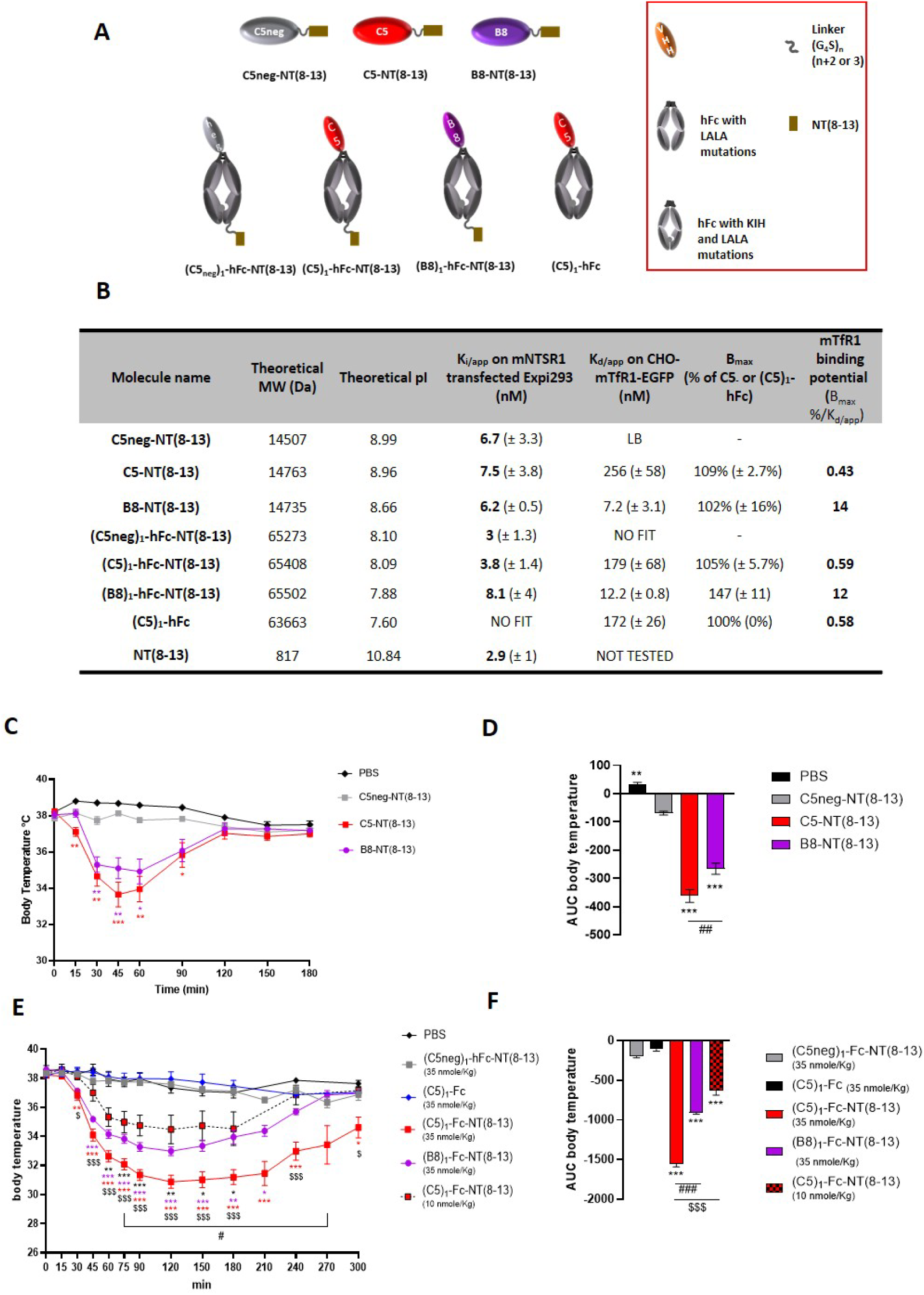
(VHH)_1_-NT(8-13) and (VHH)_1_-hFc-NT(8-13) with a monovalent TfR1 binding mode induce HT in C57BL/6 mice. (A) Scheme of the VHH fusions encompassing or not hFc and NT(8-13). (B) Table summarizing the molecular weight of the fusion molecules, K_i/app_ on mNTSR1-transfected Expi293 cells, K_d/app_, B_max_ and binding potential toward mTfR1 determined on CHO-mTfR1-EGFP cells. Each value shows the mean of n = 2 for K_i/app_ and n = 3 for K_d/app_ independent experiments. (C) C57BL/6 mice body temperature after i.v injection of C5neg-NT(8-13) (grey squares and solid line), C5-NT(8-13) (red squares and solid line) and B8-NT(8-13) (purple dots and solid line) fusions at 500 nmol / kg compared to vehicle (PBS, black diamonds and solid line), n = 4 / time point. Mouse body temperature was measured every 15 min up to 60 min, then every 30 min up to 180 min post-injection. (E) C57BL/6 mice body temperature after i.v injection at 35 nmol / kg of (C5neg)_1_-hFc-NT(8-13) (grey squares and solid line), (C5)_1_-hFc (blue diamonds and solid line), (C5)_1_-hFc-NT(8-13) (red squares and solid line), (B8)_1_-hFc-NT(8-13) (purple dots and solid line) and PBS (black diamonds and solid line), n=4-8 / time point. Additionally, (C5)_1_-hFc-NT(8-13) was administered at 10 nmol / kg (red squares and black dashed line). Mouse rectal temperature was measured every 15 min up to 90 min then every 30 min up to 300 min post-injection. (D) and (F) area under the curve (AUC) of respectively (C) and (E) determined with GraphPad Prism software. Each symbol represents the mean (± SEM) value of n = 4 / 8 mice. Differences between groups were analyzed using one-way ANOVA with Tukey’s post-hoc test. Only significant differences are shown. *p < 0.05, **p < 0.01 and ***p < 0.001 when comparing C5neg-NT(8-13) to C5-NT(8-13) and B8-NT(8-13) or (C5neg)_1_-hFc-NT(8-13) to (C5)_1_-hFc-NT(8-13) and (B8)_1_-hFc-NT(8-13). # p< 0.05, ## p <0.01 and ### p<0.001 when comparing C5-NT(8-13) to B8-NT(8-13) or (C5)_1_-hFc-NT(8-13) to (B8)_1_-hFc-NT(8-13). $ p < 0.05 and $$$ p < 0.001 when comparing (C5)_1_-hFc-NT(8-13), injected at 35 nmol/kg, to (C5)_1_-hFc-NT(8-13) injected at 10 nmol / kg.

The low MW constructs VHH-NT(8-13) at 500 nmol / kg or vehicle (PBS) were administered i.v in C57BL/6 mice (n = 4/time point) and we followed body temperature every 15 min up to 60 min, then every 30 min up to 180 min post-i.v injection (Figure 7C). C5 and B8 low MW constructs both induced rapid (as early as 15 min) and significant HT that was maximal (−3 °C and −4 °C) between 45 min and 60 min post-injection, respectively. For both constructs, body temperature rose back up to baseline (38 °C) at 120 min. Vehicle (PBS) or C5neg-NT(8-13) did not induce HT. AUC evaluation showed that the C5 construct induced a more robust HT than B8 (Figure 7D). We also followed mouse body temperature after i.v injection of the high MW constructs (VHH)_1_-hFc-NT(8-13) or (VHH)_1_-hFc at 35 nmol / kg or vehicle in C57BL/6 mice (n = 4-8 / time point) every 15 min up to 90 min, then every 30 min up to 300 min post-injection. The C5 and B8 high MW constructs both induced a strong significant HT starting at 30 min post-administration that was maximal (−7 °C and −5 °C) between 120 and 210 min post-injection, respectively. Body temperature with (C5)_1_-hFc-NT(8-13) was still lower than that with the control at 300 min, whereas it was back to baseline at 270 min for the B8 construct. Vehicle or (C5neg)_1_-hFc-NT(8-13) failed to induce HT (Figure 7E). Here again, AUC evaluation showed that the C5 high MW construct induced a more robust HT than the B8 high MW construct (Figure 7F). To evaluate a dose-response effect, the high MW C5 construct was administered at the low dose of 10 nmol / kg which also induced a significant (−3 °C) and long lasting (240 min) HT (Figure 7E and 7F). These results clearly demonstrated that i) the TfR1 binding VHHs efficiently deliver pharmacologically active amounts of the NT(8-13) peptide in a dose-dependent manner; ii) the Fc protraction strategy substantially improves the effects of NT(8-13), both in intensity and duration, and iii) the binding potential of the VHHs to mouse TfR1 must be moderate to induce a robust and long-lasting HT.

### Induction of HT by VHH-NT(8-13) variants in B-hTfR mice after i.v administration

Having demonstrated the ability of low and high MW VHH-NT(8-13) to cross the BBB in WT mice in a TfR1 dependent manner, we next assessed their trans-species potential in B-hTfR mice, a transgenic mouse model where exons 4-19 of the mouse *Tfr1* gene that encode the extracellular region of TfR1 were replaced by human *Tfr1* exons 4-19. To this end, the C5 VHH, the best BBB crosser in wild type mice, fused to NT(8-13) was chosen to assess HT in B-hTfR mice following i.v. administration. In line with the affinity, on-rate and off-rate parameters found on human TfR1 for the C5 parental VHH, SPR analysis showed that the K_D_ of C5-NT(8-13) for human TfR1 was sub-nanomolar (0.1 nM). The kinetic parameters indicated a rapid association rate (k_on_ = 1.36 E+06 M^−1^s^−1^) and a slow dissociation rate (k_off_ = 1,60E-04 s^−1^) (Table 5). A strong affinity is potentially detrimental for efficient transcytosis across brain endothelial cells (Niewoehner, Bohrmann et al. 2014, Johnsen, Burkhart et al. 2019). Accordingly, the absence of HT following C5-NT(8-13) injection in B-hTfR mice further confirmed its poor BBB crossing potential (Figure 8A). Thus, C5 and B8 VHH affinity variants were generated by introducing mono- or multiple amino-acid mutations at key positions in the VHH CDRs or frameworks, using alanine-scan and conservative or disruptive amino acid replacement. Humanization strategies where multiple mutations were introduced in the VHH frameworks to increase homology of the C5 and B8 parental VHHs with the human VH germline IGHV3-66 also proved useful for generating affinity variants. In total, more than one hundred C5 and B8 affinity variants were generated (Cohen, David et al. 2019, Cohen, David et al. 2023, David, Cohen et al. 2023). Some were selected according to their human TfR1 binding properties to undertake an *in vivo* screen of their BBB crossing capacity in B-hTfR mice, based on the HT readout. The mutations we introduced in the selected C5 and B8 variants, together with the binding kinetic parameters, and affinities determined by SPR on human and rhesus monkey TfR1 ectodomains with or without NT(8-13) fusion are summarized in Table 5. The typical sensorgrams of VHH-NT(8-13) fusions can be visualized in Supplemental Figure S4 and S5 on human and rhesus TfR1, respectively. First, as expected, the affinities of the C5 and B8 variants for human and rhesus monkey TfR1 were decreased in comparison to those of their parental VHHs (Table 3 and Table 5), shifting from sub-nanomolar/nanomolar range to tens of nanomolar range. Second, the C5 and B8 variants, along with their NT(8-13) fusions, showed better cross-species reactivity between human and rhesus monkey TfR1 compared to their parental VHHs, which had affinity ratios of 28.5 and 33.9, respectively (Table 3 and Table 5). Notably, with a rhesus monkey/human TfR1 affinity ratio close to one, B8V31, B8V40, B8V31h3 and B8V31h5 bearing the F30K or A54S mutations (IMGT numbering) demonstrated a nearly perfect human-rhesus monkey TfR1 cross-reactivity. Surprisingly, the VHH-NT(8-13) variants presented a slight K_D_ decrease of around 1.5- to 4-fold in comparison with the same variants without NT(8-13) (Table 5), meaning that the vectorized payload can influence the TfR1 binding properties of the final conjugate. Overall, K_D_ on human TfR1 of the VHH-NT(8-13) variants selected for *in vivo* evaluation ranged from around 0.1 nM to 250.8 nM, thus covering 3 logs of affinity, and these VHH-NT(8-13)s were all able to displace the binding of the reference NT(8-13) peptide to mouse NTSR1 in K_i/app_ assays (Supplemental Table S3).

**Figure 8.**
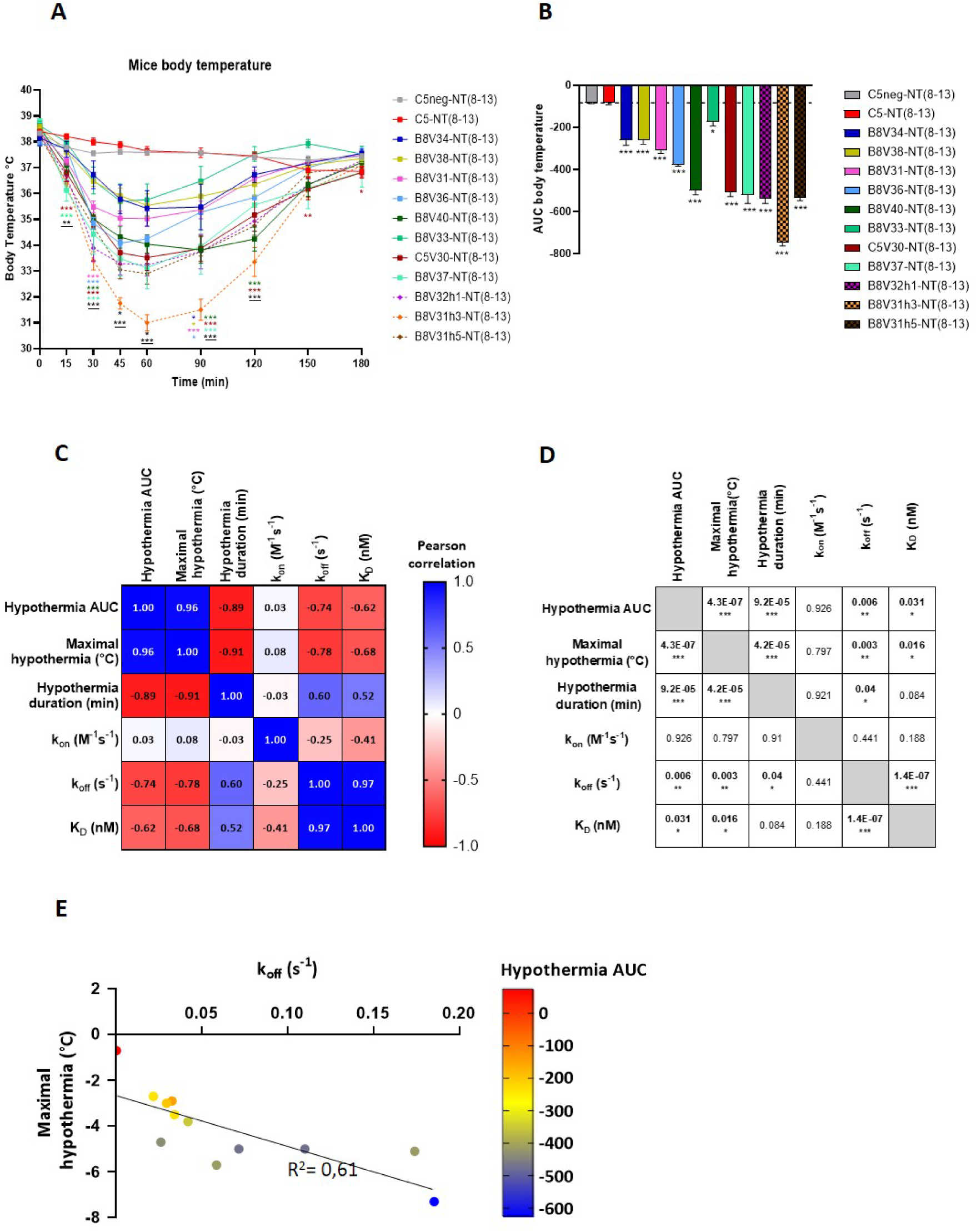
C5 and B8-NT(8-13) variants induce HT in B-hTfR mice expressing the human TfR1 ectodomain in correlation with their TfR1-binding properties. (A) B-hTfR mouse body temperature after i.v injection of C5neg-NT(8-13) (grey squares and solid line), C5-NT(8-13) (red squares and solid line), B8V34-NT(8-13) (blue squares and solid line), B8V38-NT(8-13) (yellow-green squares and solid line), B8V31-NT(8-13) (pink squares and solid line), B8V36-NT(8-13) (light-blue squares and solid line), B8V40-NT(8-13) (dark-green squares and solid line), B8V33-NT(8-13) (green squares and solid line), C5V30-NT(8-13) (burgundy squares and solid line), B8V37-NT(8-13) (light-green squares and solid line), B8V32h1-NT(8-13) (purple diamonds and dashed line), B8V31h3-NT(8-13) (orange diamonds and dashed line) or B8V31h5-NT(8-13) (brown diamonds and dashed line) at 500 nmol / kg (n = 4-23 / time point). Mouse rectal temperature was measured every 15 min up to 60 min, then every 30 min up to 180 min post-injection. Each symbol represents the mean (±SEM) value of n = 4 / 23 mice. Differences between groups were analyzed using one-way ANOVA with Dunnett’s post-hoc test. Only significant differences are shown. *p < 0.05, **p < 0.01 and ***p < 0.001 when comparing C5neg-NT(8-13) to C5-NT(8-13) and C5/B8-NT(8-13) variants. <u>**p</u> < 0.01 and <u>***p</u> < 0.001 when comparing C5neg-NT(8-13) to humanized VHH-NT(8-13). (B) area under the curve (AUC) of (A) with GraphPad Prism software. (C) Heatmap of the Pearson’s correlation coefficient matrix (r) between 6 variables (Hypothermia AUC=AUC_VHH-NT(8-13)_-AUC_C5neg-NT(8-13)_), maximal HT, HT duration, k_on_, k_off_ and K_D_ determined with GraphPad Prism software. Color intensity is proportional to the correlation coefficients (blue indicates a positive correlation between two variables, red indicates a negative correlation between two variables). (D) p value for each calculated correlation coefficient determined with GraphPad Prism. Significant p values (p < 0.05) are in bold. (E) Linear regression between maximal HT and k_off_. The dot color is related to the HT AUC value (red no HT, bleu strong HT).

**Table 5.**
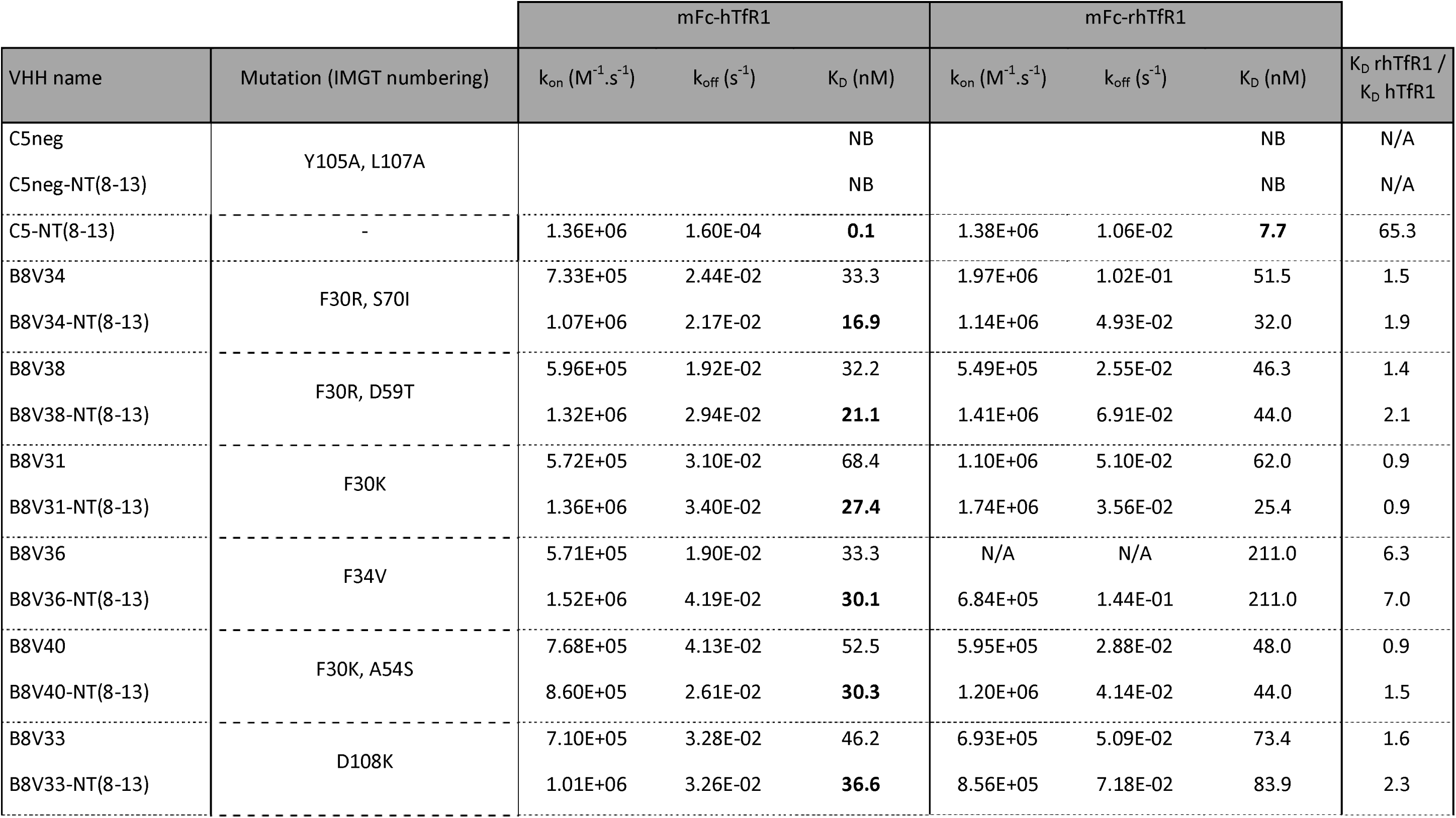
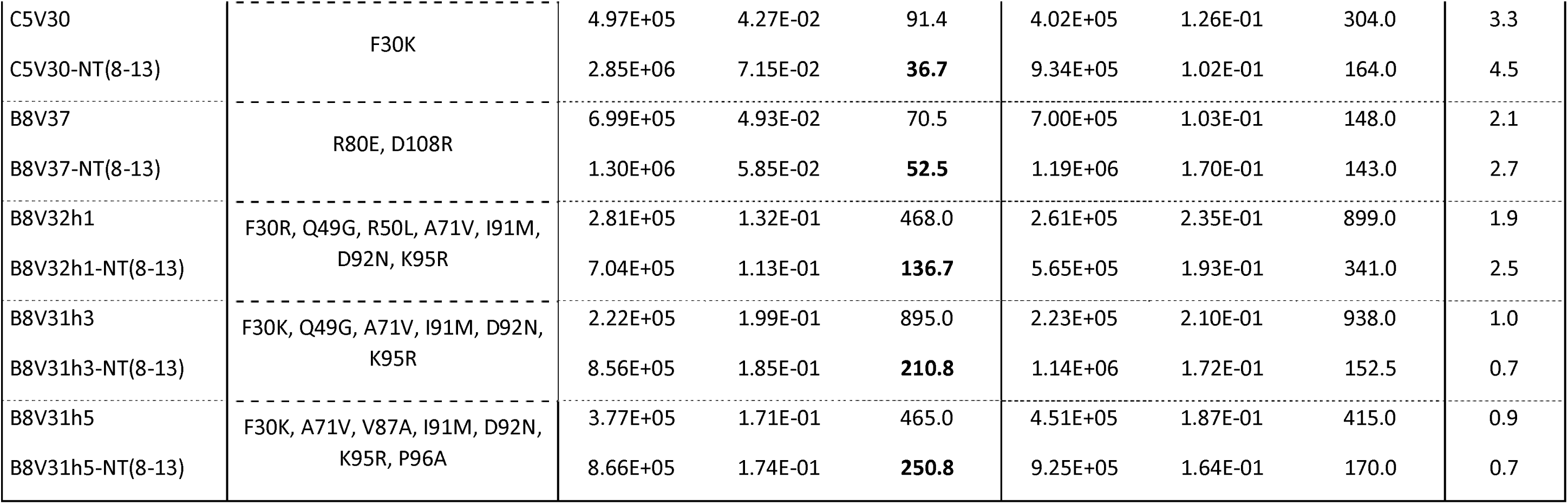
VHH and VHH-NT variants affinity and binding parameters determined by SPR. The affinity (K_D_) and binding kinetic parameters (association rate (k_on_) and dissociation rate (k_off_)) of C5 and B8 VHH variants and their NT(8-13) fusions for recombinant human and rhesus monkey TfR1 ectodomains fused to a mouse IgG2a Fc (mFc) were determined using SPR and MCK or SCK protocols. Data are presented as mean ± SD of 3 independent experiments for hTfR1 and n≥1 for rhTfR1. NB No binding, NT Not tested, N/A Not Applicable.

Next, we evaluated the potential of the VHH-NT(8-13) affinity variant fusions and the control C5neg-NT(8-13) to induce HT after single i.v. injection at 500 nmol / kg in B-hTfR mice. Mouse body temperature was monitored for 3 hours. As mentioned above, the parental C5-NT(8-13) and the C5neg-NT(8-13) control did not induce HT (Figure 8A). In contrast, all the selected C5-NT(8-13) and B8-NT(8-13) affinity variants induced a significant HT that was maximal (from −2 °C to −7 °C) between 60 and 90 min post-injection. For all the VHH-NT(8-13) affinity variants, body temperature was back to baseline (38 °C) 180 min post-injection. Comparison of body temperature AUC (Figure 8B) revealed that B8V31h3-NT(8-13), with a high K_D_ for human TfR1 (210.8 nM), induced the strongest HT, while B8V34-NT(8-13), with a low K_D_ (16.9 nM), induced only moderate HT (Figure 8B). A Pearson correlation matrix (Figure 8C) was used to investigate the potential relationship between human TfR1 binding parameters described in Table 5 (i.e. k_on_, k_off_ and K_D_) and HT data compiled in Table 6 (i.e. maximum temperature reduction, AUC _VHH-NT(8-13)_ relative to AUC_C5neg-NT(8-13)_ and duration of HT). The p-values calculated from the Pearson correlation matrix, indicating -or not-the significance of a given correlation, are shown in Figure 8D. No relationship was found between the k_on_ and the human TfR1 binding parameters but a significant inverse relationship (p-value ≤ 0.05) was found between the affinity (K_D_), the maximal HT and the AUC, and between the k_off_ and all the HT parameters. With a correlation coefficient of −0.78 and a p-value of 0.003, the inverse relationship between the dissociation rate (k_off_) and the maximal temperature reduction was high and significant, with a linear regression shown in Figure 8E. All together these correlations indicated that with a K_D_ up to 210.8 nM and a k_off_ up to 0.185, the lower the affinity for human TfR1 and the faster the dissociation rate from TfR1, the stronger the HT. Interestingly, while the C5 and B8 affinity variants lost some mouse TfR1 binding potential (not shown) and became selective for the human/rhesus monkey TfR1, their human/rhesus TfR1 binding parameters remained close to those of C5 and B8 parental VHHs on mouse TfR1. This suggests that a surrogate approach can be used to conduct preclinical pharmacology and safety studies in mouse. For example, B8V37 or C5V30 have k_on_, k_off_ and K_D_ values on human TfR1 that are very close to those of C5 on mouse TfR1. In the same vein, the binding parameters of B8V34 on human TfR1 are close to those of the B8 parental VHH on mouse TfR1. Additionally, *in vivo*, B8V37-NT(8-13) and C5V30-NT(8-13) or B8V34-NT(8-13) induced similar HT profiles in B-hTfR mice to those induced by C5-NT(8-13) and B8-NT(8-13) in WT mice, respectively.

**Table 6.**
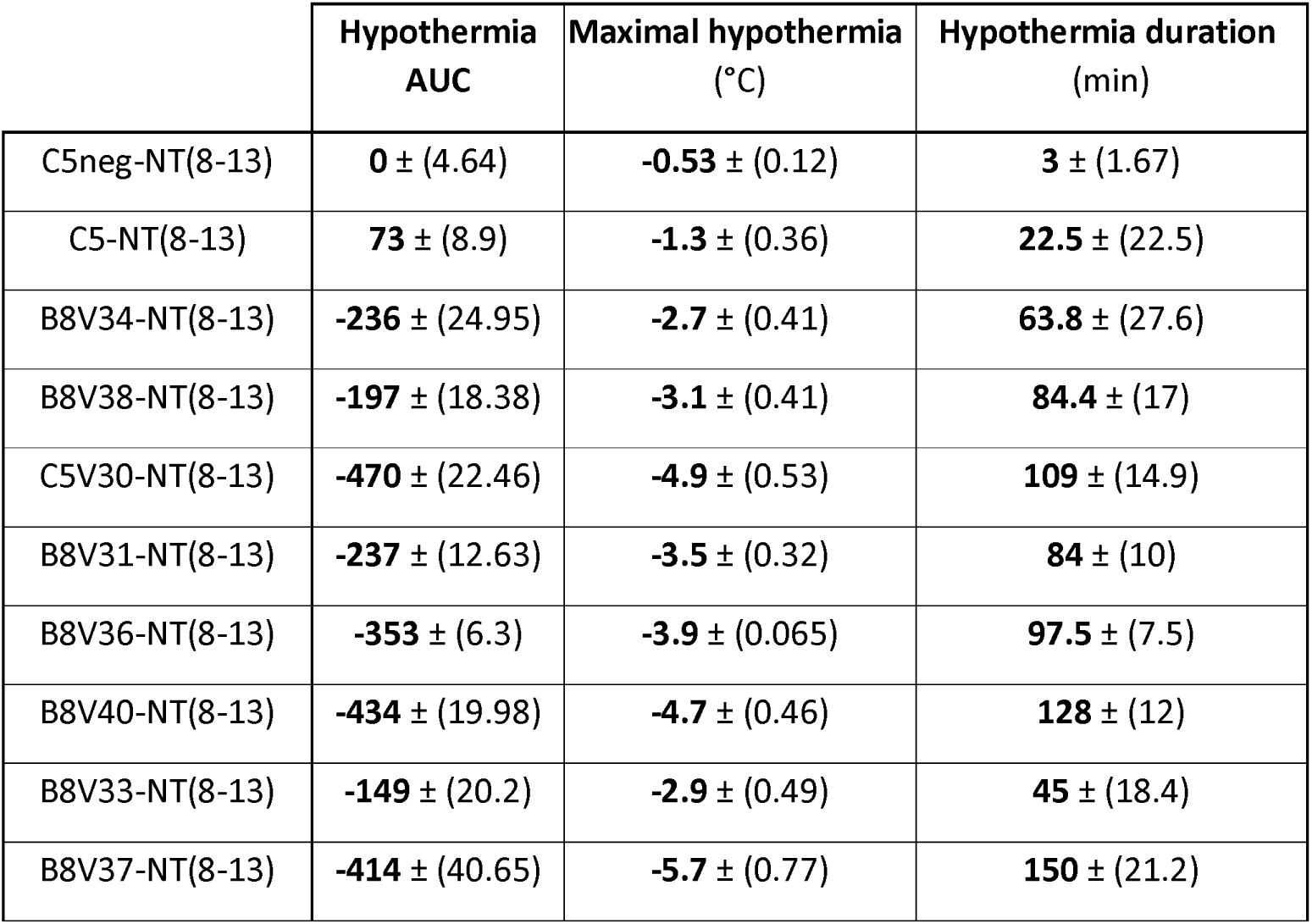
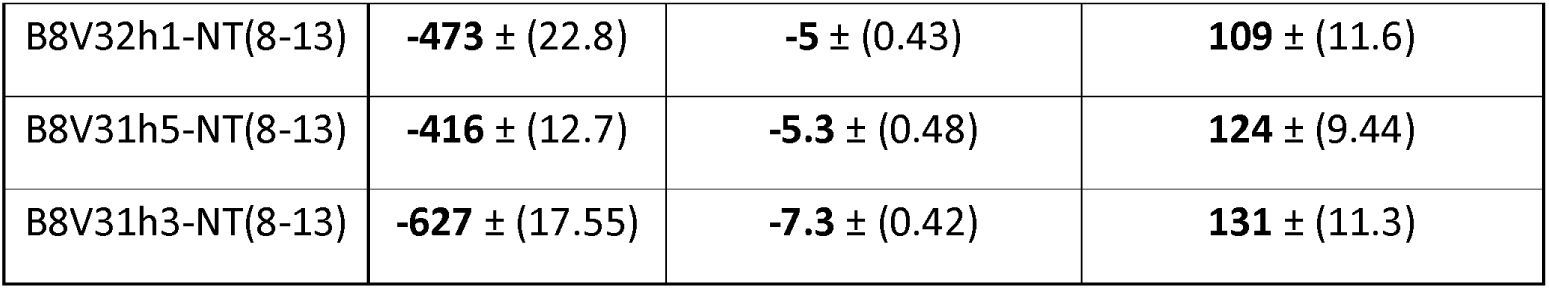
Hypothermia data. Table summarizing the main parameters recorded in BhTfR-mice injected with VHH-NT(8-13) variants at 500 nmol / kg. Maximal HT and HT duration were calculated as follows: Maximal HT = temperature prior to the injection (basal temperature) minus the temperature at the peak of the HT. Hypothermia duration = time during which the mouse body temperature is below basal temperature minus 2°C. Areas under the curve (AUC), were determined with GraphPad Prism software 10.2.0. Data are presented as mean ± SD of n=4 / 23 mice.

|  | Hypothermia AUC | Maximal hypothermia (°C) | Hypothermia duration (min) |
| --- | --- | --- | --- |
| C5neg-NT(8-13) | 0 ± (4.64) | -0.53 ± (0.12) | 3 ± (1.67) |
| C5-NT(8-13) | 73 ± (8.9) | -1.3 ± (0.36) | 22.5 ± (22.5) |
| B8V34-NT(8-13) | -236 ± (24.95) | -2.7 ± (0.41) | 63.8 ± (27.6) |
| B8V38-NT(8-13) | -197 ± (18.38) | -3.1 ± (0.41) | 84.4 ± (17) |
| C5V30-NT(8-13) | -470 ± (22.46) | -4.9 ± (0.53) | 109 ± (14.9) |
| B8V31-NT(8-13) | -237 ± (12.63) | -3.5 ± (0.32) | 84 ± (10) |
| B8V36-NT(8-13) | -353 ± (6.3) | -3.9 ± (0.065) | 97.5 ± (7.5) |
| B8V40-NT(8-13) | -434 ± (19.98) | -4.7 ± (0.46) | 128 ± (12) |
| B8V33-NT(8-13) | -149 ± (20.2) | -2.9 ± (0.49) | 45 ± (18.4) |
| B8V37-NT(8-13) | -414 ± (40.65) | -5.7 ± (0.77) | 150 ± (21.2) |
| B8V32h1-NT(8-13) | **-473** ± (22.8) | **-5** ± (0.43) | **109** ± (11.6) |
| B8V31h5-NT(8-13) | **-416** ± (12.7) | **-5.3** ± (0.48) | **124** ± (9.44) |
| B8V31h3-NT(8-13) | **-627** ± (17.55) | **-7.3** ± (0.42) | **131** ± (11.3) |

## Discussion

### Generation of VHHs with cross species reactivity that target the TfR1

Several technologies have been investigated to improve drug delivery to the CNS following peripheral administration, especially by hijacking the cellular machinery and using RMT processes (Pardridge, Boado et al. 1992, Jones and Shusta 2007, Fang, Zou et al. 2017). Foremost amongst the receptors involved in RMT is the TfR1 and our objective was to generate VHHs with cross species reactivity, that bind the mouse, rhesus monkey and human hTfR1. We used phage display technology on an immunized llama VHH library, and cross-selections on both the mouse and human TfR1. Following selection strategies performed on CHO cells expressing these receptors, we isolated several VHHs among which the C5, B8 and H3 that were shown to bind bound the human, mouse, and rhesus monkey TfR1, and the B6 that was only specific for human TfR1. Considering that the CDR3 loop is principally involved in antigen recognition (Inoue, Suganami et al. 2013, Muyldermans and Smider 2016), the three cross-specific VHHs belonged to the same VHH family, while VHH B6 belonged to another family. Thus, it appeared that the C5, B8 and H3 VHHs targeted a region conserved in the TfR1 of different species while B6 targeted a human specific TfR1 epitope. Our results suggested that our alternating boosting strategy led to an immune response restricted to two main TfR1 epitopes.

TfR1 ligands studied as shuttles for brain delivery were initially specific for rodent TfR1. These included the OX26 antibody that targets the rat TfR1 (Pardridge, Buciak et al. 1991, Moos and Morgan 2001, Haqqani, Thom et al. 2018, Thom, Burrell et al. 2018), and the 8D3 antibody or Fab (Zhang and Pardridge 2005, Boado, Zhang et al. 2009, Zhou, Boado et al. 2010, Manich, Cabezón et al. 2013, Pardridge 2015, Hultqvist, Syvanen et al. 2017, Sehlin, Fang et al. 2017), the R17-217 antibody or Nb62 nanobody (Lee, Engelhardt et al. 2000, Ulbrich, Hekmatara et al. 2009, Pardridge 2015, Wouters, Jaspers et al. 2020) that were specific for mouse TfR1. Binders selective for the cynomolgus monkey (cyno) and human TfR1 were also described (Yu, Atwal et al. 2014, Kariolis, Wells et al. 2020, Edavettal, Cejudo-Martin et al. 2022, Rué, Jaspers et al. 2023). However, these ligands do not bind the mouse TfR1 and can only be studied in transgenic mice expressing the human TfR1, hampering straightforward translation of pre-clinical studies to clinical phases. TfR1 binders with mouse-human cross species reactivity have also been developed more recently (Faresjö, Sjöström et al. 2024). Selecting such binders has proven difficult due to limited sequence identity (66 % between the human and mouse TfR1 ectodomains), and to the fact that human and mouse TfR1 extracellular domains share only 53 % homology when the Tf binding interface is excluded (Haqqani, Bélanger et al. 2024). Shark small single-domain antibodies (VNARs) cross-reactive against mouse, cyno and human TfR1 (Stocki, Szary et al. 2020, Stocki, Szary et al. 2021, Stocki, Szary et al. 2023) have been described. Unlike camelid VHH, VNARs are evolutionarily distant (Iezzi, Policastro et al. 2018) from human IgG sequences and could be difficult to humanize, thus potentially limiting their use in humans. To our knowledge, apart from VNAR TfR1 binders (Stocki, Szary et al. 2020), the VHHs we generated with well-established rodent, monkey and human cross species reactivity are unique.

### C5 and B8 VHHs target a unique epitope in the TfR1

The functional C5 and B8 binding epitopes did not overlap with the binding domain of H-Ft (Montemiglio, Testi et al. 2019), Tf (Cheng, Zak et al. 2004) or Human homeostatic iron regulator protein (HFE) (Lebrón, Bennett et al. 1998). Indeed, the C5, B8 and H3 VHH family targets an original region spanning the TfR1 homodimer interface between the helical and the apical domain that, to our knowledge, has never been targeted before. The structural symmetry of the TfR1 homodimer suggests that 2 VHH can bind simultaneously to the apical-helical domain interface. Besides the VHH95 recently described by Zhou et al. (Zhou, Bi et al. 2025), most of the TfR1 binders developed as brain shuttles target the TfR1 apical domain (Niewoehner, Bohrmann et al. 2014, Kariolis, Wells et al. 2020, Wouters, Jaspers et al. 2020, Rué, Jaspers et al. 2023). When binding to the external face of the TfR1 apical domain, these binders may compete with the binding of the H-Ft ligand and interfere with TfR1 function. When compared to human TfR1, the C5/B8 targeted region shares 95.8 % and 83.3 % sequence similarity with the rhesus monkey and mouse TfR1, respectively, while for the apical domain, the similarity percentage with the mouse TfR1 is slightly lower (below 80 %). Similarities, but also divergences in the primary sequences and structures between mouse, rhesus monkey and human TfR1 are probably key to the success of our cross-species selection strategy, but also to the differences in VHH affinities and binding kinetic parameters between the TfR1 from the 3 species. Indeed, B8 and C5 were found to have a K_D_ of approximately 30 and 110 nM for the mouse TfR1, respectively. These affinities fall within a range suitable for efficient transcytosis across the BBB in mice (Yu, Zhang et al. 2011, Bien-Ly, Yu et al. 2014, Thom, Burrell et al. 2018, Do, Capdevila et al. 2020). However, due to their sub-nanomolar affinity for the human TfR1, they were not in the optimal affinity window for achieving efficient transcytosis in humans. Nevertheless, the question of the optimal affinity for crossing the BBB and optimal molecular architecture of the TfR1-targeting brain shuttles has been mostly established with apical domain binders and remains a matter of debate (Arguello, Mahon et al. 2022, Pardridge 2023).

### VHH fusion to a human IgG1 Fc region demonstrates potency for brain delivery

To determine whether our TfR1 binders to the apical-helical domain interface could be developed as brain shuttles, and to select the most adapted TfR1 binding valency for CNS drug delivery, a human IgG1 Fc domain used as a prototypic protein cargo with partial mAb features was successfully fused to the C-terminus of the C5 and B8 VHHs. Both bivalent and monovalent VHH-hFc fusions bound to TfR1. However, the bivalent C5- and B8-hFc fusions showed respectively a 30- to 4-fold higher apparent affinity to cellular mouse TfR1 compared to the monovalent form, consistent with an avidity effect. Despite their lower plasma exposure due to TMDD by comparison with the negative control C5neg-hFc fusions, both the bivalent and monovalent B8 and C5-hFc formats exhibited similar brain-to-plasma ratio percentages of approximately 5.5 % and 6.5 % for the monovalent and bivalent binders, respectively (Figure 6F). These ratios are advantageous when compared to the brain-to-serum ratio of 0.1 % – 1 % reported for monoclonal antibodies in mouse brain post-systemic injection (Chang, Wu et al. 2021, Van De Vyver, Walz et al. 2022). Interestingly, the (VHH_TfR1_)_1_-hFc reached greater brain concentrations than (VHH_TfR1_)_2_-hFc in line with the better brain delivery potential of monovalent TfR1 binders previously described (Yu, Zhang et al. 2011, Bien-Ly, Yu et al. 2014, Yu, Atwal et al. 2014, Kariolis, Wells et al. 2020, Arguello, Mahon et al. 2022). However, whole brain accumulation of TfR1 binders does not necessarily imply effective crossing of the BBB considering that a too strong affinity, or an avidity effect may cause trapping of conjugates in brain capillary endothelial cells, at the expense of perivascular and parenchymal distribution (Yu, Zhang et al. 2011, Bien-Ly, Yu et al. 2014). We did use capillary depletion to assess the eventual BBB retention of our conjugates and their accumulation in post-vascular brain compartments (data not shown), but such approaches tend to overestimate the concentration of TfR1 binders in brain extracellular fluid compartment, in comparison for example with micro-dialysis (Chang, Wu et al. 2021). To assess the BBB crossing efficiency of a brain shuttle, brain imaging techniques have also proven valuable. Recently, Faresjö *et al*. utilized one of our patented VHH_TfR1_ (Cohen, David et al. 2019) to evaluate novel TfR1 targeting VHHs they had identified by brain autoradiography (Faresjö, Sjöström et al. 2024). To further demonstrate the BBB crossing potential of our VHH in the monovalent TfR1 binding format that gave the best absolute brain exposure, we implemented an additional readout based on a physiological brain response to the NT(8-13) peptide as an alternative cargo.

### Induction of HT by VHH-NT(8-13) fusions in C57BL/6 and B-hTfR mice after i.v. administration

NT conjugated or fused to Low-Density Lipoprotein or LDL related protein 1 Receptor binders has been successfully used by different groups (Demeule, Beaudet et al. 2014), including ours (Ferhat, Soussi et al. 2025) to induce central analgesia or HT following systemic injection. Others have induced HT using TfR1 targeting VHH-NT fusions (Stocki, Szary et al. 2020, Wouters, Jaspers et al. 2020, Wouters, Jaspers et al. 2022, Stocki, Szary et al. 2023) to demonstrate: i) the transcytosis capacity via a TfR1 dependent RMT process and ii), once the BBB is crossed, the target engagement of the NT peptide with its NTSR1 and/or NTSR2 receptors expressed in neurons and astrocytes in the hypothalamus and other brain structures, respectively (Tabarean 2020, Kyriatzis, Bernard et al. 2021). Using a low molecular weight VHH-NT(8-13) format, we demonstrated the capacity of both C5 and B8 parental VHHs to efficiently deliver their peptide cargo into the CNS in pharmacologically active amounts. This delivery resulted in a decrease in WT mouse body temperature following intravenous injection at the dose of 500 nmol / kg. TfR1-dependent capacity of our VHH to induce central HT raised the following issues: first, our VHH-NT have a basic pI, and it has been shown that VHH with such features crossed the BBB via absorptive-mediated transport (Li, Bourgeois et al. 2012, Li, Vandesquille et al. 2016). Second, the capillaries in the circumventricular organs of the hypothalamus are fenestrated enabling the parenchymal access of molecules with a molecular weight below 10 kDa (Cheunsuang, Stewart et al. 2006, Miyata 2022, Su, Esparza et al. 2022). The inclusion in our study of the C5neg-NT(8-13) control, which lacks the ability to bind to mouse TfR1 but shares similar physicochemical characteristics with C5—such as a basic pI (8.99) and a molecular weight of approximately 15 kDa—and does not induce HT, confirmed that the pharmacological response observed with B8 and C5-NT(8-13) was linked to a TfR1-dependent RMT across brain endothelial cells. This transport mechanism led to subsequent target engagement of NT receptors. To address the limited plasma half-life of VHH (Harmsen and De Haard 2007), we also studied a high molecular weight (VHH)_1_-hFc-NT(8-13) format, which presumably decreases size related hypothalamus access, renal elimination but also leverages the neonatal Fc receptor (FcRn) recycling mechanism associated with the hFc moiety to prolong the blood half-life of VHH-NT(8-13). Notably, the dose required to elicit a significant HT was reduced 50-fold compared to the low molecular weight VHH-NT(8-13) fusion and by more than 500-fold when compared to the dose of NT peptide (i.e.: 8 mg / kg or 4.7 µmol / kg) that initiates non-significant HT in mice (Ferhat, Soussi et al. 2025). Moreover, the duration of this effect improved in a dose-dependent manner. The gradual return to baseline presumably results from the desensitization of the NTSR1 receptor (Demeule, Beaudet et al. 2014) and/or from the reduction in plasma concentration of the (VHH)_1_-hFc-NT(8-13) fusion due to TMDD. The (C5)_1_-hFc-NT(8-13) fusion, which exhibits low mouse TfR1 binding potential, demonstrated a stronger and more prolonged effect compared to the (B8)_1_-hFc-NT(8-13) fusion, which has a 20-fold higher mouse TfR1 binding potential. This enhanced HT observed with (C5)_1_-hFc-NT(8-13) may be due to one or more of the following factors: i) reduced TMDD, facilitating greater and longer exposure at the BBB; ii) improved capacity of (C5)_1_-hFc-NT(8-13) for RMT; and iii) higher engagement with NTSRs relative to parenchymal TfR1, due to a favorable affinity balance for NTSRs. Using a similar Fc protraction strategy and NT peptide fusion, Stocki et al. previously demonstrated the brain delivery potential of the TXB2 VNAR that targets TfR1 and its optimized version TXB4 (Stocki, Szary et al. 2021, Stocki, Szary et al. 2023). In these studies, the homodimeric (TXB4)_2_-Fc-(NT)_2_ format showed a maximal −2.9 °C decrease in mice body temperature at 120 min at the dose of 25 nmol / kg. Interestingly, (C5)_1_-hFc-NT(8-13) demonstrated a similar maximal decrease in mice body temperature (−3.1 °C) at the same time post-injection but at a 2.5 lower dose (10 nmol / kg). Although C5 exhibited mouse TfR1-binding properties leading to efficient RMT of NT across the wild-type mouse BBB, this was not observed in B-hTfR mice where C5-NT(8-13) failed to induce HT, presumably due to elevated affinity, underscoring the need for additional optimized trans-species binders.

### Engineering and optimization of VHH binding parameters on TfR1

Antibodies and antibody fragments have been engineered to adjust TfR1 binding affinities and kinetics to identify variants and structural features optimal for BBB transcytosis (Yu, Zhang et al. 2011, Karaoglu Hanzatian, Schwartz et al. 2018, Do, Capdevila et al. 2020, Chang, Wu et al. 2021, Esparza, Su et al. 2023, Stocki, Szary et al. 2023). Following similar methodologies, we developed affinity variants of our parental TfR1 binders. Using NT-induced HT as a functional readout, we found that a reduction of at least two logs in parental VHH affinity and a two-log increase in dissociation rate were necessary for effective HT in B-hTfR mice. Additionally, we observed a correlation between K_D_, k_off_, and the ability to engage NTSRs after crossing brain endothelial cells. Edavettal et al. corroborated the importance of the k_off_ as a key kinetic parameter for successful BBB transcytosis and determined an optimal human TfR1 affinity range of approximately 6-10 nM to maximize brain uptake of their Transcytosis Enabling Module (TEM) (Edavettal, Cejudo-Martin et al. 2022). In our studies, affinity variants of B8-NT(8-13) and C5-NT(8-13) that significantly induced HT displayed lower affinities, around 20 to 220 nM, in line with the affinity range of the Brain Shuttle, which has a human TfR1 affinity of 130 nM (Grimm, Schumacher et al. 2023). Notably, the humanized B8V31h3-NT(8-13), our most potent HT-inducing human TfR1 binder exhibited an affinity for human TfR1 of 210.8 nM and the fastest dissociation rate reported in the literature (e.g., ∼0.2 s⁻¹). Conversely, Stocki et al. identified a positive correlation between the association rate of their engineered VNARs and brain uptake efficiency, despite the sub-nanomolar mouse TfR1 affinities of these binders (Stocki et al., 2023). This highlights that optimizing the TfR1-binding parameters of a BBB-crossing agent requires tailoring the specific characteristics of each agent and is likely influenced by the targeted epitopes. Through optimization of the C5 and B8 parental VHH, we identified key positions (e.g., F30K and A54S, IMGT numbering) that enhance human-rhesus monkey TfR1 cross-reactivity while reducing affinity to a range better suited for efficient RMT. These modifications generate ideal variant candidates for preclinical studies in transgenic mice expressing human TfR1 and in monkeys. Additionally, while generating surrogate binders for studies in WT mice appeared challenging (Sonoda, Morimoto et al. 2018), we demonstrated that the C5 and B8 VHHs exhibit mouse TfR1 binding parameters similar to those of certain variants on human-rhesus monkey TfR1, suggesting their suitability for preclinical studies in WT or disease mouse models.

## Conclusion

We selected and characterized cross-species reactive VHHs that target the TfR1 without competing with endogenous TfR1 ligands. These VHHs were engineered to modulate their TfR1 binding parameters and screened for optimal crossing of the BBB via a TfR1-dependent mechanism. Our VHHs targeted a novel and unique epitope on TfR1 and promoted the delivery of various payloads— such as an IgG1 Fc fragment and a neuropeptide—into the brain parenchyma. Pharmacokinetic studies in mice further confirm that monovalent VHH-Fc formats achieve superior brain exposure compared to bivalent constructs, likely due to reduced receptor-mediated clearance and improved transcytosis efficiency. Importantly, we establish a correlation between⍰TfR1 binding kinetics⍰(especially dissociation rate) and neurotensin peptide pharmacodynamics, identifying an optimal affinity window for efficient BBB transcytosis. Our VHHs represent a⍰versatile and modular platform⍰for CNS drug delivery and will contribute to the development of⍰next-generation brain-targeted biotherapeutics including enzyme replacement therapies, antibody-based treatments for neurodegenerative diseases, and oligonucleotide delivery for CNS disorders.

## Materials and Methods

### Animals

Procedures involving animals conform to National and European regulations (EU Directive N°2010/63) and to authorizations delivered to our animal facility (N° J 13 055 05) and to the projects N°35462-2022021518324714 v2 and N°40739-2023020210575425 v3 by the French Ministry of Research and Local Ethics Committee (CE N°014). All efforts were made to reduce the number of animals and to minimize animal suffering. We used 8–14 week-old C57Bl/6 wild type (WT) male mice (Janvier-labs, Le Genest-Saint-Isle, France) and B-hTfR male mice (Biocytogen, Beijing, China). Llama immunizations (outsourced) were performed by a professional veterinarian and were executed in strict accordance with good animal practices.

### Cloning, establishment of cell lines

#### Construction of the CHO cell lines expressing the mouse, human and rhesus monkey TfR1 receptors in fusion with EGFP

The plasmid coding for mouse TfR1 fused to EGFP was purchased from GeneCopoeia (Rockville, USA) and was named pmTfR1-EGFP. The full-length coding region for human TfR1 was generated by reverse transcription-PCR (RT-PCR) from human brain total RNA (Clontech, Mountain View, USA) using the following forward and reverse primers: ATATATGAATTCGGCTCGGGACGGAGGACGC and TTAATTGTCGACAGAACTCATTGTCCCAACCGTCAC, respectively. The phTfR1-EGFP plasmid construct was generated by cloning the human TfR1 cDNA in frame with EGFP into the pEGFP-C1 vector (Clontech). The plasmid coding for rhesus monkey TfR1 fused to EGFP was obtained by gene synthesis (GeneCust, Boynes, France) and cloned into the pEGFP-C1 vector and was named prhTfR1-EGFP. All the constructs were fully sequenced and used to transfect Chinese Hamster Ovary cells (CHO-K1, ATCC number CCL-61™, Manassas, USA) using jetPei™ (Polyplus Transfection, Illkirch-Graffenstaden, France) according to the manufacturer’s instructions. Forty-eight hours after transfection, 800 μg/ml geneticin (Thermo Fisher Scientific, Waltham, USA) was added to the growth medium as selection agent. One week after the beginning of the selection, individual fluorescent cells were seeded in 96 well plates. Isolated clones expressing EGFP fused to the N-termini of mouse, human, or rhesus monkey TfR1 were selected, amplified, and validated for the expression of the fusion proteins of interest by western blotting and immunocytochemistry. The validated stable cell lines were named CHO-mTfR1-EGFP, CHO-hTfR1-EGFP, and CHO-rhTfR1-EGFP and were cultured at 37 °C in a humidified atmosphere, with 5 % CO_2_ in Nutrient Mix/F-12 (Ham) GlutaMAX™ medium complemented with fetal calf serum 10 % (v / v), 100 µg / ml streptomycin, 100 U / ml penicillin and 400 µg / ml geneticin (all from Thermo Fisher Scientific). CHO WT cells were cultured in the same conditions, without geneticin.

#### Preparation of cell membrane extracts for llama immunization

Total membranes were prepared from the stable cell lines expressing mouse or human TfR1 using the following protocol. Cells were washed with PBS, incubated with PBS EDTA 5 mM, scrapped and centrifuged at 300 g for 5 min. The cell pellets were frozen at −80 °C before resuspension in lysis buffer (Tris 10 mM, EDTA 1 mM, MgCl_2_ 5 mM), and homogenized in a 7 ml Dounce homogenizer by 10 up and down strokes with each of 2 pestles of different clearances, 71 μm followed by 20 μm. After a quick sonication step, the suspensions were centrifuged 15 min at 1700 g to remove cellular debris, the supernatants were conserved, and the pellets were resuspended in 1 ml of lysis buffer before centrifugation 15 min at 1700 g. Both supernatants were pooled and centrifuged 50 min at 100 000 g, 4 °C. The pellets containing total membranes were finally resuspended in PBS and stored at −80 °C until llama immunization.

#### Llama immunization and library construction

A llama (*Lama glama*) was immunized subcutaneously 4 times, at days 0, 9, 18 and 28 with membrane preparations from CHO stable cell lines expressing the receptors of interest. The 1^st^ and 2^nd^ injections were performed with membrane preparations from the CHO-hTfR1-EGFP cell line, while the 3^rd^ and the 4^th^ injections were performed with membrane preparations from both CHO-mTfR1-EGFP and CHO-hTfR1-EGFP cell lines. The VHH library was constructed as previously described (Alvarez-Rueda, Behar et al. 2007, Behar, Chames et al. 2009). Briefly, mRNAs coding for VHHs were amplified by RT-PCR from the total RNA of peripheral blood mononuclear cells isolated by ficoll gradient and cloned into the pHEN1 phagemid.

#### Selection of TfR1 targeting VHHs by phage display

To produce phage-VHHs, 50 µl of the bacterial library was grown in 50 ml of 2xYT medium supplemented with 100 µg / ml ampicillin (2YTA) at 37 °C with shaking (250 rpm) to an OD_600_ comprised between 0.4 and 0.6. The culture was infected with KM13 helper phage using a multiplicity of infection of 20, for 30 min at 37 °C without shaking. The culture was centrifuged for 15 min at 3000 g. The bacterial pellet was resuspended in 250 ml of 2YTA supplemented with 50 µg / ml kanamycin and incubated overnight at 30 °C with shaking (250 rpm). The overnight culture was split in 10 vials and centrifuged for 20 min at 3000 g. Five ml of PEG 8000 20 % / NaCl 2.5 mM were added to the supernatant in a new vial and incubated for 1 h on ice to precipitate phage particles. The solution was centrifuged for 15 min at 3000 g at 4 °C, and the phage-containing pellet was resuspended with 1 ml of PBS. Another centrifugation step (2 min, 16000 g, 4 °C) was performed to eliminate bacterial contaminants, and 200 μl of PEG 8000 20 % / NaCl 2.5 mM were added to the supernatants in a new vial. After 30 min on ice and a last centrifugation (5 min, 16000 g, 4 °C), phage-containing pellets were resuspended in 1 ml of PBS/ bovine serum albumin 2 % (w / v, BSA).

Different phage panning strategies were performed on CHO cell lines expressing the receptors of interest fused to EGFP (20-50 × 10^6^ cells). Cells were washed 3 times in PBS using centrifugation steps (3 min at 700 g at 4 °C), and phage-VHH library (1 ml) and cells were saturated in PBS/BSA 2 %, for 1 h at 4 °C with rotation. Next, the phage-VHH library was incubated with irrelevant, non-transfected CHO-TRVb cells (cell line that does not express functional endogenous hamster TfR1) (depletion step), while cells were incubated with 100 µl of 100 µM purified anti-cell VHHs (masking) (Even-Desrumeaux, Nevoltris et al. 2014), for 1.5-2 h at 4 °C, with rotation. The phage-VHH library was recovered, added to cells in the presence of the mask, and incubated for 1.5-2 h at 4 °C with rotation. Cells were washed 10 times with 1 ml of PBS using centrifugation (3 min at 700 g at 4 °C). Bound phages were eluted with 500 µl of trypsin solution (Sigma-Aldrich) at 1 mg / ml during 30 min at RT with rotation. Elution was stopped by the addition of 500 µl PBS / BSA 2 % and the solution was centrifuged to eliminate cellular debris. For titration, serial dilutions were performed starting from 5 µl of eluted phage. Phage dilutions were incubated without shaking with log-phase TG1 cells for 30 min at 37 °C, plated on 2YTA/glucose 2 % in Petri dishes, and incubated overnight at 37 °C. The remaining 995 µl of eluted phage were incubated without shaking with log-phase TG1 cells for 30 min at 37 °C, plated on 2YTA/glucose 2 % in Petri dishes, and incubated overnight at 30 °C. Colonies were harvested from the plates incubated at 30 °C, suspended in 2YTA/glucose 2 % and used for phage production as previously described, for the next round of selection. Two rounds of selection were systematically performed. At the end of the second round, colonies isolated from titration were picked and grown overnight at 37 °C in 2 different 96-deep-well plates in 450 µl of 2YTA/glucose 2 % and stored at −80 °C after the addition of 15 % glycerol (masterplates).

#### Monoclonal VHH production for screening

A 96-deep-well plate replicator was used to replicate the masterplate, and 15 µl of output from selection round 2 were inoculated to 435 µl of 2YTA. Cells were grown at 37 °C (250 rpm) until the OD_600_ reached 0.5. VHH expression was induced by the addition of 100 µM IPTG (isopropyl β-D-thiogalactopyranoside), overnight at 30 °C (250 rpm). After centrifugation 10 min at 3000 g, 4 °C, VHH-containing supernatants were tested for binding by flow cytometry.

#### VHH screening, apparent affinity & competition studies using cell-based assays

Flow cytometry experiments were performed in 96 well plates, using the antibodies referenced in Supplemental Table S1. All cell washes or change of buffer were done by centrifugation for 3 min at 700 g at 4 °C. For VHH screening, CHO cell lines expressing the TfR1 fused to EGFP or CHO-TRVb cells at 2-3 × 10^5^ cells / well were saturated by PBS/BSA 2 % solution for 30 min to avoid nonspecific binding. After centrifugation, 75 μl of VHH-containing supernatant was added to 75 μl of cells in PBS/BSA 2 % and incubated for 1 h at 4 °C. After 3 washes in PBS/BSA 2 %, cells were incubated for 1 h with primary antibody, washed 3 times with PBS/BSA 2 %, and incubated for 45 min with secondary antibody. After 3 last washes in PBS/BSA 2 % and cell resuspension in PBS, fluorescence was measured using a MACSQuant® cytometer (Miltenyi, Bergisch Gladbach, Germany). For binding experiments, 50 µl of purified VHH or VHH-Fc in dilution series at concentrations ranging from 50 µM to 1 pM were incubated on cells at 2-3 × 10^5^ cells / well for 1 h at 4 °C. The following steps were identical to the screening steps. In some experiments, after the 3 last washes, cells were resuspended in PBS and 4 % paraformaldehyde (PFA, Sigma Aldrich, Darmstadt Germany) (v/v 1:1) for 15 min at RT to fix cells. PFA was removed by centrifugation, cells were resuspended with PBS and fluorescence was measured using a MACSQuant® cytometer (Miltenyi) or an Attune^TM^ NxT flow cytometer equipped with Attune™ Cytometric software v5.2.0 (Thermo Fisher Scientific). Binding curves were fitted with GraphPad Prism® software (version 10.2.0) using a nonlinear fit to get the K_d/app_ or EC90 values.

For competition experiments, cells at 2-3 × 10^5^ cells / well were saturated by PBS/BSA 2 % solution for 30 min at 4 °C. After centrifugation, cells were incubated for 1 h with 50 µl of competitors in dilution series, followed by the addition of 50 µl of tracers at EC90 and incubation for 1 h. The following steps were identical to the screening steps. The number of positive cells in each assay was standardized to 10 000. Results were expressed in arbitrary units corresponding to the fluorescence ratio normalized with the EGFP signal (receptor expression).

To determine apparent inhibition constants (K_i/app_), Expi293^TM^ cells (Thermo Fisher Scientific) expressing the mNTSR1-GFP receptor obtained post-transient transfection of the plasmid encoding mNTSR1-GFP generated by gene synthesis (GeneCust) were incubated at a density of 1.10^6^ cells / ml, with increasing concentrations of tested compounds in the presence of the reference AlexaFluor®680-RRPYIL-OH fluorescent compound (A680-NT(8-13)) at a sub-saturating concentration of 30 nM for 1 h at 37 °C. A680-NT(8-13) was synthesized by solid phase peptide synthesis (SPPS) based on Fmoc strategy using a Liberty (CEM^®^) microwave synthesizer (CEM Corporation, Charlotte, USA). After treatment, cells were transferred in 96-deep-well plates containing 1 % SVF 0.02 % sodium azide and 5 mM EDTA in PBS and centrifuged. After removing supernatants, living versus dead cells were discriminated using viability staining (1/2500 LIVE/DEAD Violet (Thermo Fisher Scientific)) for 30 min at 4 °C. The excess of marker was removed by centrifugation and cells were resuspended in 5 mM EDTA in D-PBS and 4 % PFA (v/v 1:1) was added into wells to fix cells for 15 min at RT. PFA was removed by centrifugation and cells were resuspended with 5 mM EDTA in D-PBS. The cellular A680-associated fluorescent signals were quantified using an Attune^TM^ NxT flow cytometer and Inhibition curves were fitted with GraphPad Prism^®^ software using a nonlinear fit to get the K_i/app_ values. Flow cytometry experimental procedure, acquisition and analysis were performed following MIFlowCyt guidance (Lee, Spidlen et al. 2008).

#### Immunocytochemistry

After 30 min cell saturation with cell culture medium supplemented with low endotoxin BSA 1 %, cells alone or incubated 1 h at 37 °C with VHH at 50 µg / ml, or VHH-Fc at 50 nM, were washed and subsequently fixed with PFA 4% for 10 min at RT, washed again and processed for immunocytochemistry. Coverslips were incubated for 30 min at RT in saturation/permeabilization PBS buffer containing BSA 3 % with or without triton X-100 0.1 % (Sigma-Aldrich). After 3 washes with PBS, cells were incubated for 1 h at RT with primary antibodies directed against cMyc (mouse, 1/1000, Thermo Fisher Scientific), or Fc (Alexa594-conjugated, 1/1000, Jackson ImmunoResearch, Cambridge, UK). Following 3 washes with PBS, cells were incubated with Alexa594-conjugated anti-mouse secondary antibodies (1 / 800, Jackson ImmunoResearch), together with Hoechst#33258 at 0.5 µg / ml for nuclei staining (Thermo Fisher Scientific). After 3 last washes with PBS, coverslips were mounted in ProLong® Gold Antifade reagent (Thermo Fisher Scientific) and were analyzed using a LSM700 (Zeiss) confocal microscope with Zen 2012 software. Images were obtained using a 63× Plan-Apochromate oil immersion objective.

#### Culture, transfection, and cytometry analysis of human TfR1 mutants in HEK cells

Mammalian expression plasmid phTfR1-EGFP was used to express GFP-human TfR1 in FreeStyle^TM^ HEK293 cells (Thermo Fisher Scientific). Human TfR1 mutants were generated through SOE PCR (Splicing by Overlap Extension Polymerase Chain Reaction) and Gibson cloning (Thermo Fisher Scientific) between EcoRI and SalI restriction sites. On the day preceding transfection, cells were subcultured to achieve a density of 2–3.10^6^ cells / ml on the following day. On the day of the transfection, the cells were subjected to a 5 min centrifugation at 100 g and subsequently resuspended in a fresh FreeStyle HEK293 Expression medium (Thermo Fisher Scientific) to achieve a density of 2.5×10^6^ cells / ml. For a 2 ml transfection volume, the cells were incubated with 3 µg of plasmid at a concentration of 0.5 µg / µl for five minutes at 37 °C, 180 rpm and 8 % CO₂. 36 µl of polyethylenimine (PEI) at a concentration of 0.5 mg / ml was added to the cells. Subsequently, the cells were incubated for 48 h at 37 °C with agitation at 180 rpm and 8 % CO₂ to allow human TfR1 expression. After 48 h, approximately 100 µl of the transfected cells were centrifuged at 100 g and washed with 1 ml PBS + 0.1 % BSA (PBSF). Cells were then resuspended in 50 µl of PBSF containing 100 nM of either biotinylated VHH C5 or biotinylated VHH B8 and 50 nM Transferrin-AF647 (Thermo Fisher Scientific) and incubated at 20 °C for 2 h under agitation. Cells were washed twice using ice-cold PBSF and labeled with Streptavidin-PE (Thermo Fisher Scientific) 1:100 in 50 µl on ice for 15 min. Cells were washed, resuspended in PBS and analyzed with BD FACS Aria™ III cytometer. Cytometer lasers gains were set so that non-transfected cells (endogenous expression) resulted in double negative fluorescence signal on both APC and PE channels.

### Cloning and production of recombinant proteins

#### Cloning of the VHH in phase with a human IgG1 Fc region as homodimers or heterodimers Homodimeric VHH-hFc

cDNAs encoding the C5, B8 and D12 VHHs were amplified by PCR and cloned into the pINFUSE-IgG1-Fc2 vector (InvivoGen, Toulouse, France) to encode a human IgG1 crystallizable fragment (hFc) fused in C-ter of the VHH. The plasmid constructs were fully sequenced and named (C5)_2_-hFc*, (B8)_2_-hFc* and (D12)_2_-hFc*, the (*) referring to the presence of effector functions. To reduce effector functions, DNA fragments encoding C5 and B8 VHH-hFc with LALA mutations (L234A/L235A numbering according to EU nomenclature) were obtained by gene synthesis and cloned in pCDNA3.4 (Thermo Fisher Scientific). A C5 negative control VHH called C5neg encompassing 2 Ala mutations in the CDR3 of C5 was cloned in the pcDNA3.4 plasmid. The (C5)_2_-hFc_LALA_, (B8)_2_-hFc_LALA_ and (C5neg)_2_-hFc_LALA_ plasmid constructs were fully sequenced and were referred to as (C5)_2_-hFc, (B8)_2_-hFc and (C5neg)_2_-hFc in the figures and in the text.

#### Heterodimeric VHH-hFc

cDNAs encoding C5, B8 and C5neg VHH-hFc with LALA mutations and knob mutation T366W (numbering according to EU nomenclature) were obtained by gene synthesis and cloned in pcDNA3.4. The (C5)_1k_-hFc_LALA_, (B8)_1k_-hFc_LALA_ and (C5neg)_1k_-hFc_LALA_ plasmid constructs were fully sequenced and referred to as (C5)_1k_-hFc, (B8)_1k_-hFc, and (C5neg)_1k_-hFc in the figures and in the text. The cDNA fragment encoding hFc with LALA mutations and hole mutations T366S, L368A, Y407V (numbering according to EU nomenclature) was obtained by gene synthesis and was cloned in pcDNA3.4 and named hFc_LALA1h_.

### Low and high molecular weight VHH-NT(8-13) fusions

#### Low molecular weight VHH-NT(8-13)

cDNAs encoding the C5, B8, C5neg, and the C5 and B8 variant VHHs fused to a (His)_6_-(G_4_S)_2_ linker and the smaller active moiety of NT (of amino-acid sequence: RRPYIL) were obtained by gene synthesis and cloned in the pHEN1 phagemid.

#### High molecular weight VHH-NT(8-13)

cDNAs encoding hFc with LALA and hole mutations with -(G_4_S)_3_-RRPYIL fused to its C-Terminal end were obtained by gene synthesis, were cloned into pcDNA3.4, and named hFc_LALA-_NT(8-13)_1h_.

Cloning of mouse, human and rhesus monkey TfR1 ectodomains in fusion with a mouse IgG2a Fc cDNAs encoding the mouse, human and rhesus monkey TfR1 ectodomains (amino acids 89 to 763 from mouse TfR1 (NP_001344227.1), amino acids 89 to 760 of the human TfR1 (AAA61153.1) and rhesus monkey TfR1 (NP_001244232.1)) were obtained by gene synthesis (Genecust) and cloned in the pINFUSE-mIgG2b-Fc2 vector (InvivoGen) to encode a mouse IgG2a Fc (mFc) fused in C-terminus to the mouse, human and rhesus monkey TfR1 ectodomains. A sequence coding for a (G_4_S)_3_ flexible linker was inserted between the human or rhesus monkey TfR1 ectodomains and mFc. The plasmid constructs were fully sequenced and named pmFc-mTfR1, pmFc-hTfR1 and pmFc-rhTfR1.

### Production of fusion proteins

#### Production and Purification of VHH

For large scale VHH or VHH-NT(8-13) production, phagemids were transformed in *E. coli* BL21(DE3) strain. Transformed bacteria were grown in 200 ml of 2YTA until 0.5 < OD_600_ < 0.8 and induced with 100 μM IPTG for an overnight growth at 30 °C with shaking (250 rpm). Bacteria were pelleted and lysed by freeze-thawing and BugBuster® Protein Extraction Reagent (Novagen, Pretoria, South Africa). After a centrifugation step (4000 g, 20 min), VHH were purified from the supernatant using metal affinity chromatography, TALON® Superflow™ (Cytiva, Marlborough, USA), according to the manufacturer’s instructions. Bound molecules were eluted with 150 mM imidazole, which was eliminated using a PD-10 Desalting column (GE Healthcare, Chicago, USA) or by diafiltration on 10 kDa MWCO Amicon® Ultra centrifugal concentrator (Merck, Darmstadt, Germany). For VHH-NT(8-13) fusions, an additional gel filtration step using a Superdex 75 column (Cytiva) together with a Mustang E (Pall, Port Washington, USA) filtration was added to remove endotoxins to a level below 0.7 EU / mg. Endotoxin level was assessed using the Pierce Chromogenic Endotoxin Quant kit (Thermo Fisher Scientific) according to the manufacturer’s instructions. Purified VHH or VHH-NT were validated for their respectively SEC-UV and LC-UV-MS on a Orbitrap Exploris 240 (Thermo Fisher Scientific).

#### Production and Purification of VHH-hFc fusions

Fusion proteins ((VHH)_2_-hFc, (VHH)_2_-hFc_LALA_, (VHH)_1_-hFc_LALA_, and (VHH)_1_-hFc_LALA_-NT(8-13)) were produced using the Expi293^TM^ Expression System following transient transfection of 1 µg of total plasmid DNA per ml of transfection, according to the manufacturer’s instructions (Thermo Fisher Scientific). To produce heterodimeric (VHH)_1_-hFc_LALA_ or (VHH)_1_-hFc_LALA-_NT(8-13)_1_ a 1 to 4 transfection ratio was used for p(VHH)_1k_-hFc_LALA_ and phFc_LALA1h_ or phFc_LALA-_NT(8-13)_1h_ respectively. The cell suspension containing the fusion proteins was harvested 72 h post-transfection and centrifuged for 5 min at 300 g. The supernatant was then centrifuged for 15 min at 5000 g before filtration on nylon filter membranes with 0.22 µm pore size (Sigma-Aldrich) and purified on an Akta Pure protein purification system using first a protein A purification on a MabSelect™ PrismA resin (Cytiva) and next gel filtration on a Superdex 200 column. Purified fusions formulated in PBS buffer 1X pH 7.4 (Euromedex, Souffelweyersheim, France) were validated for their purity and identity by respectively SEC-UV and LC-UV-MS.

#### Production and Purification of mouse Fc-TfR1 ectodomain fusion

mFc-mouse TfR1 (mFc-mTfR1), mFc-human TfR1 (mFc-hTfR1) and mFc-rhesus monkey TfR1 (mFc-rhTfR1) ectodomains were produced by transient expression in Expi293^TM^ (Thermo Fisher Scientific) using 1 µg of plasmid DNA per ml of transfection. Seventy-two hours post-transfection, the clarified supernatant was purified using protein G on a HiTrap™ Protein G HP resin (Cytiva). The purified protein was next formulated in PBS 1X pH 7.4 (Euromedex) by buffer exchange using a centrifugal concentrator with a molecular cut-off of 50 kDa MWCO Amicon® Ultra centrifugal concentrator (Merck, Darmstadt, Germany).

#### Analysis of VHH and VHH-Fc fusions by SEC-UV and LC-UV-MS

For SEC-UV analysis, 3 µg of VHH or VHH-Fc were injected into an AdvanceBio SEC column (7.8 I.D. × 300 mm, 2.7 µm particles; Agilent Technologies, Santa Clara, USA). Sodium phosphate 150 mM was used as the mobile phase with an isocratic flow rate of 1.2 ml / min. The signal was monitored at A280 and A214 nm. Data processing was performed using Thermo Chromeleon 7.2.10 software. For LC-UV-MS analysis, a Vanquish™ Flex system coupled with an Orbitrap Exploris 240 mass spectrometer (Thermo Fisher Scientific) was used. VHH were analyzed with a bioZen 3.6 µm Intact XB-C8 (2.1 I.D. × 150 mm) column, and VHH-Fc with a bioZen 2.6 µm WidePore C4 (2.1 I.D. × 150 mm) column (Phenomenex, Le Pecq, France). A gradient from H₂O to acetonitrile with 0.1 % HCOOH was used. The signal was monitored at 280 nm and 214 nm, and the mass acquisition was performed in positive mode. Thermo Biopharma Finder 3.0 software allowed data deconvolution.

#### Affinity evaluation using SPR

SPR measurements were performed at 25 °C using a Biacore T200 apparatus (Cytiva) and 50 mM HEPES-NaOH pH 7.4, 150 mM NaCl 0.005 % Tween-20 (*v*/*v*) as running buffer. An anti-mouse antibody (Sigma Alrich) was first immobilized via amine coupling at a density of 100 fmol / mm^2^ onto a Biacore^TM^ CM5 sensorchip (Cytiva). Next, the mFc-mTfR1, mFc-hTfR1 and mFc-rhTfR1 ectodomain ligands were immobilized by affinity interaction at a density of 4 – 11.1 fmol / mm^2^. A control flowcell with anti-mFc antibody but without immobilized receptor was used as reference. The multiple-cycle kinetic (MCK) or single-cycle kinetic (SCK) method was applied to study molecular interactions. VHH and VHH-NT molecules were prepared by two-fold dilutions in running buffer and injected over the flowcells during 120 s at 30 μl / min with a dissociation time of 300 - 1800 s. Blank runs of running buffer were performed in the same conditions and subtracted from sample runs before evaluation. Double-subtracted sensorgrams were globally fitted with the Langmuir 1:1 binding model from Biacore T200 Evaluation version 3.2. Data are representative of at least 3 independent experiments for VHH C5 and B8 on mouse, human and rhesus monkey TfR1 and for VHH-NT variants on hTfR1. Data for VHH and VHH-NT variants on rhTfR1 were obtained from at least one SPR experiment.

#### Anti-hFc ELISA assay

An anti-hFc ELISA assay was used to assess VHH-hFc fusion concentrations in mouse tissues. The day before the assay, microtitration plates (Nunc MaxiSorp^TM^, Thermo Fisher Scientific) were coated with a mouse anti-hFc antibody at 10 µg / ml (Jackson ImmunoResearch) in NaHCO_3_ 0.1 M buffer and incubated overnight. The day of the assay, the plate was blocked for 1 h at RT under shaking with Blocker Casein 1X in PBS (Thermo Fisher Scientific). After 6 washes with PBS/Tween 20 0.05 % (PBST), the samples were incubated for 2 h at RT under shaking. After 6 washes with PBST, an HRP-conjugated mouse anti-hFc detection antibody 1 / 5000 (Jackson Immunoresearch) was incubated for 1 h at RT under shaking. After 6 PBST washes, revelation of the plate was performed using 50 µl SureBlue™ TMB 1-Component Microwell Peroxidase Substrate (SeraCare, Milford, USA) and stopped with 50 µl H_2_SO4 2N after 15 min incubation. Plates were read at 450 nm with a SpectraMax M2e microplate reader (Molecular Device, San Jose, USA) and data analyzed using SoftMaxPro software version 6.1.

#### *In vivo* plasma and brain pharmacokinetics of Fc fusion proteins

WT C57BL/6 male mice aged 8 to 10 weeks were used for pharmacokinetic/brain uptake studies. Mice were injected subcutaneously with 250 μl of Fc fusions at 35 nmol / kg. At 2, 6, 18 and 42 h post-injection, mice were deeply anesthetized with a cocktail of 100 mg / kg ketamine and 10 mg / kg xylazine (Bayer, Berlin, Germany). Blood was collected in heparinized tubes (Sigma-Aldrich) and plasma was isolated after 10 min centrifugation at 1500 g, 4 °C. Mice were transcardially perfused with 50 ml of NaCl 0.9 %. Brains were extracted, weighed, and lysed in 4-times weigh / volume of PBS / triton X-100 0.1 % containing a protease inhibitor cocktail (Sigma-Aldrich) and frozen at −80 °C before sonication 3 × 10 s. Fc concentrations in the tissues and plasma were measured with the in-house anti-Fc ELISA assay. Results are expressed as VHH-Fc concentration in tissues. All data were expressed as mean ± SEM and p values were assessed by ordinary one-way ANOVA, with Tukey’s multiple comparison test. Results were considered statistically significant at p ≤ 0.05, p ≤ 0.01, p ≤ 0.001 or p ≤ 0.0001; n = 4 mice per molecule and time point.

#### Evaluation of NT(8-13) induced HT in mice

WT C57BL/6 or B-hTfR mice were used to evaluate the potential of low or high molecular weight NT(8-13) fusions to induce HT upon systemic administration. For low molecular weight fusions, mice were injected intravenously with 150 μl vehicle (PBS 1X) or NT(8-13) fusions at 500 nmol / kg. Mouse rectal temperature was measured every 15 min up to 60 min, then every 30 min up to 180 min post-injection using a lubricated thermometer. For high molecular weight fusions, mice were injected intravenously with 150 μl of NT(8-13) fusions at 35 or 10 nmol / kg. Mouse rectal temperature was assessed every 15 min up to 90 min, then every 30 min up to 300 min post-injection. Maximal HT and HT duration were calculated as follows: Maximal HT = temperature prior to the injection (basal temperature) minus the temperature at the peak of the HT. Hypothermia duration = time during which the mouse body temperature is below basal temperature minus 2°C. Areas under the curve (AUC), were determined with GraphPad Prism software 10.2.0. Data were expressed as mean ± SEM and p values were assessed by ordinary one-way ANOVA with Tukeys’ or Dunnetts’ multiple comparison test and represented the mean (± SEM) value of n = 4/ 23 mice. Results were considered statistically significant at p ≤ 0.05, p ≤ 0.01, or p ≤ 0.001.

#### Bioinformatic evaluation of the homology of targeted regions

The structure of the hTfR1 ectodomain was retrieved from the Research Collaboratory for Structural Bioinformatics Protein Data Bank (RCSB PDB) under the reference 1SUV (Cheng, Zak et al. 2004). The structures of the mTfR1 and rhTfR1 ectodomains were predicted using AlphaFold3 (Abramson, Adler et al. 2024), and the best models according to predicted local distance difference test score (pLDDTs) were selected. Root-Mean-Square Deviation (RMSD) matrices were determined by comparing the structures of different ectodomains or regions using the Alignment/align plugin of PyMOL 3.1.3 with default parameters (Yuan, Chan et al. 2017). Percentages of similarity or identity were determined using the Ident and Sim tool available on Sequence Manipulation Suite platform with default parameters (Stothard 2000).

## Acknowledgements

Financial support was provided by the French National Agency for Research (ANR) (NANOVECTOR ANR-15-CE18-0010-03 project coordinated by MK), by the CNRS, Aix Marseille University and Vect-Horus.

## Supplemental Data

**Supplemental Figure S1.**
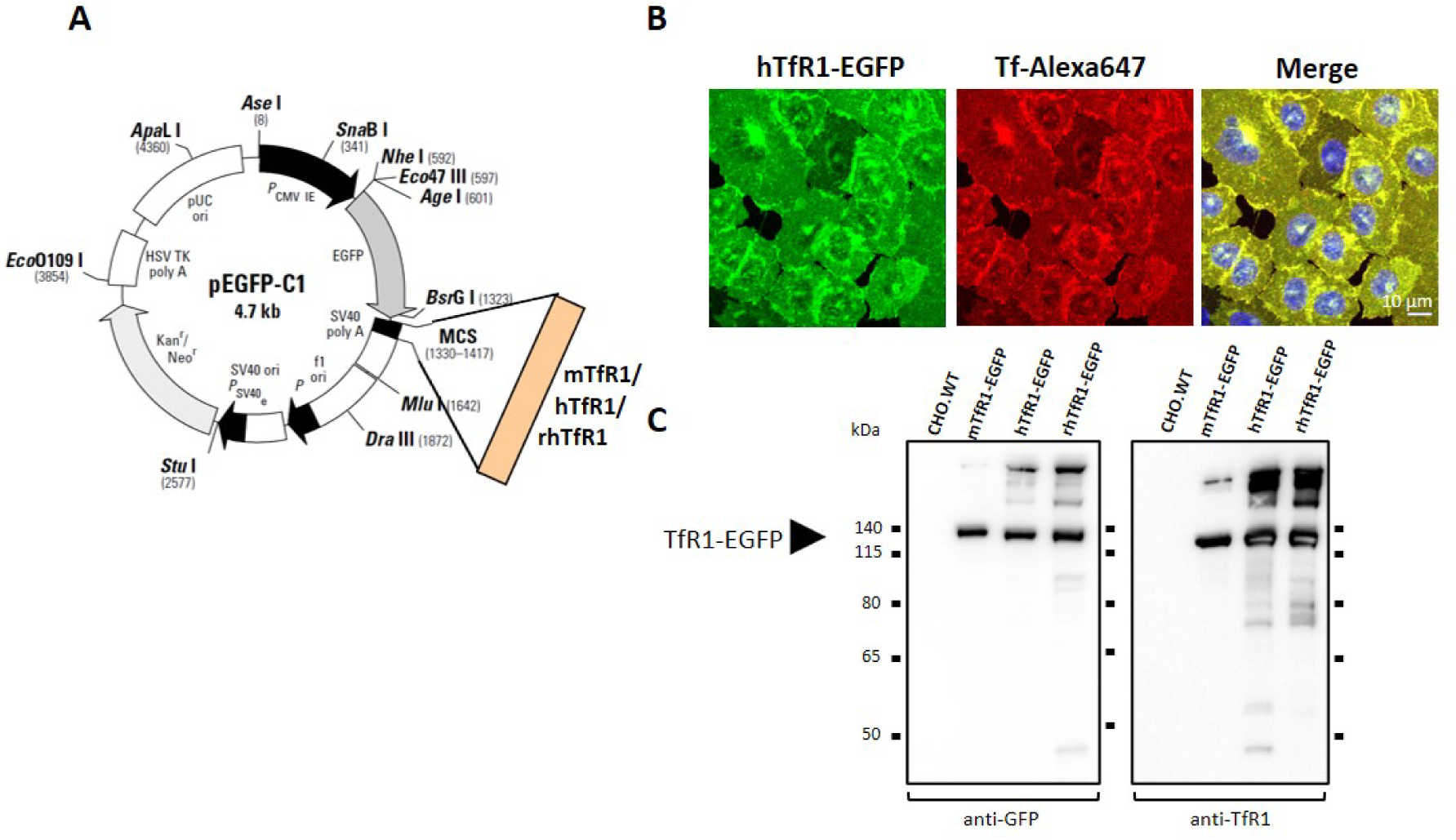
Validation of CHO cell lines expressing the mouse, human and rhesus monkey TfR1 receptor. (A) Map of the plasmid construct used to generate the CHO-hTfR1-EGFP cell line. (B) Representative confocal photomicrographs of CHO-hTfR1-EGFP cells (green) incubated 1 h at 37 °C with Tf-Alexa647 (250 µg/ml). Cell nuclei were labeled with Hoechst#33342 at 0.5 μg/ml (blue). Co-labeling appears in yellow in the merged picture. (C) Western blots performed on cell membrane preparations of CHO cells expressing mTfR1-EGFP, hTfR1-EGFP and rhTfR1-EGFP compared to CHO WT, using a mouse anti-GFP antibody (1/1000) or a rabbit anti-TfR1 antibody (1/1000), followed by HRP-conjugated anti-mouse or anti-rabbit secondary antibodies (1/10000). Arrow indicates the size of the TfR1-GFP monomer.

**Supplemental Figure S2.**
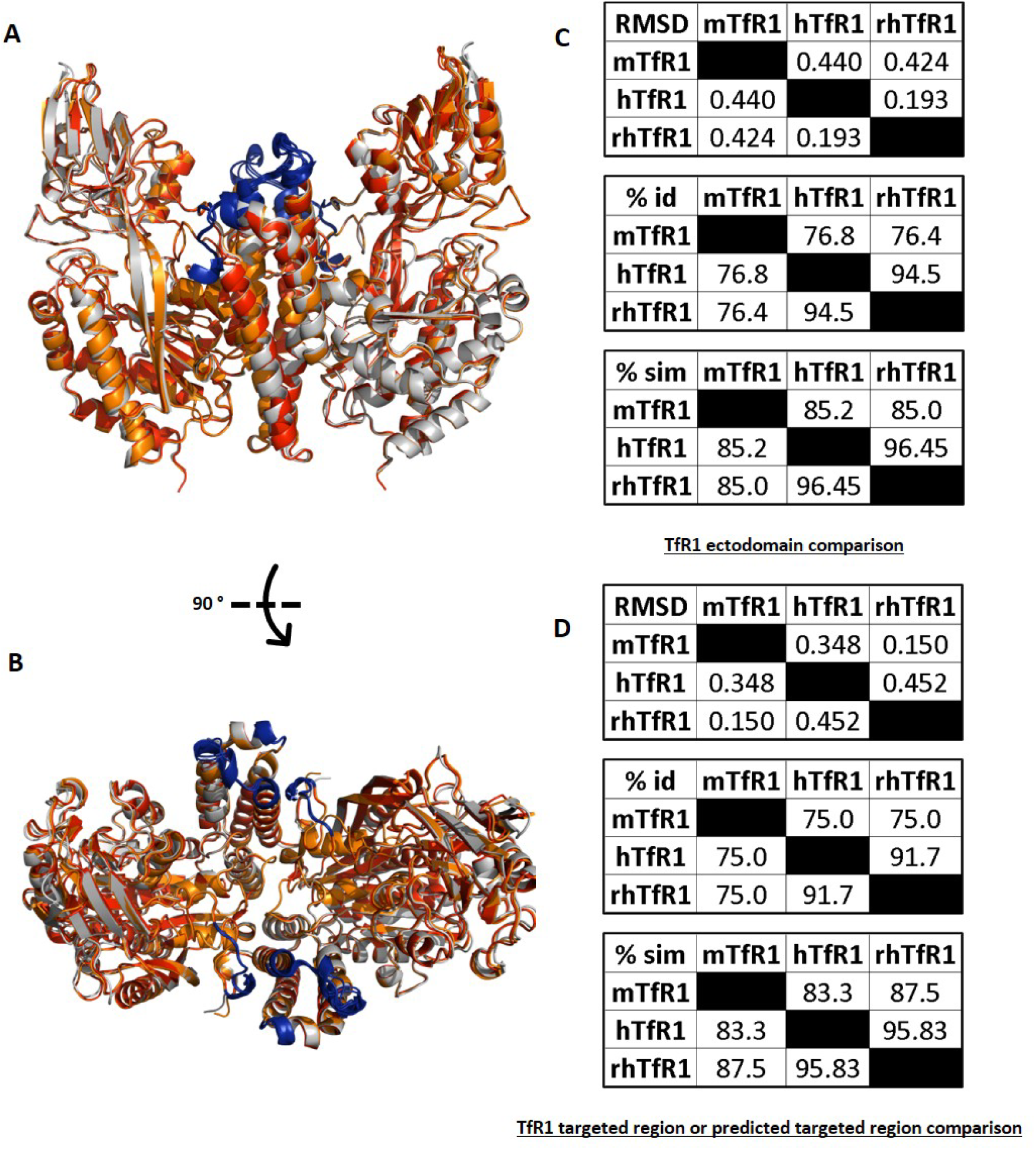
Structural comparison of mouse, human and rhesus monkey TfR1 predicted VHH targeted regions. (A) Side view and (B) top view representations of 3D structures predicted by AlphaFold3 alignments of mTfR1 (gray), rhTfR1 (orange), and hTfR1 (red) ectodomains and epitopes (blue). (C) Root Mean Square Deviation (RMSD), percentage of identity (% id) and percentage of similitude (% sim) matrixes of mTfR1, rhTfR1 and hTfR1 ectodomains. (D) Root Mean Square Deviation (RMSD), percentage of identity (% id) and percentage of similitude (% sim) matrixes of mTfR1, hTfR1 and rhTfR1 predicted targeted regions. RMSD calculations were performed using alignment/align plugin of PyMOL Molecular Graphics System version 3.1.3 (Schrödinger, LLC) and percentage of identity or similitude were determined by Sequence Manipulation Suite.

**Supplemental Figure S3.**
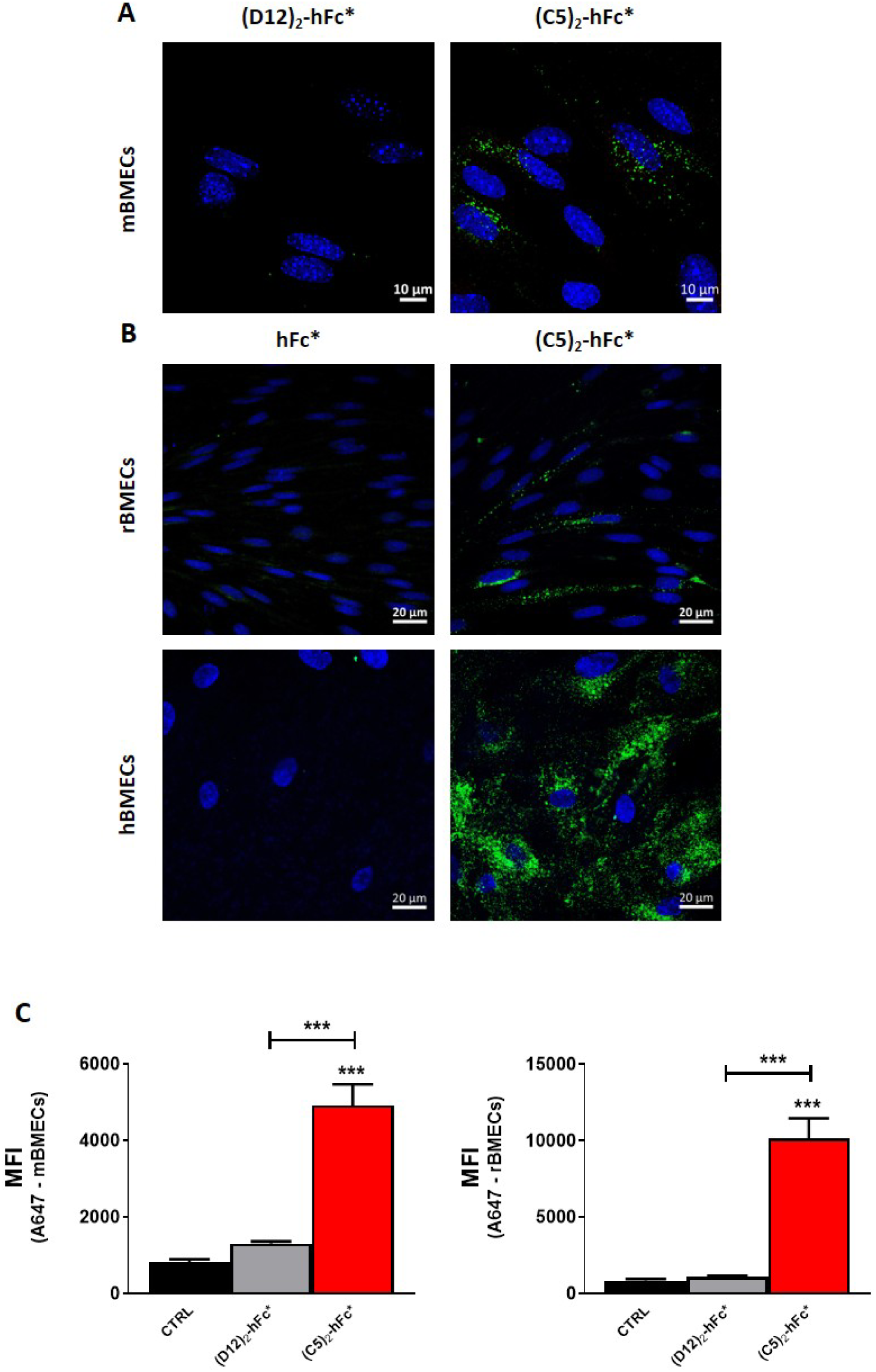
Binding and uptake of (C5)_2_-hFc* fusion by primary mouse, rat and human brain microvascular endothelial cells (BMECs) (A-B) Representative confocal photomicrographs of mouse BMEC monolayers incubated 2 h at 37 °C with (D12)_2_-hFc* or (C5)_2_-hFc* fusions at 500 nM (A) or rat and human BMEC monolayers incubated 24 h at 37 °C with hFc* or (C5)_2_-hFc* fusion at 300 nM (B). Following incubation, BMEC monolayers were fixed with 4 % PFA, permeabilized with 0.1 % saponine, stained with an Alexa488-conjugated anti-hFc antibody (green) and cell nuclei were labeled with Hoechst#33342 at 0.5 µg / ml (blue). (C) Flow cytometry analyses of mouse and rat BMEC monolayers untreated (CTRL) or incubated 2 h at 37 °C with (D12)_2_-hFc* or (C5)_2_-hFc* fusions at 500 nM. Following incubation, BMECs were enzymatically and mechanically dissociated, fixed with 4 % PFA, permeabilized with 0.1 % saponine and then stained with an Alexa647-conjugated anti-hFc antibody. The bar graphs display mean fluorescence intensity (MFI) results. Values reported are the mean (± SEM) of 3 independent experiments with a minimum of 2 technical replicates per experiment. Differences between groups were analyzed using one-way ANOVA with Tukey’s post-hoc test. Only significant differences are shown with *p < 0.05, **p < 0.01 and ***p < 0.001.

**Supplemental Figure S4.**
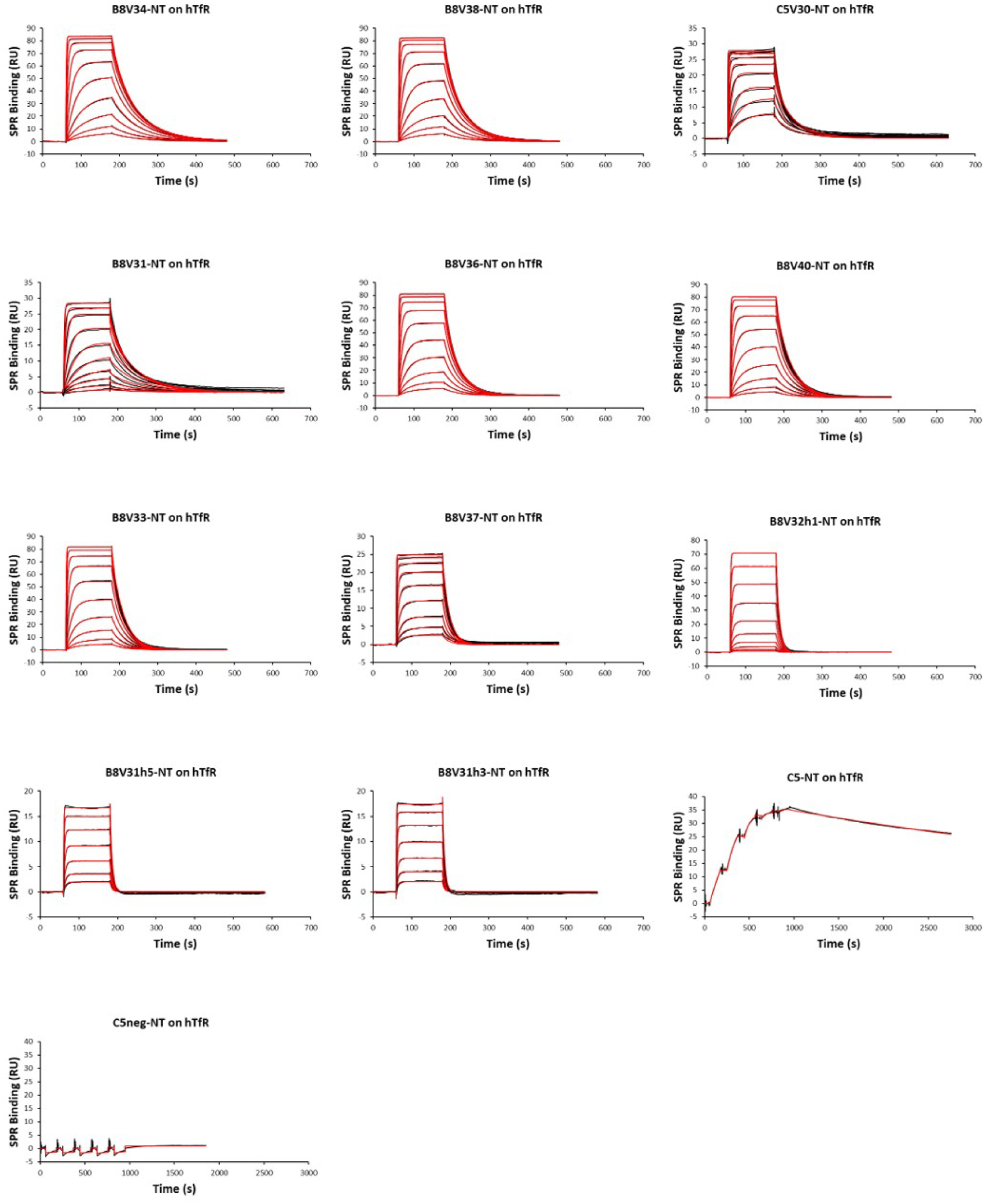
Characterization of VHH-NT(8-13) binding to hTfR by SPR. Increasing concentrations of VHH-NT(8-13) variants (2-2560 nM excepted for C5-NT and C5neg-NT: 2.5 - 40 nM) were injected over immobilized mFc-hTfR using MCK or SCK protocol. The red traces correspond to a global fit of the experimental data (black traces) with a 1:1 interaction model.

**Supplemental Figure S5.**
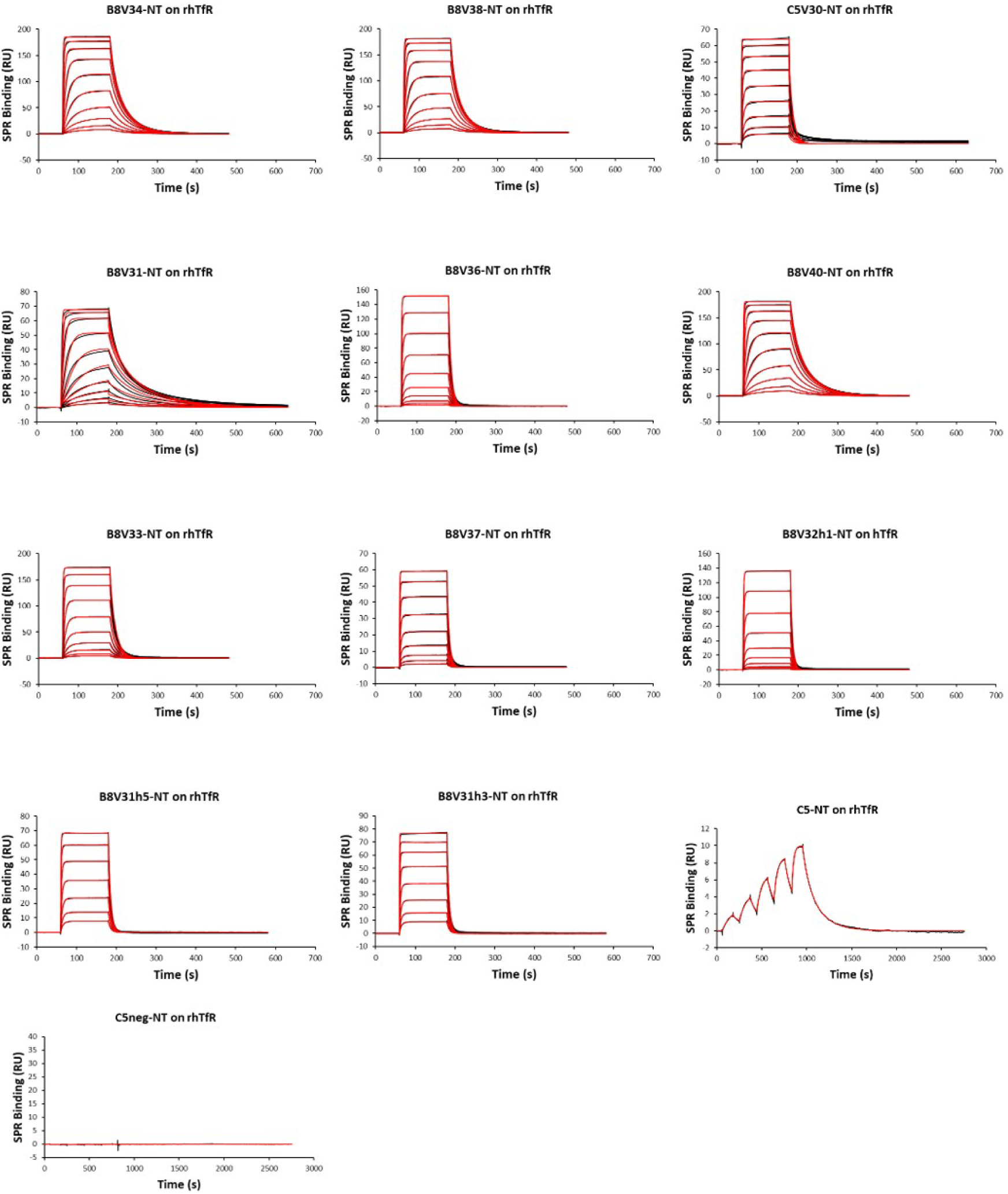
Characterization of VHH-NT(8-13) binding to rhTfR by SPR. Increasing concentrations of VHH-NT(8-13) variants (2-2560 nM excepted for C5-NT and C5neg-NT: 2.5 - 40 nM) were injected over immobilized mFc-rhTfR using MCK or SCK protocol. The red traces correspond to a global fit of the experimental data (black traces) with a 1:1 interaction model.

**Supplemental Table S1.**
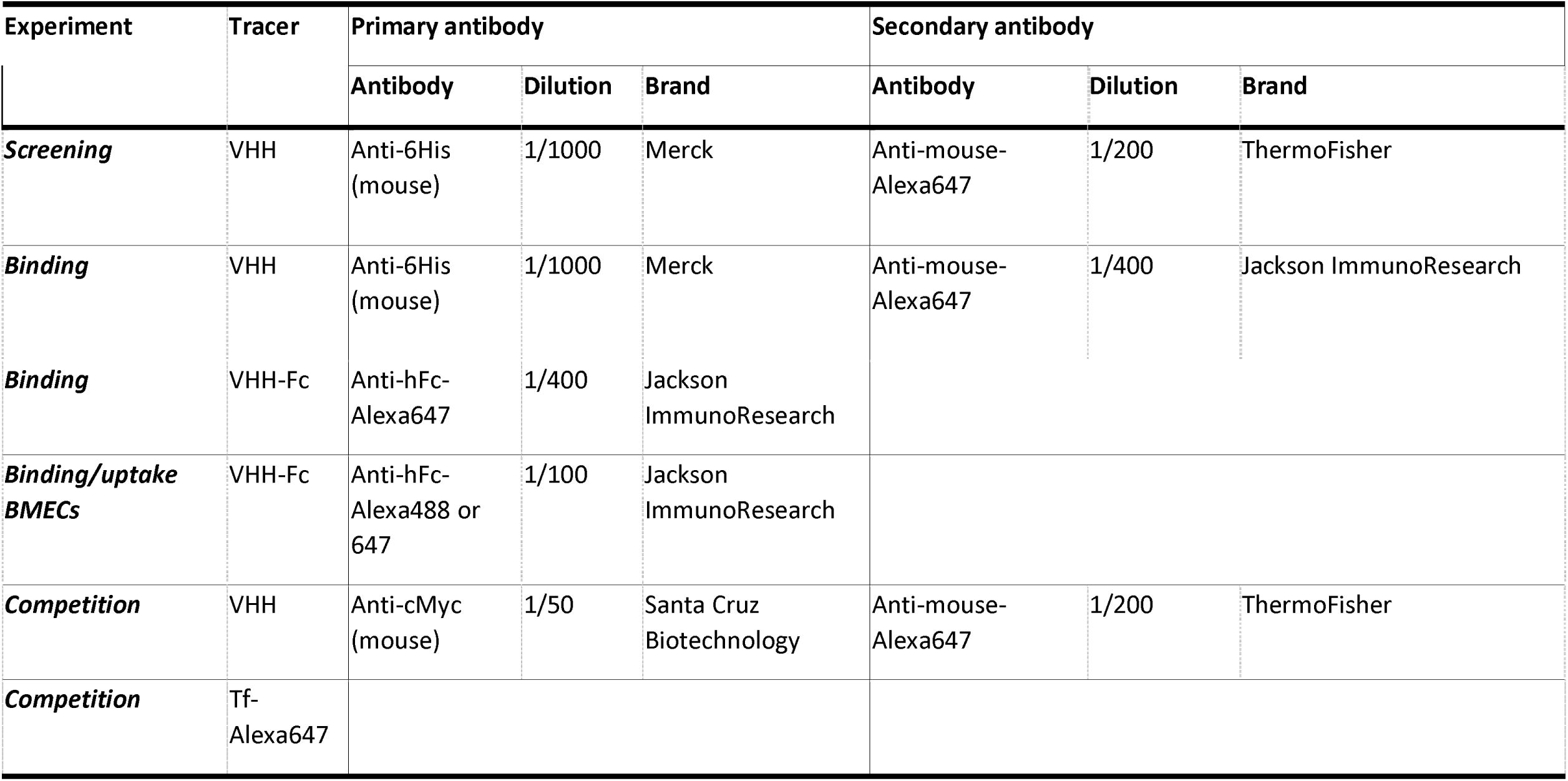
Antibodies used in flow cytometry experiments.

**Supplemental Table S2:**
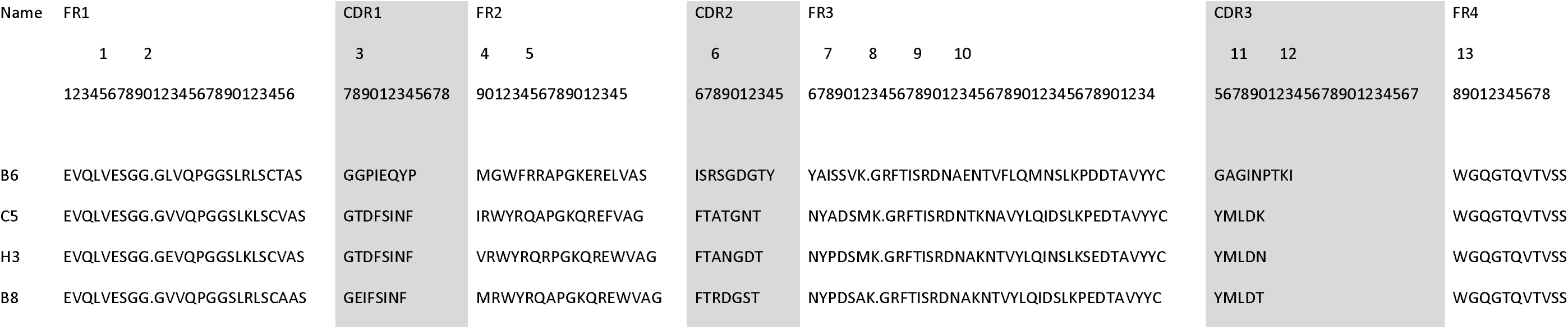
Selected VHH sequences (IMGT numbering)

**Supplemental Table S3:** K_i/app_ on mNTSR1 of VHH-NT affinity variants.

| VHH name | $K_{i/app}$ |
| --- | --- |
| C5neg | NB |
| C5neg-NT(8-13) | $6.7 \pm (3.3)$ |
| C5-NT(8-13) | $7.5 \pm (4)$ |
| B8V34 | NB |
| B8V34-NT(8-13) | $5.5 \pm (4.7)$ |
| B8V38 | NB |
| B8V38-NT(8-13) | $6.1 \pm (3)$ |
| C5V30 | NB |
| C5V30-NT(8-13) | $5 \pm (1.2)$ |
| B8V31 | NB |
| B8V31-NT(8-13) | $4.9 \pm (1.4)$ |
| B8V36 | NB |
| B8V36-NT(8-13) | $4.1 \pm (2.1)$ |
| B8V40 | NB |
| B8V40-NT(8-13) | $6.4 (4.9)$ |
| B8V33 | NB |
| B8V33-NT(8-13) | $9.1 \pm (2.8)$ |
| B8V37 | NB |
| B8V37-NT(8-13) | $4.8 \pm (2)$ |
| B8V32h1 | NB |
| B8V32h1-NT(8-13) | $4 \pm (2.3)$ |
| B8V31h5 | NB |
| B8V31h5-NT(8-13) | $2.7 \pm (0.3)$ |
| B8V31h3 | NB |
| B8V31h3-NT(8-13) | $4 \pm (1.8)$ |

