## Supplemental Data for "Novel VHH targeting a unique TfR1 epitope for efficient cross-species delivery of drugs in the CNS"

**Supplemental Figure S5. Characterization of VHH-NT(8-13) binding to rhTfR by SPR**

Increasing concentrations of VHH-NT(8-13) variants (2-2560 nM excepted for C5-NT and C5neg-NT: 2.5 - 40 nM) were injected over immobilized mFc-rhTfR using MCK or SCK protocol. The red traces correspond to a global fit of the experimental data (black traces) with a 1:1 interaction model.

**Supplemental Table S1. Antibodies used in flow cytometry experiments**

| **Experiment** | **Tracer** | **Primary antibody** | | | **Secondary antibody** | | |
| --- | --- | --- | --- | --- | --- | --- | --- |
|  |  | **Antibody** | **Dilution** | **Brand** | **Antibody** | **Dilution** | **Brand** |
| ***Screening*** | VHH | Anti-6His (mouse) | 1/1000 | Merck | Anti-mouse-Alexa647 | 1/200 | ThermoFisher |
| ***Binding*** | VHH | Anti-6His (mouse) | 1/1000 | Merck | Anti-mouse-Alexa647 | 1/400 | Jackson ImmunoResearch |
| ***Binding*** | VHH-Fc | Anti-hFc-Alexa647 | 1/400 | Jackson ImmunoResearch |  | | |
| ***Binding/uptake BMECs*** | VHH-Fc | Anti-hFc-Alexa488 or 647 | 1/100 | Jackson ImmunoResearch |  | | |
| ***Competition*** | VHH | Anti-cMyc (mouse) | 1/50 | Santa Cruz Biotechnology | Anti-mouse-Alexa647 | 1/200 | ThermoFisher |
| ***Competition*** | Tf-Alexa647 |  | | |  | | |

**Supplemental Table S2: Selected VHH sequences (IMGT numbering)**

| Name | FR1  1 2  12345678901234567890123456 | CDR1  3  789012345678 | FR2  4 5  90123456789012345 | CDR2  6  6789012345 | FR3  7 8 9 10  678901234567890123456789012345678901234 | CDR3  11 12  56789012345678901234567 | FR4  13  89012345678 |
| --- | --- | --- | --- | --- | --- | --- | --- |
| B6 | EVQLVESGG.GLVQPGGSLRLSCTAS | GGPIEQYP | MGWFRRAPGKERELVAS | ISRSGDGTY | YAISSVK.GRFTISRDNAENTVFLQMNSLKPDDTAVYYC | GAGINPTKI | WGQGTQVTVSS |
| C5 | EVQLVESGG.GVVQPGGSLKLSCVAS | GTDFSINF | IRWYRQAPGKQREFVAG | FTATGNT | NYADSMK.GRFTISRDNTKNAVYLQIDSLKPEDTAVYYC | YMLDK | WGQGTQVTVSS |
| H3 | EVQLVESGG.GEVQPGGSLKLSCVAS | GTDFSINF | VRWYRQRPGKQREWVAG | FTANGDT | NYPDSMK.GRFTISRDNAKNTVYLQINSLKSEDTAVYYC | YMLDN | WGQGTQVTVSS |
| B8 | EVQLVESGG.GVVQPGGSLRLSCAAS | GEIFSINF | MRWYRQAPGKQREWVAG | FTRDGST | NYPDSAK.GRFTISRDNAKNTVYLQIDSLKPEDTAVYYC | YMLDT | WGQGTQVTVSS |

**Supplemental Table S3: K_i/app_ on mNTSR1 of VHH-NT affinity variants**

| VHH name | K_i/app_ |
| --- | --- |
| C5neg | NB |
| C5neg-NT(8-13) | 6.7 ± (3.3) |
| C5-NT(8-13) | 7.5 ± (4) |
| B8V34 | NB |
| B8V34-NT(8-13) | 5.5 ± (4.7) |
| B8V38 | NB |
| B8V38-NT(8-13) | 6.1 ± (3) |
| C5V30 | NB |
| C5V30-NT(8-13) | 5± (1.2) |
| B8V31 | NB |
| B8V31-NT(8-13) | 4.9± (1.4) |
| B8V36 | NB |
| B8V36-NT(8-13) | 4.1 ± (2.1) |
| B8V40 | NB |
| B8V40-NT(8-13) | 6.4 (4.9) |
| B8V33 | NB |
| B8V33-NT(8-13) | 9.1± (2.8) |
| B8V37 | NB |
| B8V37-NT(8-13) | 4.8 ± (2) |
| B8V32h1 | NB |
| B8V32h1-NT(8-13) | 4 ± (2.3) |
| B8V31h5 | NB |
| B8V31h5-NT(8-13) | 2.7 ± (0.3) |
| B8V31h3 | NB |
| B8V31h3-NT(8-13) | 4 ± (1.8) |

**Supplemental Materials and Methods**

**Preparation of cell membrane extracts**

To assess the expression of the different TfR1 constructs in stable cell lines, cells were washed with PBS, scrapped, centrifuged and treated as recommended in the ProteoExtract® Subcellular Proteome Extraction Kit (Calbiochem, San Diego, USA) to prepare total cell membranes. Protein contents in membranes were quantified using the DC™ Protein Assay (Bio-Rad, Hercules, USA) following the manufacturer's instructions.

**Western blot assay**

Membrane proteins (1 µg) from the CHO cell lines were separated by SDS-PAGE on 4-12 % Tris-bis plus gel (Thermo Fisher Scientific) and transferred onto nitrocellulose membranes (Amersham Biosciences, Buckinghamshire, UK) using the iBlot™ 2 Dry Blotting System (Thermo Fisher Scientific). Nitrocellulose membranes were blocked in TBS/Tween-20 0.1 % (TBS-T)/skimmed milk 5 % and probed overnight at 4 °C with rabbit anti-TfR1 (Genetex, Irvine, USA) or with mouse anti-GFP (Sigma-Aldrich) primary antibodies diluted 1/1000 in blocking buffer. After several washes in TBS-T, membranes were incubated for 1 h at RT with horseradish peroxidase (HRP)-conjugated anti-rabbit or anti-mouse secondary antibodies (Jackson ImmunoResearch) diluted 1/10 000 in blocking buffer. After several washes, proteins were detected using chemiluminescence (ECL) immunoblotting detection reagents (GE Healthcare) and revealed with the G:Box chemi XX6 system (Syngene, Cambridge, UK).

**Immunocytochemistry**

After 30 min cell saturation with cell culture medium supplemented with low endotoxin BSA 1 %, CHO-hTfR1-EGFP cells were incubated 1 h at 37 °C with Alexa647-conjugated transferrin (Tf-Alexa647, Thermo Fisher Scientific) at 250 µg / ml and were washed and fixed with 4 % PFA for 10 min at RT, washed again and processed for immunocytochemistry. Cells were incubated with Hoechst#33258 at 0.5 µg / ml for nuclei staining (Thermo Fisher Scientific). After 3 washes with PBS, coverslips were mounted in ProLong® Gold Antifade reagent (Thermo Fisher Scientific) and were analyzed using a LSM700 (Zeiss) confocal microscope with Zen 2012 software. Images were obtained using a 63× Plan-Apochromate oil immersion objective.

**Binding and uptake assays on endothelial cells**

**Brain microvascular endothelial cells (BMECs) production**

Rat and mice primary BMECs were produced as previously described (Molino, Jabes et al. 2014). Briefly, brain microvessels were isolated from cerebral cortices of 5-week-old male Wistar rats and C57BL/6 mice (Janvier Labs, France). Brains were dissected, meninges removed, and cerebral cortices harvested. Then, cortices were mechanically dissociated using Dounce homogenizers and tissue homogenates centrifugated (1000 g, 5 min, 4 °C). Pellets were subjected to a first enzymatic digestion (Liberase DH, Merck) 30 min at 37 °C followed by a centrifugation (3600 g, 15 min at 4 °C) in a solution containing 25 % bovine serum albumin (BSA, Interchim, Montluçon, France) in HBSS (Thermo Fisher Scientific) to separate vessel fragments from nervous tissue. The vessel fragments underwent a second enzymatic digestion with the same enzymatic solution 30 min (mice) or 60 min (rat) at 37 °C under agitation to free endothelial cells from their extracellular matrix. After centrifugation (1000 g, 5 min, 4 °C), residual small vessel fragments and dissociated cells were seeded in petri dishes. Primary BMECs were cultured and amplified at 37 °C in 5 % CO_2_ in DMEM/F12 Glutamax medium (Thermo Fisher Scientific) supplemented with 15 % platelet poor plasma derived bovine serum (First Link, UK), 100 µg / ml heparin (Sigma-Aldrich), 50 µg / ml gentamicin, 5 mM HEPES and 2 ng / ml bFGF (all from Thermo Fisher Scientific). During the first week of culture, remaining contaminating cells such as pericytes were further eliminated using gradually decreasing concentrations of puromycin (from 4 µg / ml to 1 µg / ml, Sigma-Aldrich). After amplification and purification, rat and mouse BMECs were seeded (passage 1) on 12-well ThinCert® inserts (Greiner Bio-One) with polyester (PET) microporous membranes (1 µm) for immunocytochemistry analyses (monoculture) or in 24-well culture plates (Falcon) for flow cytometry evaluations. Both inserts and 24-well were precoated with human collagen type IV (2 µg/cm2, Sigma Aldrich) and fibronectin (2 µg / cm2, Corning). Cells were incubated 3 days to establish BMEC monolayers in the same culture medium with the addition of hydrocortisone 500 nM (Sigma Aldrich) before incubation with VHH-hFc fusions 2 h or 24 h at 37 °C.

Human primary BMECs were produced by Brainplotting™ (Paris, France) in partnership with Sainte-Anne Hospital (Paris, France) from brain tissue harvested during a scheduled tumor resection surgery with written informed consent from the patient (authorization number CODECOH DC-2014-2229). Human brain microvessels were obtained from a surgical resection of a 36-year-old patient with a diffuse oligodendroglial glioma. Microvessels were isolated from healthy peritumoral brain tissue using an enzymatic procedure (Chaves, Campanelli et al. 2020) adapting methods previously published for rats (Perrière, Yousif et al. 2007). Briefly, tissue samples were carefully cleaned from meninges and excess of blood and dissociated using an enzymatic mix. Then, microvessels were isolated by retention on a 10 μm mesh. After seeding brain capillaries in petri dishes, primary BMECs were cultured at 37 °C in 5 % CO_2_, purified using puromycin and amplified in EBM-2 medium (Lonza) supplemented with 20% serum and growth factors (Sigma-Aldrich). After amplification and purification, human BMECs were seeded (passage 1) on 12-well Transwell™ inserts (Corning, Glendale, Arizona, USA) with polyester (PET) microporous membranes (0.4 µm) and incubated with VHH-hFc fusions 24 h at 37 °C.

**Immunocytochemistry and confocal microscopy**

Following incubation, inserts with human and rat BMEC monolayers were washed thrice with D-PBS and fixed in 4 % PFA for 15 min. Microporous membranes were gently dissociated from the plastic inserts with a razor blade before immunocytochemistry. Cells were then blocked and permeabilized for 30 min at RT in blocking buffer that contained 3 % BSA and 0.1 % saponin (Sigma Aldrich). Subsequently, cells were incubated for 1 h at RT with an Alexa488-conjugated anti-hFc antibody (Supplemental Table S1) diluted in PBS with 1 % BSA and 0.1 % saponin and Hoechst 33342 (0.5 μg / ml Sigma-Aldrich). Cells were washed thrice with PBS, and membranes were mounted in Prolong Gold antifade mounting medium (Thermo Fisher Scientific). Images were acquired and processed using a confocal microscope (LSM 700) and Zen software (Carl Zeiss).

**Flow cytometry**

Following incubation, rat and mouse BMEC monolayers were washed twice in D-PBS supplemented with 5 mM EDTA and then harvested using prewarmed trypsin-EDTA solution (all from Thermo Fisher Scientific) during 5 min at 37 °C and centrifugation (300 g, 5 min, 4 °C). Cells were fixed with D-PBS / EDTA 5 mM / PFA 2% solution for 15 min at room temperature. After washing in D-PBS / EDTA 5 mM solution and centrifugation (300 g, 5 min, 4 °C), cells were then blocked and permeabilized for 15 min at RT in solution containing D-PBS / 5 mM EDTA / 0.1 % saponin / 1 % BSA. Subsequently, cells were incubated for 1 h at RT in the same solution containing an Alexa647-conjugated anti-hFc antibody (Supplemental Table S1) and Hoechst 33342 (0.5 μg / ml Sigma-Aldrich). After incubation, cells were washed in D-PBS / EDTA 5 mM solution, centrifugated (500 g, 5 min, 4 °C) and resuspended in a total volume of 300 μl before analysis. Alexa680-associated fluorescence was quantified with Attune™ NxT flow cytometer equipped with Attune™ Cytometric software v5.2.0 (Thermo Fisher Scientific). Mean fluorescence intensity (MFI) data are presented and refer to the total population analyzed. At least 10,000 events per condition were recorded. Results are representative of at least three independent experiments with a minimum of two technical replicates per experiment.
