## Supplemental Figures for "Novel VHH targeting a unique TfR1 epitope for efficient cross-species delivery of drugs in the CNS"

### Slide 1
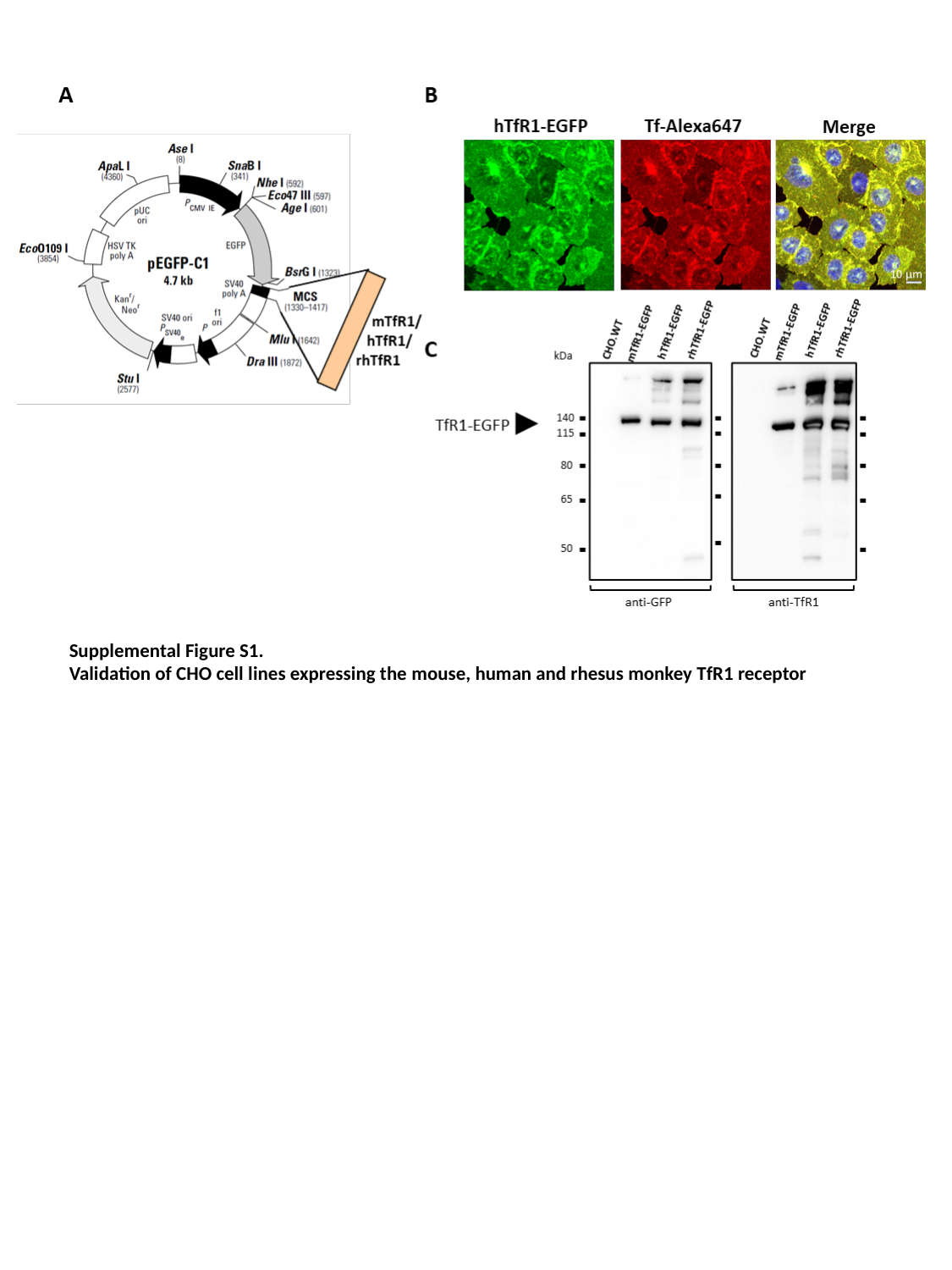

Supplemental Figure S1.
Validation of CHO cell lines expressing the mouse, human and rhesus monkey TfR1 receptor

### Slide 2
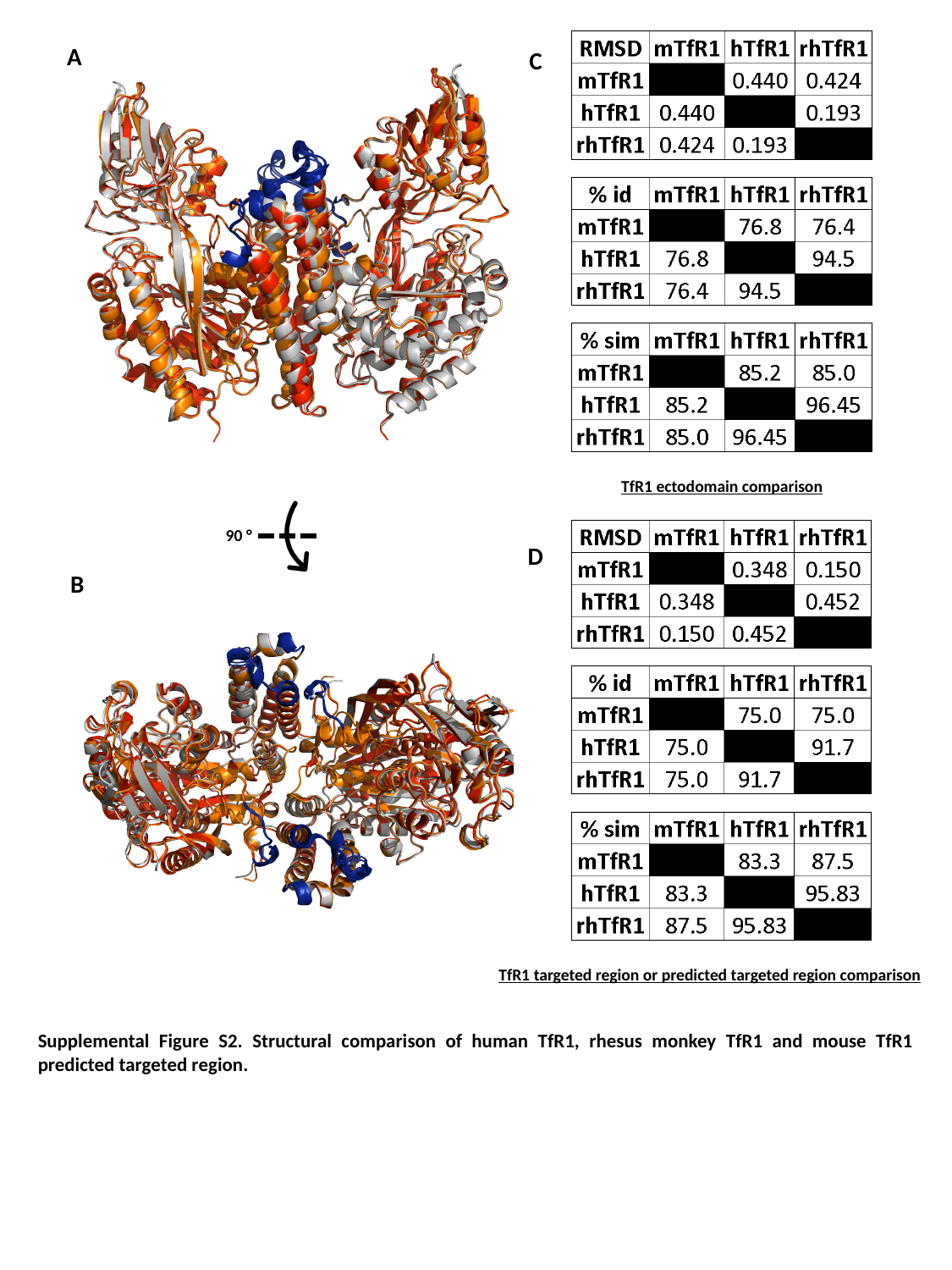

A
C
TfR1 ectodomain comparison
90 °
D
B
TfR1 targeted region or predicted targeted region comparison
Supplemental Figure S2. Structural comparison of human TfR1, rhesus monkey TfR1 and mouse TfR1 predicted targeted region.

### Slide 3
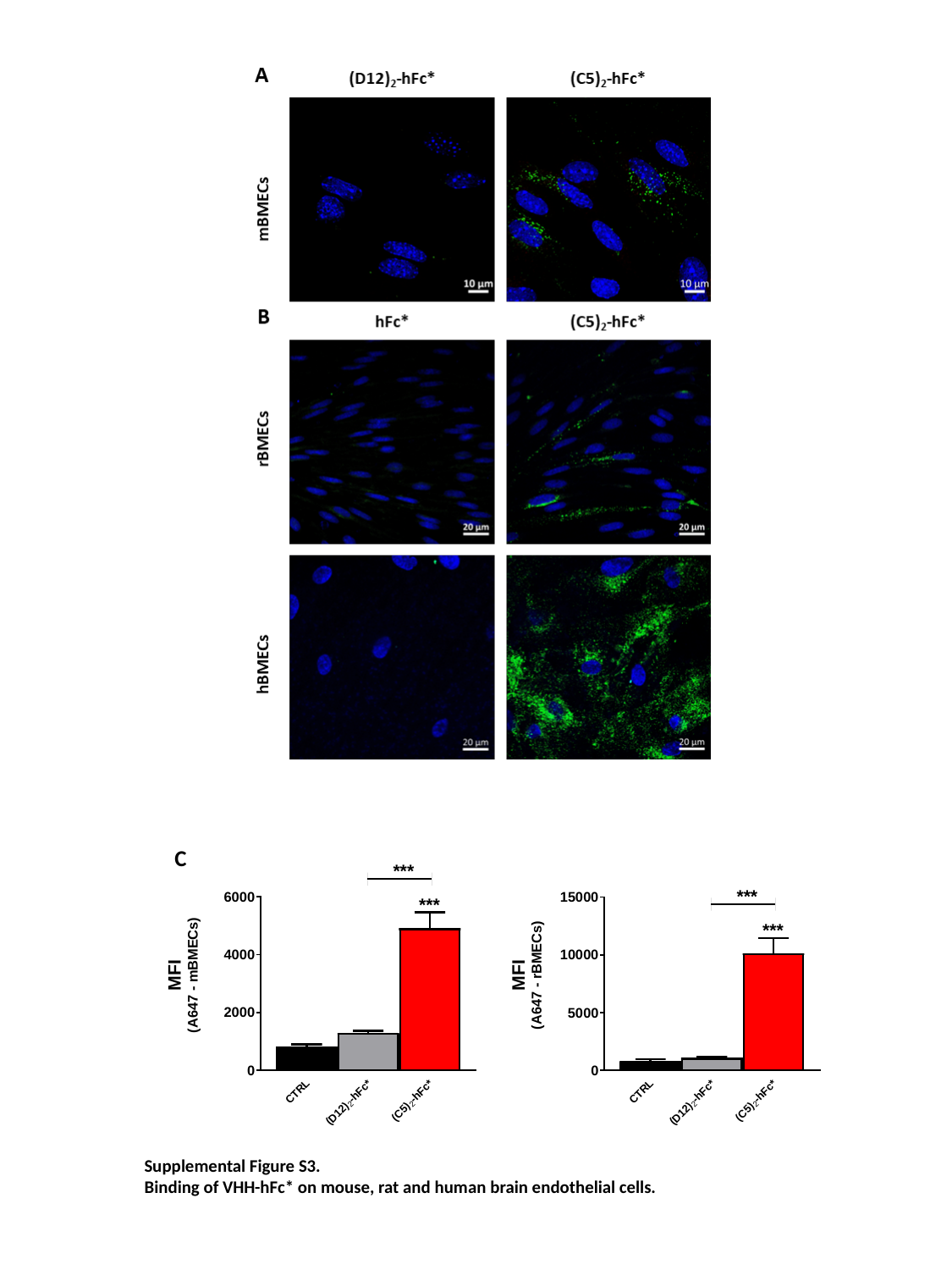

C
Supplemental Figure S3.
Binding of VHH-hFc* on mouse, rat and human brain endothelial cells.

### Slide 4
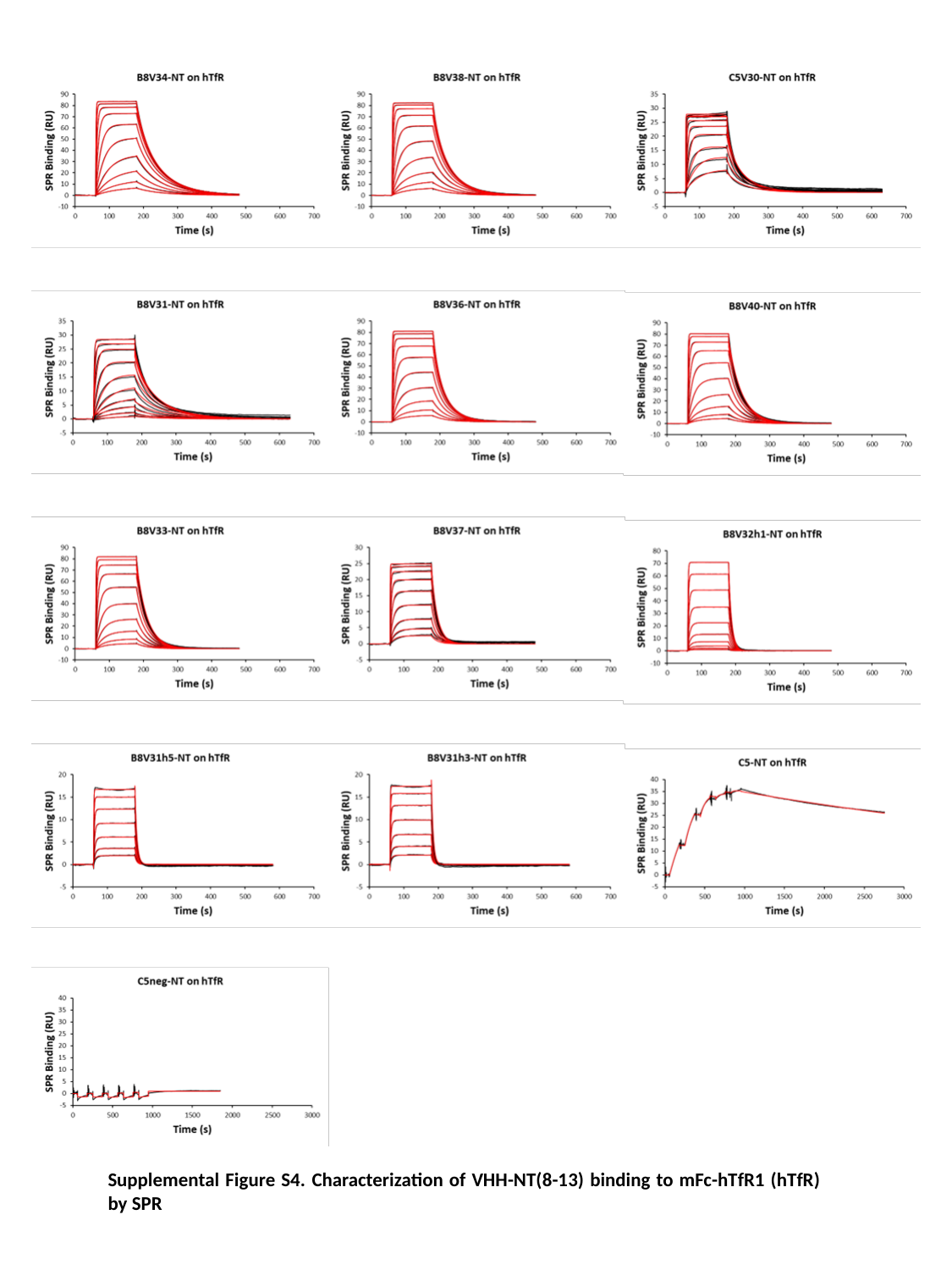

Supplemental Figure S4. Characterization of VHH-NT(8-13) binding to mFc-hTfR1 (hTfR) by SPR

### Slide 5
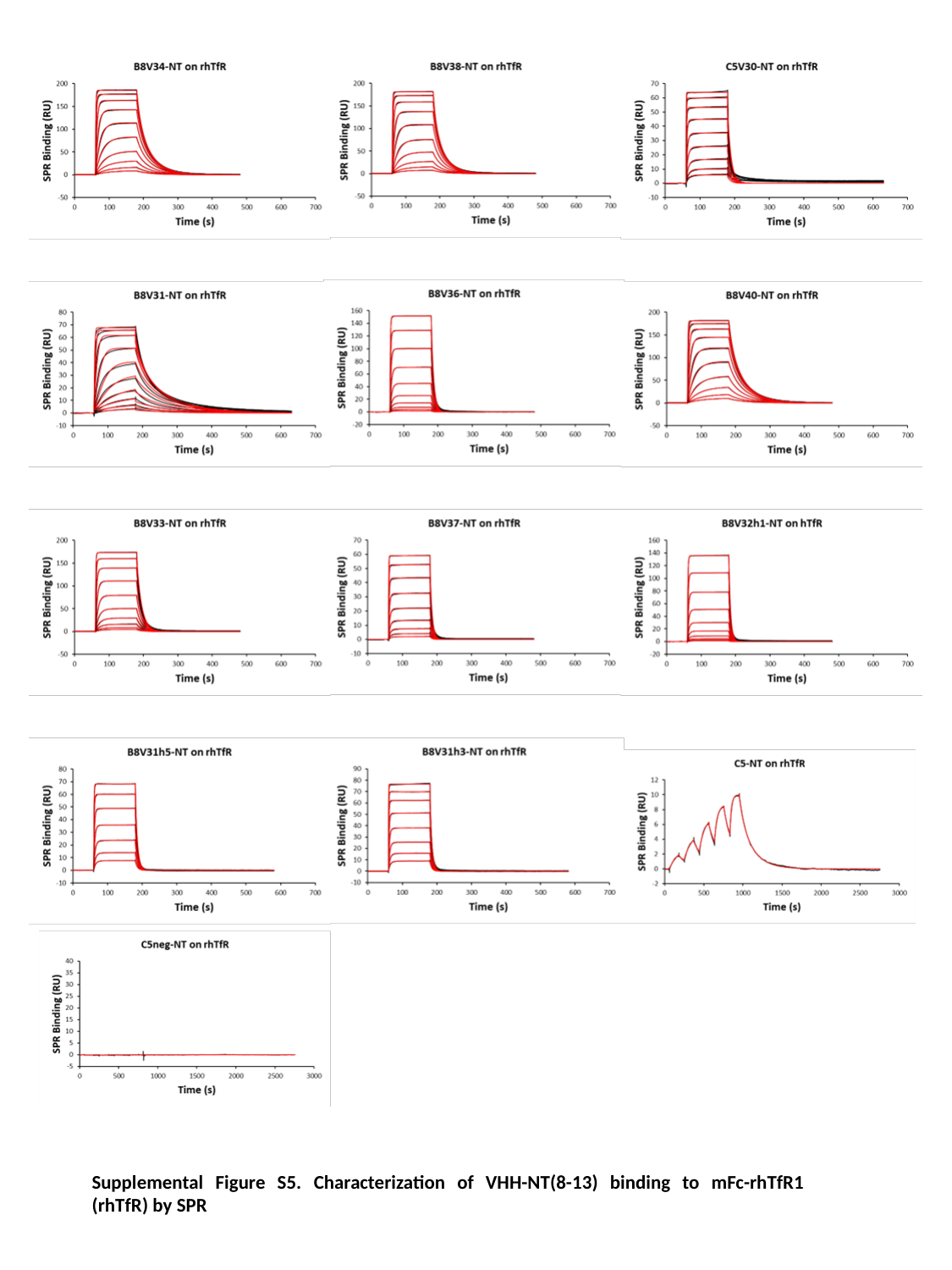

Supplemental Figure S5. Characterization of VHH-NT(8-13) binding to mFc-rhTfR1 (rhTfR) by SPR
